# Stereopy: modeling comparative and spatiotemporal cellular heterogeneity via multi-sample spatial transcriptomics

**DOI:** 10.1101/2023.12.04.569485

**Authors:** Shuangsang Fang, Mengyang Xu, Lei Cao, Xiaobin Liu, Marija Bezulj, Liwei Tan, Zhiyuan Yuan, Yao Li, Tianyi Xia, Longyu Guo, Vladimir Kovacevic, Junhou Hui, Lidong Guo, Chao Liu, Mengnan Cheng, Li’ang Lin, Zhenbin Wen, Bojana Josic, Nikola Milicevic, Ping Qiu, Qin Lu, Yumei Li, Leying Wang, Luni Hu, Chao Zhang, Qiang Kang, Fengzhen Chen, Ziqing Deng, Junhua Li, Mei Li, Shengkang Li, Yi Zhao, Guangyi Fan, Yong Zhang, Ao Chen, Yuxiang Li, Xun Xu

## Abstract

Tracing cellular dynamic changes across conditions, time, and space is crucial for understanding the molecular mechanisms underlying complex biological systems. However, integrating multi-sample data in a unified and flexible way to explore cellular heterogeneity remains a major challenge. Here, we present Stereopy, a flexible and versatile framework for modeling and dissecting comparative and spatiotemporal patterns in multi-sample spatial transcriptomics with interactive data visualization. To optimize this flexible framework, we have developed three key components: a multi-sample tailored data container, a scope controller, and an analysis transformer. Furthermore, Stereopy showcases three transformative applications supported by pivotal algorithms. Firstly, the multi-sample cell community detection (CCD) algorithm introduces an innovative capability to detect specific cell communities and identify genes responsible for pathological changes in comparable datasets. Secondly, the spatially resolved temporal gene pattern inference (TGPI) algorithm represents a notable advancement in detecting important spatiotemporal gene patterns while concurrently considering spatial and temporal features, which enhances the identification of important genes, domains and regulatory factors closely associated with temporal datasets. Finally, the 3D niche-based regulation inference tool, named NicheReg3D, reconstructs the 3D cell niches to enable the inference of cell-gene interaction network within the spatial texture, thus bridging intercellular communications and intracellular regulations to unravel the intricate regulatory mechanisms that govern cellular behavior. Overall, Stereopy serves as both a bioinformatics toolbox and an extensible framework that provides researchers with enhanced data interpretation abilities and new perspectives for mining multi-sample spatial transcriptomics data.

## Introduction

Cells are not static. Dynamical and orderly cellular proliferation, differentiation, and maturation accomplish their functions by spatially interacting with the microenvironment consisting of external stimuli and other cells, which forms the complex architecture of multicellularity. However, understanding the underlying mechanisms that govern disease, development, and homeostasis is still an open question in scientific investigations. Such investigations often require the simultaneous analysis of datasets comprising multiple samples, enabling researchers to effectively track the specificity and variation of cells and genes across different conditions, time points, and spatial dimensions [1, 2]. The advent of high-resolution spatial resolved transcriptomics (SRT) technologies, such as Stereo-seq [3], Slide-seq [4], MERFISH [5], SeqFish [6], STARmap [7], and Xenium [8], holds immense potential for generating large-scale multi-sample datasets. These advancements also underscore the demand for more advanced analytical approaches, enabling the exploration of molecular alterations and characteristics in various contexts—be it conditional, temporal, or spatial [9]. These contexts span a wide spectrum of applications, ranging from tracking disease progression [10, 11] and monitoring temporal cellular development [12] to dissecting the intricacies of spatial organogenesis [3, 13].

Pioneering analysis frameworks for spatial or single-cell transcriptomics data such as Squidpy [14], Giotto [15], Scanpy [16], Seurat [17], and scvi-tools [18] have been widely employed, enabling temporally and/or spatially resolved studies with spatial omics data. However, they were primarily designed for single-sample analysis [19]. Multi-sample analyses also necessitate tailored data containers to enable efficient data organization, flexibility, and scalability. However, existing data containers including AnnData (used in Scanpy [16]), SeuratObject (used in Seurat [17]), GiottoObject (used in Giotto [15]), and MuData (used in Muon [20]) are inherently limited in their capacity to manage multiple samples. Furthermore, existing methods [21-23] lack the capacity for advanced multi-sample analysis, including individual sample analysis, managing analysis step dependencies, and transitioning between single-sample and multi-sample results. With decreasing data costs and growing tissue complexity, efficient methods for storing, integrating, and visualizing multi-sample omics data across various dimensions are urgently needed.

However, developing a multi-sample analysis framework requires solving complex challenges. Key among these is the need to establish a standardized framework for contextualizing analysis modules and visualization functions, design scalable data representation for multi-sample data management, and provide integrated solutions for multi-purpose tasks. In response to these challenges, we propose Stereopy, a comprehensive multi-sample analysis toolkit that includes a complete set of extensible tools for managing, analyzing, and visualizing multi-sample spatial omics data. To efficiently manage multiple samples in a unified and convenient manner, we address the flexible cross-sample storage of input data and analysis results while ensuring the accessibility and traceability of outcomes. A flexible analysis framework is also designed to enable analysis on specific samples, manage dependencies between different analysis steps, and facilitate the transformation of single-sample results into integrated results.

The resulting comprehensive analysis solutions improve the utilization of information when applied to different datasets (Supplementary Note 1). For comparative analysis, Stereopy compares disturbed or disease samples to control samples, analyzes the diversity at both global and local levels in the spatial context, and identifies the changes in functional mechanisms that arise as a result of stress responses or disease perturbations. The multi-sample cell community detection (CCD) algorithm developed in Stereopy introduces an innovative capability to detect variations at the cell community level in comparative samples (Supplementary Note 2.1). For temporal analysis, it explores temporal variations in cell types and gene expression over time, reflecting the temporal and molecular intricacies of organismal development with spatial resolution. The proposed spatial resolved temporal gene pattern inference (TGPI) algorithm in Stereopy presents a significant enhancement in detecting spatiotemporal gene patterns marking the first instance which simultaneously considering spatial and temporal features (Supplementary Note 2.2). For 3D integrated analysis, it provides a powerful tool, named NicheReg3D, for reconstructing the cell niche and investigating the effect of intercellular signaling on the intracellular regulation within spatial constraints, enabling new insights into organ development (Supplementary Note 2.3). Additionally, Stereopy offers flexible data visualization techniques for both 2D and 3D datasets to researchers to conveniently explore the intricate spatiotemporal changes of genes and cells and accurately model underlying biological processes across different dimensions. Stereopy is available at https://github.com/STOmics/Stereopy. Its documentation and extensive tutorials can be explored at https://stereopy.readthedocs.io/en/latest.

## Results

### Overview of Stereopy

Stereopy provides a comprehensive and robust solution for multi-sample analysis, comprising three main components: a multi-sample data container, multi-sample data analysis modules, and a multi-sample interactive visualization (Fig. 1a). This platform is designed to support the closed loop of data management through a multi-sample data (MsData) container, multi-sample scope (MSS) controller, multi-sample analysis transformer and multi-sample data file format. Stereopy also offers well-organized analysis modules and key algorithms for spatial omics data, covering three main multi-sample data analysis scenarios: comparative, spatiotemporal, and 3D integrated analysis (Supplementary Note 1). These analytical capabilities include the identification of specific cell communities and functional modules in comparative multi-sample datasets (Fig. 1b (i) and Supplementary Note 1.1), detection of temporal variable genes and gene patterns in time-series datasets (Fig. 1b (ii) and Supplementary Note 1.2), and inference of complete signaling paths from cell-cell communication to gene regulation networks in 3D datasets (Fig. 1b (iii) and Supplementary Note 1.3). Stereopy’s representative key algorithms for each data type are highlighted: 1) the cell community detection algorithm, which finds common or specific communities between comparative samples, enriching the comparative analysis capability (Fig. 1c (i) and Supplementary Note 2.1); 2) the spatially resolved temporal gene pattern identification method, which explores specific genes and modules related to temporal development under spatial restriction (Fig. 1c (ii) and Supplementary Note 2.2); and 3) 3D regulation mechanism inference, which probes complete gene regulation mechanisms by mining extracellular ligand-receptor interactions, intracellular regulation networks, and signaling pathways between them in the entire 3D tissue level (Fig. 1c (iii) and Supplementary Note 2.3). Furthermore, Stereopy provides 2D and 3D spatial omics interactive visualization [24], which generates high-quality data explorations and supports user-defined browsing. These unique capabilities of Stereopy make it a valuable tool for researchers to analyze and interpret multi-sample SRT data, with powerful functionalities that enables a deeper understanding of biological processes and mechanisms.

**Fig. 1.**
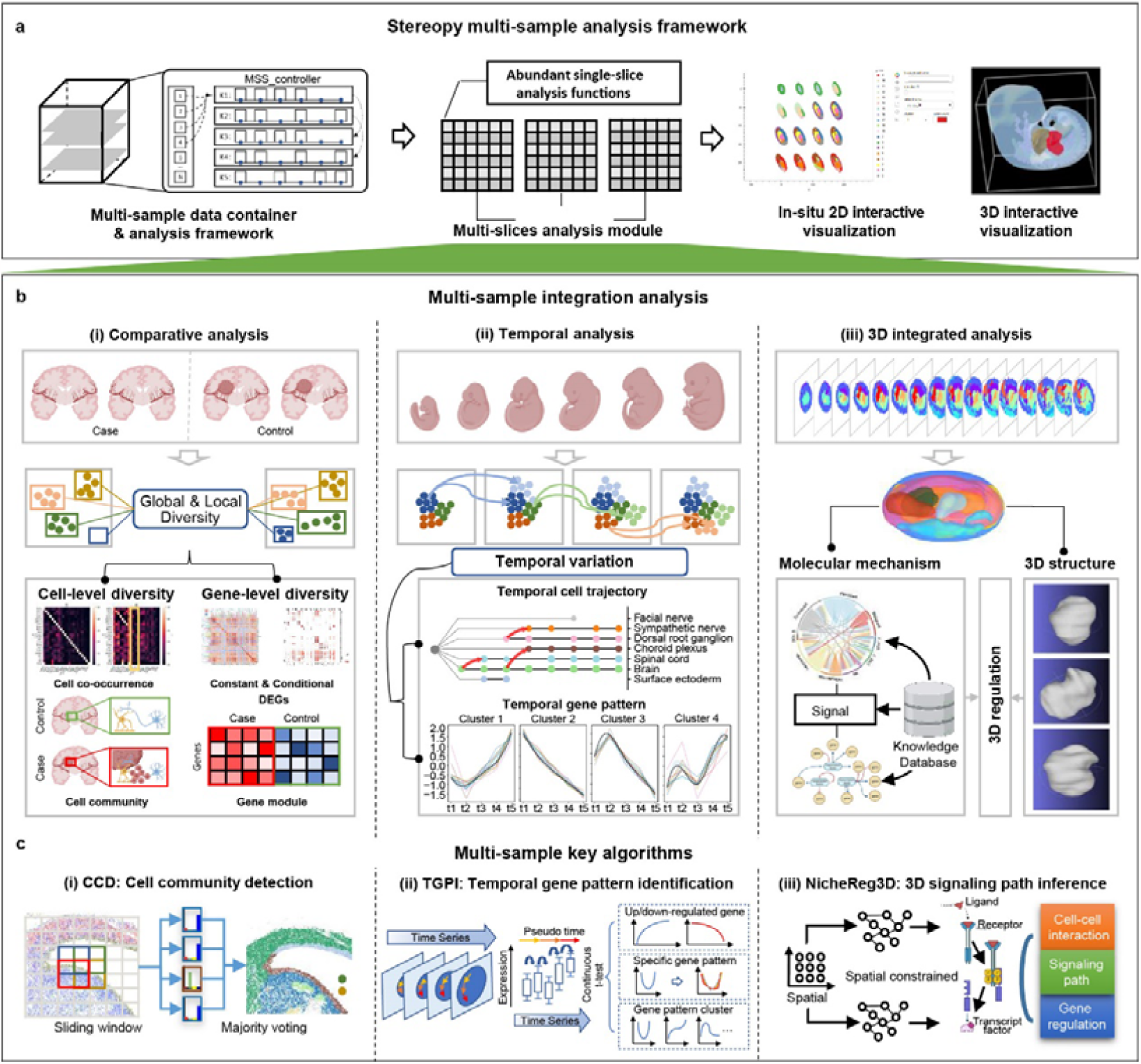
Overview of Stereopy. **a**, Stereopy provides solutions for multi-sample analysis, from multi-sample data container and framework, multi-sample data analysis modules, to multi-sample interactive visualization. **b**, Stereopy offers key analysis modules for three main multi-sample data analysis scenarios. It includes **(i)** Comparative analysis. Stereopy provides functions from cell level and gene level to infer the global and local similarity and diversity for comparative SRT datasets. **(ii)** Temporal analysis. Stereopy provides temporal trajectory analysis and spatial resolved temporal gene pattern analysis to phase the temporal variable datasets. **(iii)** 3D integrated analysis. Stereopy enables 3D data reconstruction and 3D signaling path identification function to explore regulation mechanisms. **c**, Simultaneously, Stereopy contributes the key algorithms for the above three kinds of analysis scenarios. It includes **(i)** Cell community detection (CCD) algorithm aims to detect cell communities on single/multi-sample datasets, which is supposed to find common and specific community, especially for comparative samples. **(ii)** Temporal gene pattern identification (TGPI) algorithm aims to identify temporal variable gene patterns with spatial restriction, which is supposed to find gene pattern related to development or temporal variation. **(iii)** 3D cell-cell signaling path inference tool aims to identify regulation mechanisms from the 3D aspect.

### Stereopy develops an efficient multi-sample data analysis framework

Stereopy offers comprehensive analysis framework to support multi-sample analysis through its MsData container, MSS controller, and multi-sample analysis transformer. The MsData container builds upon the AnnData format, incorporating new features applicable to multiple samples while preserving single-sample dependencies (Fig. 2a). Users can conveniently access the entire dataset and individual samples through a single handler, enabling flexible analysis across multiple samples (Supplementary Fig. 1). The MSS controller plays a critical role in managing result storage, tracking analysis dependencies, and visualizing corresponding outcomes (Fig. 2b). It empowers users to associate meta-information and results generated with corresponding samples for subsequent association analysis. Stereopy’s multi-sample analysis trransformer further supports customized analysis of multi-sample datasets with diverse demands (Fig. 2c), providing functions to integrate single-sample results into the multi-sample context or reversibly split multi-sample data for single-sample analysis. These transformations are particularly useful for analysis modules like clustering and annotation, which may involve manual curations or calculation comparisons by different algorithms. This whole framework facilitates parallel or integrated analysis across multiple samples (Fig. 2d and Extended Data fig. 1a). Meanwhile, it empowers researchers to conduct comprehensive multi-sample joint analyses and interactive visualization on multi-sample data with different demands (Fig. 2e and Extended Data fig. 1b-c). In addition, Stereopy is a powerful tool for dealing with single-sample spatial omics data, providing researchers with an extensive range of analysis functions and sharing several common features and same functions with its multi-sample analysis module (Fig. 2f and Extended Data fig. 1).

**Fig. 2.**
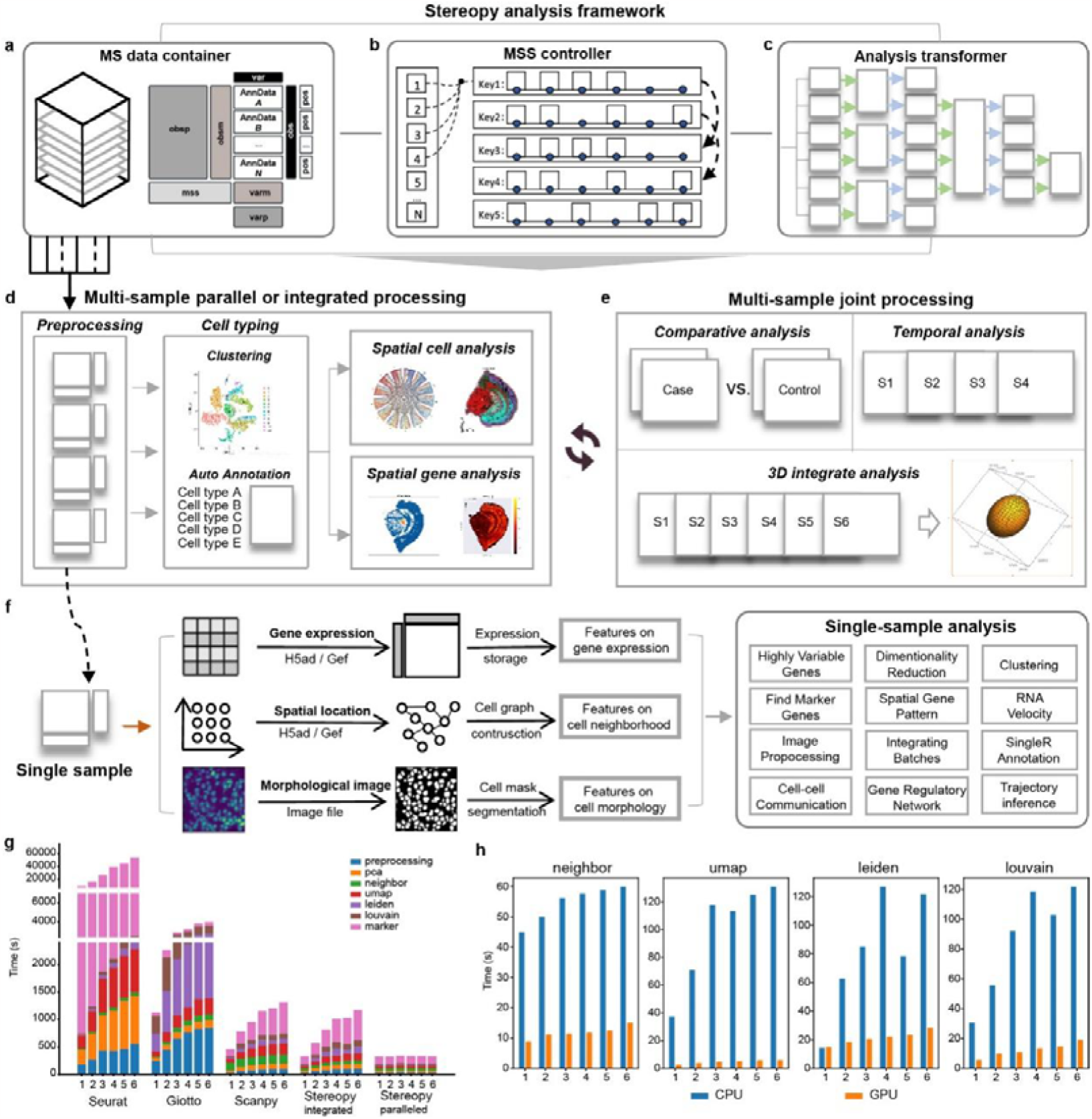
Stereopy designs flexible multi-sample data analysis framework, and accelerates multi-sample analysis. **a**, Stereopy designs multi-sample data (MsData) container, **b**, Multi-sample scope (MSS) controller, and c, Multi-sample flexible analysis transformer to support multi-sample SRT scalable analysis. Based on that, Stereopy is able to support. **d**, Steroepy provides multi-sample parallel or integrated processing and **e**, joint multi-sample processing for comparative analysis, temporal analysis, and 3D integrative analysis. **f**, Stereopy restores and enables processing of gene expression, spatial information and corresponding image features for each sample of spatial omics. **g**, Comparison of execution time of Stereopy, Seurat, Giotto and Scanpy in the basic processes, including preprocessing, PCA, finding neighbors, UMAP, Leiden clustering, Louvain clustering, and finding marker genes. **h**, Comparison of execution time of basic processes, including finding neighbors, UMAP, Leiden clustering, Louvain clustering with GPU mode or without GPU mode.

Stereopy accelerates multi-sample analysis in both algorithmic and parallel computing levels. With the ability to apply parallel analysis to multiple samples for dependent functions, including preprocessing, cell clustering, and annotation, Stereopy significantly reduces the overall processing time. The common SRT analysis modules embedded in Stereopy consume less time for both integrated and parallel processing on different numbers of samples, compared to existing tools such as Giotto, Scanpy, and Seurat (Fig. 2g). Meanwhile, Stereopy supports GPU acceleration for time-consuming but necessary functions such as PCA (Principal component analysis), neighborhood searching, Leiden [25] / Louvain [26] clustering, and SingleR annotation [27] (re-implemented in Python as a part of Stereopy). The GPU-accelerated functions demonstrate a substantial improvement in execution time compared to their CPU counterparts (Fig. 2h).

### Stereopy unveils cell and gene diversity in comparative SRT analysis

In scientific research, comparing disturbed or disease samples with control samples allows for the exploration of changes in functional mechanisms at both local and global levels. Stereopy is designed to analyze and identify global and local diversities in comparative samples by employing cell-level and gene-level analysis modules, supported by novel algorithms (Supplementary Note 1.1). The cell-level analysis focuses on cell diversity in terms of cell type, cell co-occurrence, and cell community via multi-sample comparisons. Stereopy provides enhanced detection of cell communities with our innovative multi-sample Cell Community Detection (CCD) algorithm (Supplementary Fig. 2, Supplementary Note 2.1 and Methods). At the gene level, Stereopy investigates gene diversity within cell types and cell communities (Fig. 3a), proposing the concept of constant and conditional markers. Constant markers exhibit stable expression and remain unaffected by disturbance, while conditional markers respond to disturbances and contribute to functional changes.

**Fig. 3.**
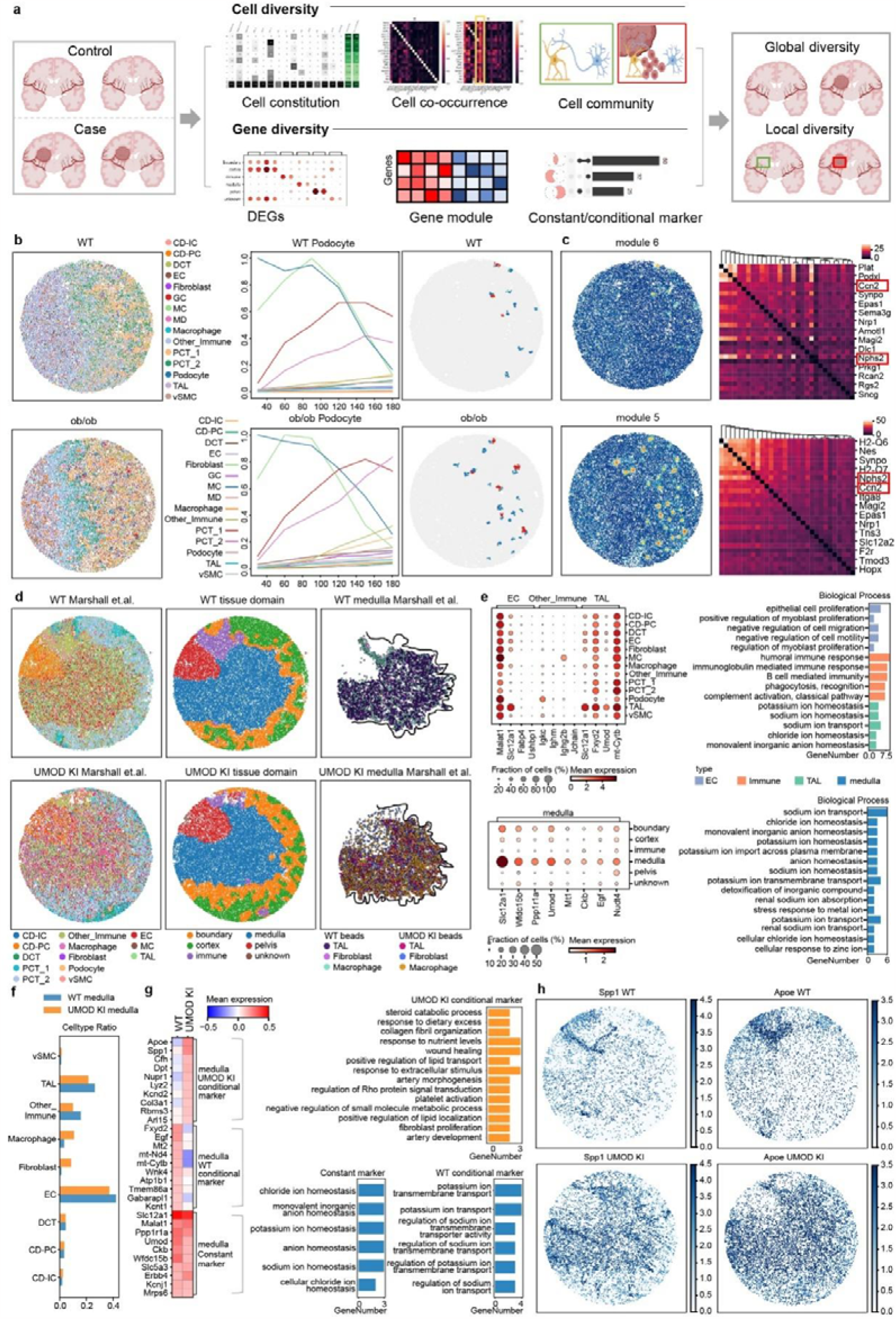
Stereopy facilitates comparative analysis of spatial transcriptomes with multiple samples. **a**, A graphical abstract for Stereopy comparative multi-sample analysis. Stereopy offers analytic functions on the diversity of cell constitution, co-occurrence and cell community at cell aspect as well as differential expression gene, spatial gene module and constant/conditional marker at gene aspect. Combining Stereopy’s complete and comprehensive analysis workflow, spatial pattern diversity can be easily explored at both global and local levels. **b**, Co-occurrence result for BTBR kidney sample. left: spatial map of WT and BTBR diabete (ob/ob) kidney samples; middle: line plot shows the podocyte cell co-occurrence with other cell types; right: spatial map on the right side confirmed the co-occurrence with GCs and MC. Upper and lower part represents WT and *ob* samples, respectively. **c**, Left column: spatial hotspots gene module that corresponds to podocyte location. Right column: local auto-correlation of corresponding gene module. Upper and lower part represents WT and *ob* samples, respectively. **d**, Spatial map of cell type annotation, tissue domain identified by Stereopy-CCD algorithm and medulla defined by Marshall et al. for WT and UMOD KI kidney samples. Left, middle and right part represents cell type annotation, tissue domain annotation and medulla defined by Marshall et al., respectively. Upper and lower represent WT and UMOD KI samples, respectively. **e**, Left part: differential expressed genes for medulla in WT sample as well as its composing cell types EC, TAL and other immune. Right part: GO enrichment for medulla, EC, TAL and other immune. **f**, The cell type constitution and proportion for medulla of WT and UMOD KI samples. **g**, Constant and conditional marker for medulla of WT and UMOD KI samples. Left part shows the heatmap of constant and conditional markers. High expression is only found under certain condition for conditional marker while both conditions have a high expression for constant marker. Right part shows GO enrichment for each group of genes. UMOD KI conditional marker (orange) enriched GO terms including wound healing and so on. **h**, Spatial heatmap of *Spp1* and *Apoe* for WT and UMOD KI samples.

Stereopy-CCD outperforms SpaGCN, GraphST and Giotto in both single-sample scenarios (e.g., whole mouse embryo brain) and multi-sample scenarios (e.g., continuous adult mouse brain and mouse kidney) (Extended Data Fig. 2-4, Supplementary Table 1-2, and Methods). In the whole mouse embryo brain dataset, Stereopy-CCD is capable of effectively identifying cell communities or domains that align with existing knowledge (Extended Data Fig. 2). In continuous adult mouse brain, Stereopy-CCD detected common cell communities among three slides (Extended Data Fig. 3). In our analysis of mouse kidney samples, which included a diabetic sample (UMOD KI-homozygous gene UMOD-C125R knock-in mice with monogenic disorder) and a WT sample [28], the Stereopy-CCD algorithm successfully identified a central community present in both samples (Extended Data Fig. 4). The central community closely corresponds to the region annotated as ‘medulla’ in the study conducted by Marshall et al [28] (Fig 3d and Extended Data Fig. 4).

We applied Stereopy to comparative mouse kidney datasets to assess the its efficacy in detecting global diversity [28]. Co-occurrence calculations developed in Stereopy were performed on a pair of Slide-seq v2 samples: wild-type (WT) and diabetic (*ob/ob* genetic model of early diabetic kidney disease) (Supplementary Fig. 3). The results corroborated Marshall’s previous findings [28] regarding Podocytes’ co-occurrence with GC cells (Fig. 3b) and inferred a higher co-occurrence in the *ob/ob* sample compared to the WT sample. Notably, Stereopy demonstrated a greater significance in detecting the co-occurrence of Podocytes with GCs when compared to Squidpy’s co-occurrence algorithm [14] (Supplementary Fig. 4a-c). Subsequently, gene modules were identified in both samples, revealing the co-expression of *Nphs2* (a Podocyte marker) and *Cgtf* (a Podocyte injury marker) in both the WT and *ob/ob* samples (Fig. 3c). Local autocorrelation analysis revealed a stronger correlation between *Nphs2* and *Cgtf* in *ob/ob* sample (Fig. 3c), and the analysis of differentially expressed genes (DEGs) provided evidence of *Cgtf*’s higher rank among Podocytes markers (Supplementary Fig. 5).

**Fig. 4.**
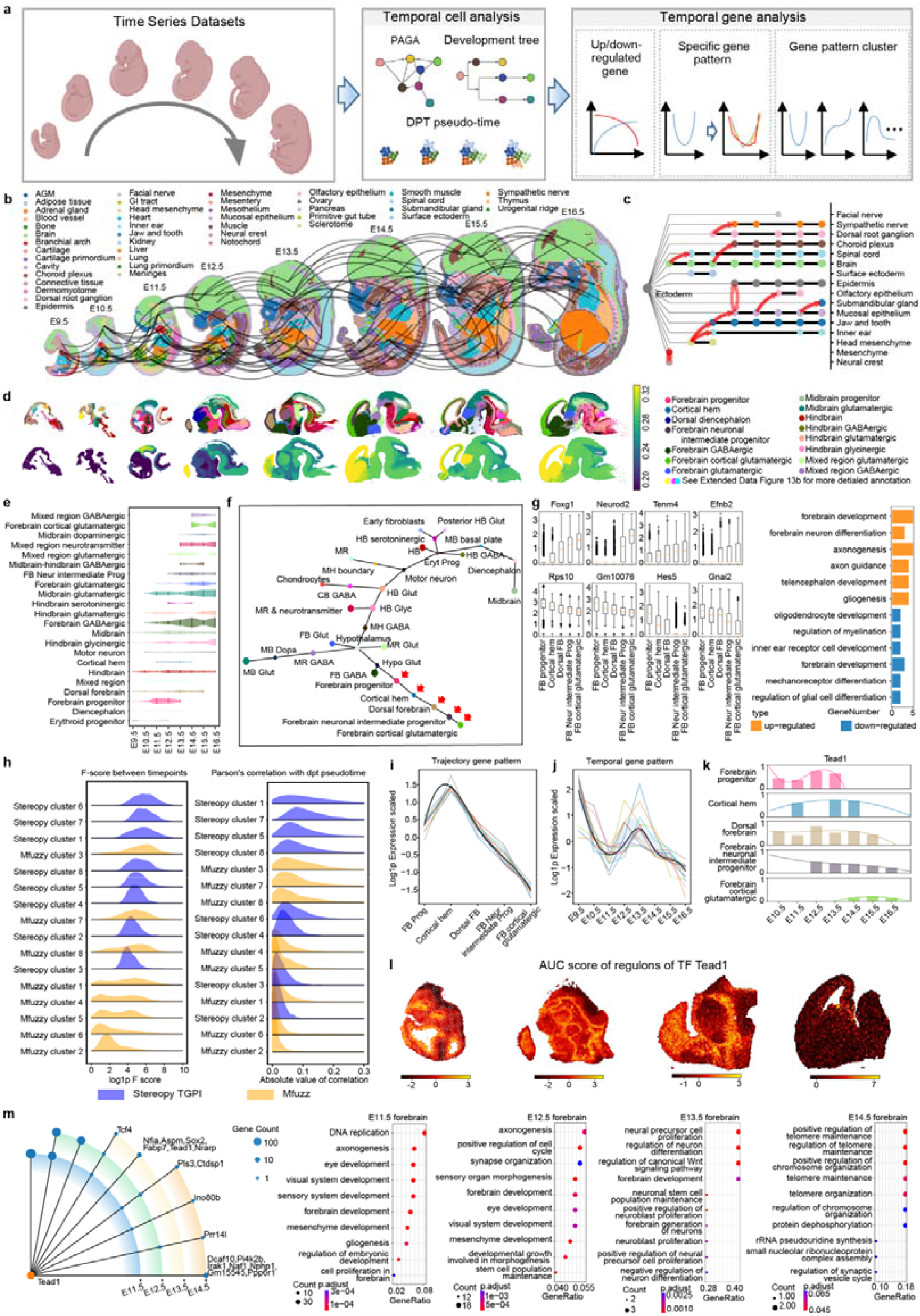
Stereopy enables temporal analysis of spatial transcriptomes with multiple samples. **a**, A graphical abstract for the time series analysis pipeline. For cell aspect, Stereopy integrates PAGA and diffusion pseudotime; for gene aspect, Stereopy proposed an algorithm of spatial resolved temporal gene analysis that can search up or downregulated genes as well as gene clusters with similar temporal patterns. **b**, Spatial trajectory visualization of mouse embryo multi-sample transcriptomes from E9.5 to E16.5. **c**, A tree plot to indicate the develop of mouse embryo ectoderm. X-axis represents time point. Dot size indicate the cell number and red arrow indicate the trajectory from PAGA. **d**, Manual annotation and pseudotime for time series mouse brain samples. **e**, Development tree for cell types in time series. X-axis represents time point. The height of each Sankey represents the cell amount of cell type at a certain time point. **f**, PAGA graph for mouse brain trajectory inference. Red arrow points at cell types for downstream analysis. **g**, Up and downregulated genes for mouse forebrain trajectory and corresponding GO enrichment analysis. **h**, The F-score among time point and correlation with pseudotime of top 1000 gene of each cluster of Stereopy-TGPI and Mfuzz. Blue and yellow represent Stereopy-TGPI and Mfuzz, respectively. **i**, A temporal gene pattern identified by Stereopy-TGPI for mouse forebrain trajectory. **j**, A temporal gene pattern identified by Stereopy-TGPI for mouse forebrain time series datasets. **k**, Gene expression of *Tead1* in each cell type at each time point. **l**, Spatial heatmap for AUC score of TF *Tead1* regulons in each time point and corresponding GO enrichment analysis. **m**, Gene network for *Tead1* in time series. Radial line represents a group of genes and points on it indicate time points when these genes occurred. Point size indicates the gene number.

**Fig. 5.**
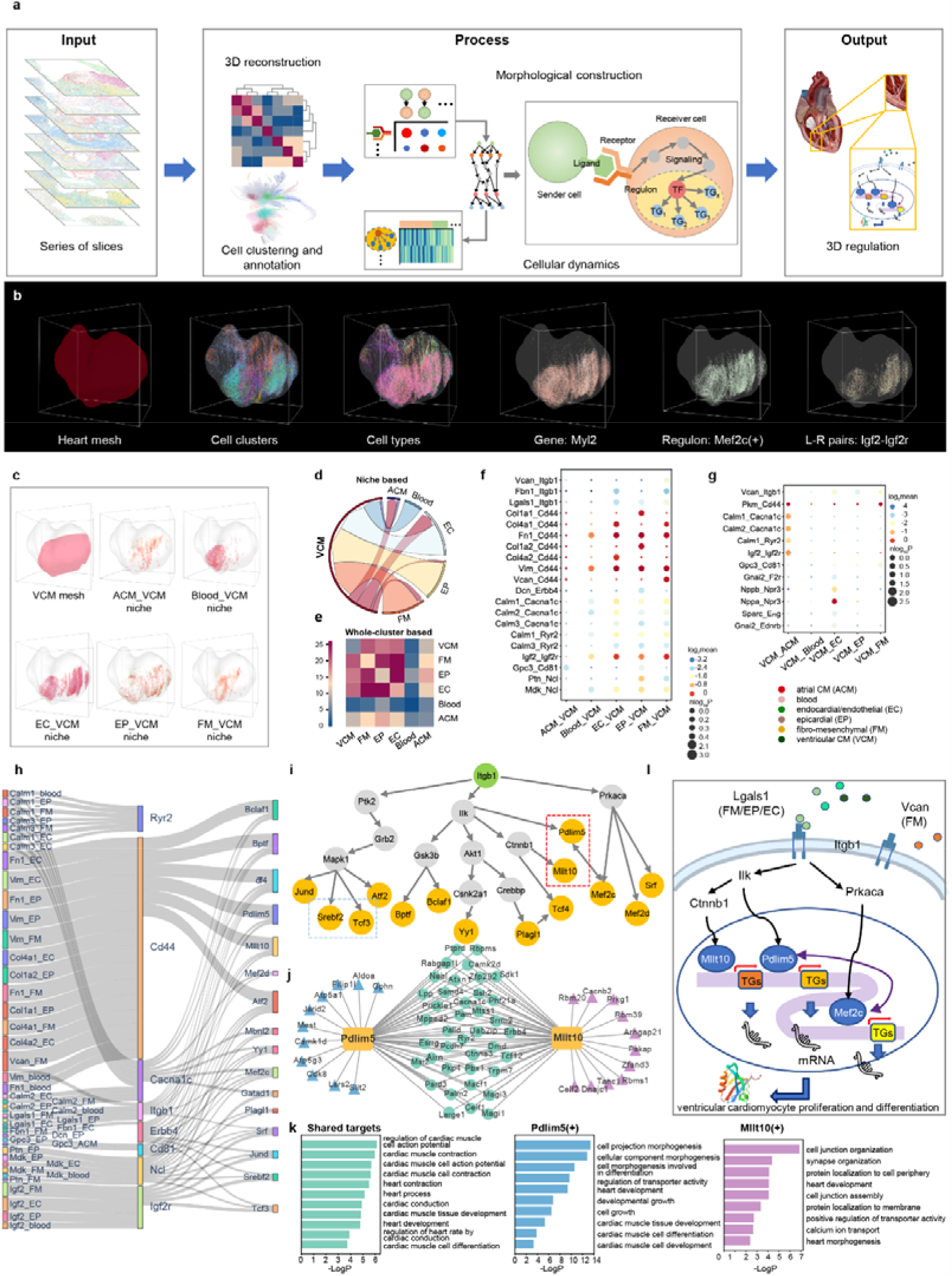
Stereopy integrates spatial multi-sample data and reveals novel 3D regulatory mechanisms related to cardiac development. **a**, Stereopy-NicheReg3D’s overall workflow. **b**, Stereopy-NicheReg3D illustrating multi-hierarchically transcriptomic architecture, ranging from heart organ meshes, heart cell types and clusters, spatially variable genes (*Myl2*), spatially specific regulons (*Mef2c*(+)), and niche-specific L-R pairs (*Igf2-Igf2r*) from left to right. **c**, Spatial distribution of VCM’s niche compositions composed of neighboring ACM, blood, EC, EP and FM cells in the boundary. **d**, Circos plot showing bidirected cell-cell interactions in five niches. The width of an arrow correlates with the number of significant L-R pairs. **e**, Heatmap showing the CCC intensities without niche restriction, which is different from Fig. 5d. **f-g**, Bubble plots demonstrating cell-type-specific L-R pairs. f) niches to VCM. g) VCM to niches. Circle color indicates the mean expression of each L-R pair, while circle size indicates its p-value. **h**, Sankey plot connecting intercellular ligand-receptor interactions from sender niche cells to receiver VCM cells to VCM intracellular downstream TFs via deductive specific signaling pathways. Bandwidth indicates the mean expression of the two genes at both ends. **i**, Regulatory network showing inferred intracellular signaling paths from receptor *Itgb1* to downstream TFs within the same cells. **j**, Shared and specific TGs in *Pdlim5*(+) and *Mllt10*(+) regulons showing the 3D co-regulation function. **k**, GO enrichment analysis indicating the collective function of shared targets of regulons in Fig. 5k (shared, *Pdlim5*(+) and *Mllt10*(+) from left to right). **l**, 3D regulation model of extracellular signaling to intracellular gene regulatory network.

To evaluate the capability of Stereopy of detecting local diversity, we conducted further analysis on the cell communities of mouse kidney samples identified by CCD. We annotated the cell communities according to anatomical structure, including kidney cortex, medulla, the boundary of cortex and medulla, pelvis, and immune region (Fig. 3d). As previous mentioned, the medullary region identified by Stereopy-CCD closely corresponds to the region annotated in the study conducted by Marshall et al. and the marker genes of each region are clearly discernible (Fig 3d, Extended Data Fig.4 and Supplementary Fig. 6). The community ‘medulla’ exhibited similar proportions of thick ascending limb (TAL), endothelial cells (EC), and other immune cell types, despite the diversity in cell type distribution (Supplementary Fig. 7). The UMOD KI sample showed increased percentages of fibroblast and macrophages in the medulla compared to WT (Fig. 3f and Supplementary Fig. 7), consistent with Marshall’s findings [28]. To analyze markers of both tissues and cell types, we calculated DEGs and enriched gene ontology (GO) terms specifically for the medulla and its constituent cell types, such as TAL, EC, and other immune cell types (Fig. 3e). The marker genes exhibited greater significance, and the enriched GO terms in the renal medulla were highly relevant to renal function, including sodium ion transport, potassium ion transmembrane transport, and chloride ion homeostasis, highlighting the functional relevance of thetissue compared to separate cell types. With that finding we confirmed that examining the gene divergence of cell communities provides deep insights into tissue function. To comprehensively assess the response of various regions to the UMOD KI disturbance, we analyzed the number of conditional markers. Our calculations indicated a substantial increase in the number of top DEGs in the medulla region compared to other regions, along with a greater diversity of cell types (Supplementary Fig. 8). These findings suggest that examining the overall differences of cell communities and condition markers may yield more meaningful biological discoveries than focusing solely on individual cell types. Specifically, we observed a consistent marker related to renal function, including sodium, potassium, and chloride ion homeostasis, in the renal medulla across both healthy and disease samples. However, the UMOD KI sample exhibited conditional markers involved in renal function damage, such as response to nutrient level, wound healing, and response to extracellular stimulus (Fig. 3g and Supplementary Table 3). Notably, *Spp1* emerged as a significant conditional marker (Fig. 3h), which has been proved as the top hub gene associated with Kidney stone disease [29]. Further analysis revealed that renal stone risk persisted when both *Spp1* and *Umod* had variants, indicating the importance of these two genes in the development of kidney disease [30]. Another conditional marker, *Apoe*, was reported to be associated with glomerular disorders due to its central role in lipoprotein metabolism. The increased abundance of macrophages in the UMOD KI sample is consistent with the hyperactivity of macrophages involved in *Apoe*-related glomerular disorders [31].

In conclusion, Stereopy provides a systematic analysis of cell-level and gene-level similarity and diversity between case and control samples, with high biological significance. The use of the Stereopy-CCD algorithm and identification of conditional markers significantly contribute to the understanding of tissue structure and function in comparative analyses.

### Stereopy identifies spatiotemporal variation in time-series SRT analysis

Growth and development of organisms involve complex biological processes characterized by Variations in cell types and gene expression over time. These intricate molecular changes described as temporal variation, captures the intricate molecular changes occurring during development. To investigate temporal variations in time-series datasets, Stereopy emphasizes dynamic changes in both the spatial and temporal dimensions (Supplementary Note 1.2). In terms of cell type changes, Stereopy adopted a manifold partitions-based method [32] to preserve the global topology and to infer the trajectory of cell types across different samples, providing a visual representation of cell trajectory and changes in cell numbers across time points (Supplementary Note 3). Meanwhile, Stereopy proposes a spatially resolved Temporal Gene Pattern Identification (TGPI) method for finding genes with similar temporal expression changes, including continuous up- or downregulated genes, as well as other complex patterns observed in real time and pseudotime (Fig. 4a and Supplementary Note 2.2).

Our proposed statistic metric, the plain false discovery rate (pFDR), has been utilized in Stereopy-TGPI for detecting continuous up- and down-regulated genes by merging p-values. To evaluate its effectiveness, we compared pFDR with Fisher’s method using mouse embryo forebrain datasets consisting of three cell types with 7, 5, and 3 time points, respectively. The results demonstrated that pFDR is a more stable and reliable approach for identifying genuine up- and down-regulated genes with continuous changes in gene expression across multiple time points (Extended Data Fig. 5, Supplementary Fig. 9-10 and Methods). Stereopy-TGPI serves as a valuable tool for not only identifying temporal gene up- and downregulation but also for elucidating intricate temporal or pseudotime expression patterns. Stereopy-TGPI simultaneously considers the consistency of gene expression in both temporal and spatial aspects (Methods). We conducted an evaluation of the significance of spatial features in Stereopy-TGPI and found that they play a crucial role in enhancing the consistency of temporal gene pattern detection (Extended Data Fig. 6 and Supplementary Fig.11). Compared with Mfuzz, Stereopy-TPGI’s identification was more correlated to real and pseudotime tendencies and capable of enriching significant GOs relevant to neuron development in the time-series whole mouse brain (Extended Data Fig. 7 and Methods) [33].

To illustrate the capability of Stereopy, we investigated the trajectory and temporal gene pattern across eight time-point samples of Stereo-seq mouse embryos from E9.5 to E16.5 [3]. We inferred the trajectory of the integrated mouse embryo dataset from eight time points and displayed the cell type development with a tree plot (Fig. 4b-c). Next, the flexibility of Stereopy’s data container enabled manual clustering and annotation for the brains of each sample, independently (Fig. 4d, Supplementary Fig. 12a). Pseudotime analysis [32] verified a gradual increase in pseudotime, and higher pseudotime values in the forebrain region indicated later development (Fig. 4d, Supplementary Fig. 12b). Additionally, we provided statistics on the cell number of each cell type across time points and inferred the cell trajectory for the time-series mouse brain development (Fig. 4e-f and Supplementary Fig. 12c). We then focused on the temporal gene pattern in the forebrain trajectory series, encompassing the forebrain progenitor, cortical hem, dorsal forebrain, forebrain intermediate progenitor, and forebrain cortical glutamatergic stages.

Applying Stereopy-TGPI to calculate gene up- and downregulation, we identified *Foxg1* as the top-ranked temporal upregulated gene, a key transcription factor (TF) that showed gradual upregulation along the forebrain trajectory, consistent with its dynamic regulation of forebrain development [34] (Fig. 4f-h and Supplementary Fig. 13). Conversely, *Hes5*, a gradually downregulated gene, exhibited high expression in embryonic neural precursor cells and played a crucial role in negatively regulating neural and oligodendrocyte differentiation [35]. Furthermore, we applied Stereopy-TGPI to identifying cell-type-specific expression patterns along the forebrain trajectory. Among them, a distinctive cell-type-trajectory gene pattern was observed, with characterized genes exhibiting an upregulation trend prior to the cortical hem stage, followed by continuous downregulation thereafter. GO terms enriched in this pattern included neural precursor and forebrain cell proliferation, implying an important role for cortical hem during forebrain development (Fig. 4g). We observed cortical hem existed exclusively before E14.5 in this dataset, which aligned with a previous study that reported the emergence of Cajal-Retzius neurons, the constituent cell type of the cortical hem, during the early developmental stages [32]. To investigate the functions of cortical hem and key factors leading to its disappearance, we examined temporal gene patterns related to time points with consistent occurrence of cortical hem at the forebrain. A distinct temporal gene pattern was observed, with peak expression levels during the developmental stages from E11.5 to E14.5, coinciding with the presence of the cortical hem and a noticeable decrease thereafter (Fig. 4i). We intersected genes from the cell-type-trajectory gene pattern and temporal gene pattern. Within this intersection, we identified *Tead1* as a key TF that exhibited high expression in cortical hem and low expression after its disappearance (Fig. 4j). *Tead* TFs have previously been implicated in regulating cortical development [33]. Leveraging the interactive visualization capabilities of Stereopy, we performed gene regulatory network (GRN) analysis on the forebrain of each sample by easily selecting regions of interest (Supplementary Fig. 14). Strikingly, we observed a decrease in the number of genes regulated by *Tead1*, from 338 at E12.5 to 7 at E13.5 (Fig. 4k). The enriched GO terms at E11.5 and E12.5 were similar and highly associated with forebrain development, while those at E13.5 were related to neuroblast proliferation and forebrain neuron generation. For example, *Tcf4*, a target gene (TG) regulated by *Tead1* from E11.5 to E13.5, controls the positioning of cortical projection neurons [38]. However, by E14.5, GO terms were no longer related to neurons or the brain (Fig. 4l). Notably, E14.5 marked the appearance of forebrain cortical glutamatergic appeared, suggesting that *Tead1* and cortical hem had finished their neurogenesis function. Our findings underscore the significance of *Tead1* and cortical hem in forebrain cortical development, providing valuable insights into brain development.

Stereopy offers comprehensive solutions for temporal multi-sample analysis, with a particular emphasis on the temporal gene patterns. TGPI, a key feature of Stereopy, enables the detection of temporal gene patterns within time-series datasets, facilitating the exploration of dynamic changes across various time points. By leveraging Stereopy’s TGPI functionality, researchers can systematically investigate and uncover developmental-related temporal gene patterns, thereby revolutionizing our understanding of temporal dynamics in biological systems.

### Stereopy reveals principles of niche-mediated regulations in 3D SRT analysis

Multicellular organisms are inherently comprised of cells and tissues organized within a 3D structure, thereby giving rise to intricate cellular interactions that cannot be adequately replicated in 2D culture [39]. Unfortunately, conventional analytical approaches remain confined to 2D methodologies, inevitably resulting in the loss of crucial interaction information along the z-axis. However, Stereopy’s NicheReg3D pipeline addresses this limitation by precisely characterizing the cellular constitution of 3D niches and facilitating the comprehensive exploration of intercellular and intracellular interactions (Supplementary Note 1.3). It seamlessly combines data preprocessing, 3D alignment and reconstruction, cell-niche communication, ligand-receptor (L-R)-TF-TG pathway inference, and intracellular TF-centered regulatory network prediction (see Methods). The core algorithms underpinning 3D joint analysis and the underlying 3D regulation model are elucidated in Fig. 5a. When applied to the well-studied system of the mouse cortical region sequenced by BARseq, a high-throughput in situ sequencing technique [40], our results demonstrated that 3D niches, composed of complete and accurately defined cells from diverse cell types after 3D reconstruction of these consecutive 2D slices. It outperformed 2D niche composition analysis of each individual slice, benefiting downstream analyses such as the predictive identification of cortical areas (Supplementary Fig. 15).

In the 3D context, cellular heterogeneity is not only governed by the intracellular regulatory network but also influenced by the extracellular microenvironment to collaboratively accomplish diverse biological tasks [41, 42], yet computational methods for modeling both interactions simultaneously are insufficient [43]. Our approach enables the joint analysis of spatial multiple samples in 3D, providing unique insights into the intracellular regulation mediated by biochemical signals of intercellular crosstalk in multiple dimensions. To showcase its effectiveness in exploring niche-mediated regulations, we applied Stereopy-NicheReg3D to analyze the cardiac development of a mouse embryo sequenced by Stereo-seq [3]. We extracted 59 10-μm-thick 2D serial cryosections at a distance of 10 μm, covering the entire mouse embryonic heart. For the SRT data of 90,411 high-quality segmented cells with 30,254 genes inferred from subcellular spots, we performed unsupervised clustering analysis for each individual sample and identified six cardiac cell clusters (Supplementary Fig. 16 and Supplementary Table 4). The 3D reconstructed model provided a multi-hierarchical transcriptomic architecture, ranging from organ meshes, cell types and clusters, spatially variable genes, spatially specific regulons, and niche-specific L-R pairs (Fig. 5b).

Based on the reconstructed 3D murine heart data, our investigation focused on the development of ventricular cardiomyocytes (VCMs), in which the cardiac niche played significant roles in cardiogenesis through intricate intercellular signal transduction [44]. The VCM niche encompassed five other cell types: approximately 28% atrial cardiomyocytes (ACMs), 27% blood cells, 23% endocardial cells (ECs), 13% epicardial cells (EPs), and 9% fibro-mesenchymal cells (FMs) (Fig. 5c). Within a 25-μm physical distance in 3D, expressed L-R pairs revealed that VCMs were the prominent receiver of signals from surrounding cells (Fig. 5d). This finding contradicted the conventional understanding of communication activities specific to whole cell clusters [45] (Fig. 5e), underscoring the essentiality of 3D spatial neighborhood pruning. Notably, our computational analysis predicted a significant molecular interaction between the ligand *Vcan* and its receptor *Itgb1* in FM-VCM cells, with moderate presence in other niches (Fig. 5f). This observation aligns with previous studies that have emphasized the critical role of *Vcan* in the extracellular matrix for supporting and remodeling VCMs [46, 47]. More interestingly, apart from ACMs, the four distinct niche compositions collectively influenced VCM gene expression through the same L-R pair sets (*Vim-Cd44, Calm1-Ryr2, Igf2-Igf2r*…), most of which have been implicated in the regulation of CM proliferation, migration, and differentiation [48, 49] (Extended Data Fig. 8 and Supplementary Fig. 17). Compared to state-of-the-art CCC tools, including single-cell CellPhoneDB [45] and spatially resolved NICHES [50] (Supplementary Table 5), Stereopy achieved the most complete identification of specific L-R pairs that covered nearly all those derived by other tools, thanks to the precise niche extraction (Extended Data Fig. 9). The majority of these L-R pairs are involved in mammalian cardiac growth and development (Supplementary Table 6). On the other hand, VCM reversely influenced the cell state or function of the cell microenvironment through specific L-R pairs (Fig. 5g).

Furthermore, we inferred the specifically expressed GRNs on VCM cells adjacent to the niches (Fig. 5h and Supplementary Fig. 18 and Supplementary Table 7). This enrichment analysis yielded a set of candidate core TFs and their corresponding regulons, suggesting their potential susceptibility to cell-niche communications and warranting further inspection of their regulatory effects. We then established deductive signaling paths to connect intercellular signaling activities from niche cells with intracellularly influenced TFs. To simplify the complex network, we retained connections between receptors and TFs involving a maximum of two intermediate genes (Fig. 5i). Among them, *Cd44* emerged as the recipient of the most extracellular signaling, stimulated by specific ligands (*Col1a1, Col4a1/2, Vcan*) or collectively expressed from different niches (*Fn1, Vim*). This signaling could up- or downregulate various TFs such as *Tcf4*. Previous studies have elucidated the ability of *Cd44* to activate the canonical Wnt/β-catenin signaling pathway, impacting the expression of *Tcf4* and downstream genes [51], thereby exerting temporal and spatial control over heart maturation [52]. *Igf2-Igf2r* also collectively regulated VCM proliferation and differentiation by activating PI3K/Akt pathways, as previously reported [53, 54]. Moreover, the shared *Calm1/3-Cacna1c* family displayed potential regulation of *Mef2c/d* expression, which has been linked to excitation-contraction coupling in VCM function through calmodulin-dependent signaling pathways [55, 56]. Our framework additionally facilitated the investigation of detailed GRNs for each user-defined receptor in the same cell. For instance, Fig. 5j depicted the GRN of the *Itgb1* receptor as a directed graph, encompassing various modes, including directed acyclic (such as *Srebf2* and *Tcf3*) and bidirected acyclic (such as *Pdlim5* and *Mllt10*). Importantly, the inferred GRN, extended to downstream TGs, highlights the potential for intercellular communication to regulate the same set of genes, culminating in collective regulation (Fig. 5k and Supplementary Fig. 19). For example, *Itgb1*-related CCC might modulate both *Pdlim5* and *Mllt10* through Ilk-related pathways. GO enrichment analysis indicated that their shared TGs jointly managed cardiac muscle development and contraction (Fig. 5l), corroborating prior findings [57, 58]. In contrast to other tools connecting the outside and inside of the cells, such as NicheNet [59], Stereopy-NicheReg3D provided a more definitive and complete network for inferring how cell-niche-specific L-R pairs regulate intracellular regulon activities related to specific cellular functions.

In this scenario, we have witnessed that our 3D joint analysis pipeline explores how spatially informed extracellular signaling at the niche influences intracellular gene regulation in the cell of interest, beyond the limitations of 2D data analysis (Extended Data Fig. 10 and Supplementary Fig. 20). The integration of CCC and GRN could presumably improve the accuracy of context-specific L-R-TF-TG predictions concerning morphological phenotypical changes. As such, we derived an improved model of 3D regulation implicating VCM development in cardiac maturation and physiology (Fig. 5m). During heart development, VCMs constitute a fundamental element of heart function, while EC, EP, FM, and blood cells are key components of the microenvironment promoting CM maturation. Niche components collectively or specifically transmit signals through shared or distinct L-R pairs, which further promote or inhibit specific TFs inside VCM cells through specific signaling pathways. These TFs ultimately influence the expression of downstream TFs and TGs, jointly demonstrating the cellular functional state and subtype.

Therefore, we anticipate that Stereopy-NicheReg3D will serve as a valuable tool with an interactive visualization browser in the 3D space (Supplementary Fig. 21) for better dissecting the functional consequences of spatially informed inter-intracellular regulation networks, thereby facilitating the prediction of cellular function, state, and corresponding phenotype.

## Discussion

The interpretation of similarities, differences, and developmental changes across multiple samples is non-trivia to unravel complex biological regulatory mechanisms using multi-sample spatial omics datasets. In this study, we introduce Stereopy, a comprehensive toolkit for managing, analyzing, and visualizing multi-sample spatial omics data. It offers the MsData container, MSS controller, and a multi-sample analysis transformer, effectively addressing the challenges encountered in jointly analyzing multi-sample data. Stereopy also provides a wide array of analysis solutions and algorithms tailored specifically for comparative, temporal, and 3D integrated analysis in multi-sample endeavors.

Firstly, we employed Stereopy on comparative kidney datasets to validate the co-occurrence of Podocytes with GCs and identified *Spp1* as a potential significant UMOD KI conditional marker. The Stereopy-CCD algorithm proved its efficacy in detecting important cell communities across multiple samples, thereby expanding the scope of diversity analysis in comparative studies. Subsequently, we harnessed the capabilities of Stereopy to delve into temporal datasets, highlighting the function of *Tead1* and the cortical hem in forebrain cortical development. This investigation provides valuable insights into the intricate dynamics of mouse forebrain development using mouse embryonic brain datasets. The Stereopy-TGPI algorithm demonstrated its ability to accurately infer temporal gene patterns by integrating spatial information, thereby revealing potential gene patterns and key TF genes related to forebrain development. Finally, we leveraged Stereopy to explore the 3D multi-sample datasets, specifically investigating the developing ventricular cardiomyocytes in the mouse embryonic cardiac dataset. Through this analysis, we identified an *Itgb1*-stimulated co-regulation network, illuminating the intricate inter- and intracellular regulatory mechanisms in the 3D niche-based microenvironment. The Stereopy-NicheReg3D pipeline proved its superiority in identifying more complete specific LR pairs and comprehensive signaling paths compared to existing tools when applied to 3D datasets.

Stereopy represents a comprehensive and robust solution that surpasses the mere provision of functionalities and algorithms for analyzing complex spatial omics datasets. Its advanced features, including batch effect evaluation and removal processing of multiple samples, as well as multi-sample joint analysis functions such as 3D registration, 3D data trajectory inference and visualization, amplify the utility of Stereopy in the field. Moreover, Stereopy incorporates numerous data analysis functions, including several well-known functions adapted from R code, such as scTransform and SingleR. Additionally, Stereopy can handle diverse data types, including GEF and GEM files generated by Stereo-seq, as well as the commonly used h5ad file format, enabling the analysis of data from different platforms. It is worth noting that Stereopy can analyze SRT datasets as long as they provide both spatial information and gene expression at the same resolution. However, some algorithms bundled within Stereopy expect high-resolution datasets as input for optimal performance, rendering it more suitable for high-resolution than low-resolution spatial omics datasets.

Stereopy has effectively tackled key challenges in multi-sample spatial omics analysis, including data management, analysis module planning, algorithm development, and interactive visualization of 2D/3D data. Nonetheless, there are opportunities for further improvement to enhance and enrich multi-sample data analysis by accommodating new modalities, addressing new analysis demands, and incorporating new omics to support scientific research. It is imperative to leverage spatial and feature information, particularly in spatiotemporal datasets (referred to as 4D datasets), to unlock insightful biological discoveries. Stereopy is committed to expanding its analysis functions and extending its applications to diverse areas, including clinical and immune research. The support for multimodal analysis and multi-omics datasets should be prioritized as they provide richer biological information and represent the future of spatial omics technologies.

Although research involving multi-sample datasets is commonplace, the research community dedicated to multi-sample analysis remains relatively underdeveloped. This deficiency can be attributed to the absence of a standardized multi-sample analysis framework that seamlessly integrates various analysis tools and elucidates the canonical forms of multi-sample multi-omics analysis. Additionally, the integration of certain algorithms and tools into a unified framework poses significant challenges. Consequently, the joint analysis for multiple samples becomes a formidable hurdle, compelling researchers to either forego the valuable insights embedded within multi-sample datasets or invest substantial time in searching for appropriate analysis tools and determining the optimal analysis framework. Stereopy emerges as a foundation for building a vibrant multi-sample omics community and promotes the establishment of canonical forms for data analysis. Meanwhile, the introduction of the developer mode invites contributions from the expansive bioinformatics community, fostering collaborative efforts. With unwavering dedication, Stereopy strives to furnish researchers with a user-friendly analysis toolkit and robust analysis modules. Simultaneously, it offers novel perspectives and profound insights into the interpretation of multi-sample spatial omics data, empowering researchers to unlock the full potential of these datasets.

## Methods

### Comparison of general single-cell analysis between Stereopy, Scanpy, Seurat, and Giotto toolkits

Many toolkits have provided functions for single-cell or spatial transcriptomic analysis. Scanpy is a widely used package for single-cell analysis in Python while Seurat [4] is in R. Giotto [2] is also an R package with specific designs for ST. In order to figure out the time consumption performance among toolkits including Stereopy, Scanpy, Seurat, and Giotto, we tested the most general analysis in a single cell and SRT including pre-processing, principal components analysis (PCA), Uniform Manifold Approximation, and Projection (UMAP), cell neighbors finding, Louvain clustering, Leiden clustering and find gene markers. To test performance on multiple samples, we use Stereo-seq mouse embryo datasets from E9.5 to E14.5. Since only Stereopy provides analysis on multi-sample data, we test other toolkits by merging multi-sample data as one data. To have a fair competition, we kept all hyper-parameters the same. For pre-processing, we test 3 steps: normalize, log1p, and scale. For PCA, we retained the top 30 principal components (PCs) without using highly variable genes. For cell neighborhood, we calculate based on PCA results with 20 top PCs and 10 nearest neighbors. For Louvain and Leiden, the resolution is set to be 1 as default. To find gene markers, we test on pre-annotation clustering results and use a t-test based on all versus rest way. All of the toolkits are tested on a Linux machine with 64 cores CPU and 512 GiB of RAM.

### Cell co-occurrence detection algorithm

To explore the changes in cell neighborhood, we developed a global co-occurrence method (Supplementary Fig. 3) to reflect the spatial distribution relationship between cell types or clusters. The presented co-occurrence method is composed of 3 steps: 1. Calculation of cell-to-cell spatial distance, 2. Spatial graph construction, and 3. Counting of cell-type contacts. For the first step, we calculate a cell-cell pairwise spatial distance matrix based on Euclidean distance. Secondly, with the distance matrix used as the adjacent matrix of cell neighborhood graph, we only retain edges with a distance range from minimal distance threshold to maximal distance threshold. The minimal and maximal distance thresholds could be selected manually. After constructing the cell neighborhood graph, we calculate the probability that cell type A has the edge with cell type B. This probability represents the co-occurrence probability of cell type A with cell type B. The following equations explain this process in more detail. We mark cells belonging to the cell type A[0, N] and B[0, M] as:

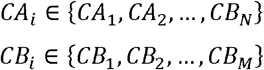

Cell counts of cell type A are given by the number of A cells that are located around cell type B from the minimum distance to the maximum distance:

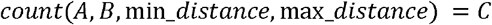

The co-occurrence of A with B:

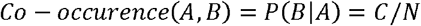

Notably, the co-occurrence of A with B is not equal to the co-occurrence of B with A, which equals e/n and e/m respectively in our method. The asymmetry of our co-occurrence stands because the spatial distribution of a cell type includes another cell type and is more universally distributed. For example, ECs are universally distributed in mouse kidney which means most other cells are connected to ECs while not every EC is connected to other cell types. Owing to the asymmetry of our co-occurrence method, we can also detect the wideness of spatial distribution for cell types. Additionally, we created the co-occurrence result integration method for multiple samples based on the weighted mean of the group, where weights equal to the 17 cell counts ratio in multi-sample data

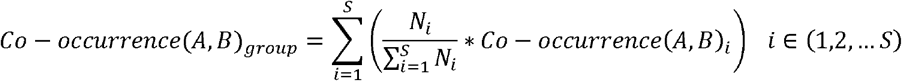

The grouped co-occurrence is equal to the result of merged multiple samples calculated per sample since all samples originate from the same tissue. On the other hand, we use the difference to indicate the co-occurrence between two samples or two groups. The differential co-occurrence value ranges from -1 to 1, where the positive value represents improvement of co-occurrence, and vice versa.

### Benchmark of co-occurrence algorithm

To compare the performance of cell type co-occurrence of Stereopy with Squidpy, we tested on mouse kidney WT and BTBR samples [28]. Since there is no ground truth for cell co-occurrence. we compared the results with previously reported findings. The co-occurrence is calculated based on the cell spatial neighborhood and the distance traverse from 0 to 180 in steps size of 30, unit same as the resolution of slide-seq V2 technology which is 10μm (Supplementary Fig. 4a). For Stereopy, we use *co-occurrence* function with default parameters while for Squidpy we use *co_occurrence* with parameters *spatial_key = ‘spatial’, interval = np*.*array([0,30,60,90,120,150,180])* and *n_splits = 1*. As a result, Stereopy shows a more obvious co-occurrence of podocytes and GC cells than Squidpy, which is consistent with Marshall’s findings [28]. In addition, with the help of grouped and differential co-occurrence among multi-sample analysis, Stereopy is capable of finding the similarities and diversities of cell type co-occurrence among multiple samples. Compared to the significant decrease in co-occurrence of GC, MC with itself in Squidpy, Stereopy can exhibit more significant changes between multiple articles, such as the reduction of co-occurrence between PCT_ 1 and PCT_2. (Supplementary Fig. 4c)

### Cell community detection (CCD) algorithm

The function of the tissue is tightly coupled with the cell populations inhabiting it. The cell neighborhood largely affects the essential gene expression patterns of each cell [15]. For that reason, detecting areas of the tissue with similar cell type distribution and cell type co-occurrence represents an important finding about the structure and function of the tissue. The main idea behind defining functional tissue domains (communities) can be narrowed to detecting tissue areas with the same cell mixture (percentages of cell types). For this purpose, we developed a Cell Community Detection (CCD) algorithm that uses annotated cell types together with spatial coordinates of each cell-spot to assign community labels. CCD divides the tissue using sliding windows by accommodating multiple window sizes and enables the simultaneous analysis of multiple samples from the same tissue. It consists of the three main steps (Supplementary Fig. 2):

1. Single or multiple-size sliding windows (*w*) are moved through the surface of the tissue with defined horizontal and vertical steps while calculating the percentages ([*p*_1_, *p*_2,_…, *p*_n_]) of each cell type inside of it. A feature vector (*fv*) with a size equal to the number of cell types (*n*) is created for each processed window across all available tissue samples:

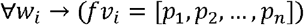
2. Feature vectors from all windows are fed to the clustering algorithm (*C*) such as Leiden [25], Spectral [60], or Hierarchical [61] to obtain community labels (*l*). The number of the desired communities (*cn*) can be predefined explicitly as a parameter (Spectral or Hierarchical clustering) or by setting the resolution of clustering (Leiden):

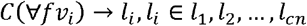
3. A community label is assigned to each cell-spot (*cs*) by majority voting (*MV*) using community labels from all windows covering it:

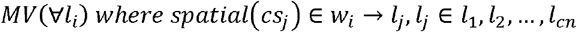

The window size and sliding step are optional CCD parameters and when not provided the optimal window size is calculated throughout the iterative process. In the first iteration, the initial window size is obtained by dividing the minimum of *x* and *y* spatial coordinates’ ranges by 100 and rounding to the closest even number. Then, we calculate for each window of the obtained size the average number of cells being covered by it. If the average number is below 30 the window size is increased by 10% and, if larger it is decreased by 10%. The step is repeated until the average number of cell spots in all windows is in the range [30, 50]. The sliding step is set to half of the window size.

CCD also includes several filtering steps controlled with parameters, such as the removal of cell types present in all parts of the tissue and removal of windows with too small number of cell spots. The spatial distribution of each cell type can be evaluated using 2D entropy [62, 63] and scatteredness [62] metrics. CCD supports setting the threshold values for these metrics in order to exclude cell types that are randomly or evenly spread throughout the tissue from processing. Removing cell types with high entropy and scatteredness improves clustering and provides more robust cell communities. The robustness and quality of CCD strongly depend on clustering. For clustering to be stable, feature vectors need to contain a significant amount of information, that is, enough cell spots in each evaluated window. CCD gathers data on total cell numbers per window and supports setting a threshold value for the minimum cell-spot number for the window to be included in the clustering process. Cell-spots are marked with the ‘unknown’ label if there are no cell community labeled windows that overlap them.

### Benchmark of cell community detection algorithm

To assess the stability and reliability of Stereopy’s CCD, we conducted a comparison with existing algorithms for domain detection on three samples: single sample Stereo-seq mouse embryo whole brain [3], Slide-seq V2 UMOD KI kidney comparative samples [28], and Stereo-seq multi-sample adult mouse brain [64] (Supplementary Note 2.1). For single sample, we included Giotto’s Spatial Domain Identification (GSDI) [15], SpaGCN [65] and GraphST [23]for comparison. In addition, for the multi-sample analysis, we included PRECAST [21] and BASS [22] (Supplementary Note 2.2). Giotto and SpaGCN only support single-sample processing, creating results that require cluster matching to support further analysis. Both GraphST, BASS and PRECAST are able to process multiple slices simultaneously. CCD is able to process single sample as well as multiple samples simultaneously. SpaGCN was run with the default parameters (resolution = 1.5). Giotto’s SDI required adjustment of gene expression and cell location data to a defined input format. Data was normalized with normalizeGiotto using scalefactor = 6000. Then, the functions createSpatialNetwork, binSpect and initHMRF_V2 were processed with k =16 for the brain sample, and k = 7 for the kidney sample. Annotation was extracted with the doHMRF_V2 function and visualized independently. GraphST is run with default parameters to obtain a 64-dimensional representation of cells. Then, Louvain is applied to cluster each sample by adjusting the resolution until a similar number of clusters as CCD is achieved. Seurat objects for each slice were created for both BASS and PRECAST, and default values were used for all parameters, together with the desired number of clusters. CCD for mouse embryo whole brain sample was ran with win_sizes = 150, sliding_steps = 50, cluster_algo = ‘spectral’ and n_clusters = 16, while for multi-sample adult dataset parameters were winsizes = 200, sliding_steps = 50, cluster_algo = ‘agglomerative’ (Hierarchical) and n_clusters = 16. All parameters were chosen to provide, on average, 30-40 cells per window, while keeping the communities smooth and coherent.

#### Evaluation metrics

We utilized two metrics to evaluate the performance of various algorithms in generating results: Scatter and Density BetWeen clusters (S-Dbw) [66] and SD validity index [58]. S-Dbw considers both cluster separation and cluster cohesion. It measures how well-separated clusters are from each other (good separation) while also considering how tightly the data points are grouped within each cluster (good cohesion). SD validity index combines the measures of average cluster scattering and total separation between clusters. These dual considerations make S-Dbw and SD more comprehensive metrics for this purpose than the silhouette score that measures how similar each data point in one cluster is to the data points in the neighboring clusters. The total benchmark result can be found in Supplementary Table 1, CCD provides lower S-Dbw and SD scores than other algorithms, confirming better cluster cohesion and groupin. Meanwhile, we compared the execution time and memory consumption of GSDI, SpaGCN, GraphST, PRECAST, BASS and CCD (Supplementary Table 2). The execution time of the CCD is notably faster compared to the GSDI and SpaGCN, demonstrating a speedup of at least 90 and 35 times, respectively. The peak memory consumption is affected by the dimensions of the input file, rendering CCD significantly more efficient due to its independence from gene expression matrices.

#### Comparison on mouse embryo brain sample, and region analysis

The mouse embryo brain (Extended Data Fig. 7a), a structurally well-explored sample, was used for comparison of spatial domain detection methods, and for further analysis of the biological significance of CCD communities. Extended Data Fig. 2b provides a comparison of spatial regions obtained by Stereopy’s CCD, Giotto’s SDI, SpaGCN, and GraphST, with the numbers of domains fixed. SpaGCN fails to provide domain integrity. Both GSDI and CCD detect layers in the dorsal pallium, as well as the thalamus. However, CCD provides smoother and more coherent regions (Supplementary Table 1), with the detection of several more separate communities. The cell communities detected by Stereopy’s CCD are composed of multiple neighboring cell types and correspond to functional tissue domains. To evaluate the CCD’s ability to infer biological function and structure, we analyzed separate regions and their correspondence with known functional and anatomical regions. Extended Data Fig. 2c-d displays the region and composure of two communities which show significant spatial matching with Hotspot [9] gene modules, and anatomical regions from Allen brain map [10]. The orange community represents a cell type-homogenous region, with 70% of dopaminergic neurons (Die GNeu) and 23% of midbrain glutamatergic neuroblasts (Mb Glu Neu) as main components, where other cell types appear in abundancies less than 4%. Although these cells can be found in other areas of the tissue (Extended Data Fig. 2c, second column), this region is defined by the specific mixture of cell types, that is, a specific tissue domain. This community is spatially matched with the Hotspot gene module, as well as with the anatomical region of dorsal tier of thalamus (Extended Data Fig. 2c, columns three and four). The brown community is heterogeneous and contains, on average, 30% forebrain GABAergic neuron cells (Fb Glu NeuB), 29% cortical intermediate progenitor cells (Corti prog), 13% of cortical or hippocampal glutamatergic neuron cells (CortiHippo Glu Neu) and 10% of cortical glutamatergic neuron cells (Corti Glu Neu) (Extended Data Fig. 2d, first and second column). This region is shown to coincide with the gene module obtained by Hotspot, and when comparing with Allen brain atlas annotation, it corresponds to the mantle zone of dorsal pallium (Extended Data Fig. 2d, third and fourth column). These results confirm the ability of CCD to extract biological information.

#### Comparison on multi-sample adult mouse brain sample

Three samples were processed separately by GSDI, GraphST and SpaGCN, while BASS, PRECAST and CCD employed their multi-sample approach (Extended Data Fig. 3b). SpaGCN manages to obtain anatomical regions with clear borders but provides an unstable number of domains for consecutive samples while using the same parameters. (Extended Data Fig. 3b). When comparing per sample, domains obtained by GSDI, PRECAST, BASS, GraphST and CCD are similar by constitution. However, multi-sample processing provides more coherent (Supplementary Table 1) and anatomically matching results with higher reliability of inter-sample domain matching. Selected samples have similar cell type shares (Extended Data Fig. 3c). Thus, the consistency of CCD’s communities throughout samples is confirmed with stable tissue-share communities in all samples (Extended Data Fig. 3d) together with lowest S-Dbw and SD scores (Supplementary Table 1). Execution of 3 slices of adult mouse brain CCD finishes in 214 seconds while consuming 25716 MB. It costs less in terms of execution time and memory compared to other tools (Supplementary Table 2).

#### Comparison on UMOD KI / WT sample

CCD, BASS and PRECAST provide joint analysis of both UMOD KI and WT samples, while Giotto’s SDI, GraphST and SpaGCN perform on each sample separately (Extended Data Fig. 4c). We compared the results generated by these algorithms, especially the medulla region according to the annotation obtained from the Marshall et al. paper. CCD provides domains of higher integrity and robustness compared to BASS, PRECAST, GSDI and SpaGCN, especially in the medulla region on which CCD identified almost the same region with the annotation from the original paper. GSDI, BASS, PRECAST and SpaGCN detected more than one region and even mixed regions in the medulla area, while CCD and GraphST detect regions consistent with Marshall et al paper. To further demonstrate the consistency of CCD regions, we calculated the marker genes for each of them. Marker genes show consistency of the gene expression and the cell community region in both UMOD KI and WT samples (Supplementary Fig. 6). CCD manages to process these two kidney samples in 31 seconds while consuming only 684 MB. It costs less in terms of execution time and memory compared to other tools (Supplementary Table 2).

### Temporal gene pattern identification (TGPI) algorithm

It is of interest that the expression level of a gene shows a certain pattern during certain biological processes. Among various kinds of gene patterns, up- or down-regulation is a common pattern. Here we developed a method to find up- or downregulated genes utilizing the serial t-test along time series and cell type trajectory. We use a one-tailed t-test to get the statistic score and p-value between adjacent time points. Both p-values of greater and less test will be calculated to represent up- and down-regulated genes, respectively. Then we provided two metrics to combine the p-value so that we can sort out the most up- or downregulated genes. The two p-value combination metrics include:

1. Fisher’s method. It is based on the hypothesis that the sum of the -2 logarithm of the p-values from k-independent experiments follows a chi-squared distribution with 2k degrees of freedom. Then a combined p-value is tested from the chi-squared test.

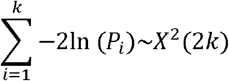
2. Plain False Discovery Rate (pFDR) method. We proposed it and integrated the metrics into TGPI based on the hypothesis that the alternative probability indicate the increasement of expression between adjacent time point and each time point is independent. Then, the false discovery rare without any correction is utilized to indicate the significance of serially up or down-regulation. The pFDR is calculated according to the following formula:

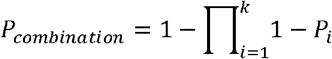

Except for serially up or downregulated genes, some genes show more complicated patterns during certain biological processes. To get all kinds of patterns automatically, we use fuzzy C means to cluster genes inspired by Mfuzz [33]. Stereopy considered both spatial and temporal expression features leading to more biological significant result. For spatial feature, Stereopy calculate pca based on rasterized expression on a certain bin size, and use first several principal components as spatial feature.

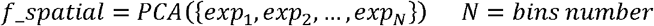

For temporal feature, Stereopy utilizes the result from serially up/downregulated genes as input. We use the serial greater p-value Pgi and serial less p-value Pli as features for each gene based on the following formula:

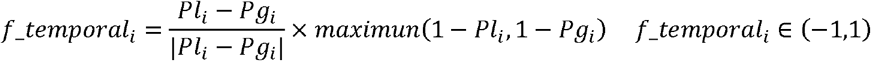

After calculation, the lower fi represents downregulated and the higher fi represents upregulated. In this way, we regard fi as the tendency of a gene between adjacent time points. Compared to Mfuzz which takes mean expression as input, Stereopy’s temporal feature will place more emphasis on the tendency rather than the original gene expression. To combine feature of both temporal and spatial, we concatenate the scaled spatial features with first N spatial feature and to temporal feature for each gene. A parameter alpha is also used to weight the effect of spatial features.

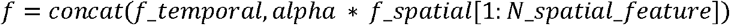

Finally. Fuzzy C means is used to cluster genes into groups. The main principle of fuzzy C means is to minimize J according to the following equation:

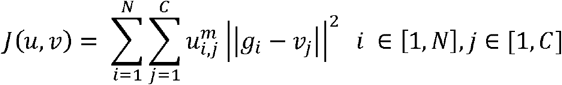

In this equation, fi represent the feature combined both temporal and spatial, vj belongs to the center of each cluster. m is the fuzziness and equals to 2 by default. 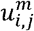 is the membership of i In this equation, fi represent the feature combined both temporal and spatial, vj belongs to the gene in j cluster it subject to:

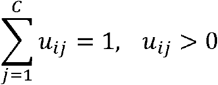

With the help of spatial feature, Stereopy’s TGPI can further distinguish gene clusters with similar temporal expression but spatially differential expressed, which makes the result more biologically significant (Extended Data Fig.6 and Supplementary Fig. 11).

### Benchmark of temporal gene pattern identification algorithm

The evaluation of Stereopy’s TGPI contains two important modules: 1) the p-value combination statistic metric ‘pFDR’ proposed in TGPI to find the up-/down-regulated genes. 2) the whole temporal gene pattern detection algorithm ‘TGPI’ in spatial-resolved temporal datasets.

#### P-value combination statistic metrics evaluation

We benchmarked our proposed p-value combination metric ‘pFDR’ with fisher’s method on temporal mouse forebrain datasets. We made three comparisons with respect to three cell types including dorsal forebrain, forebrain neuronal intermediate progenitor, and forebrain cortical glutamatergic in these datasets (Extended Data Fig 5 and Supplementary Fig. 9-10). According to the occurrence of three cell types, three comparisons contain 7, 5, and 3 time points respectively. In each comparison, we used ‘pFDR’ and fisher’s method to detect the continuous up-/regulated genes and visualized the corresponding gene expressions. It is obvious that ‘pFDR’ can better find real continuously up-/down-regulated genes which has a stable tendency of rise or fail gene expression along with the time series in all these comparisons.

#### Temporal gene pattern detection algorithm evaluation

We first tested the effect of spatial features. N top spatial features range from 3 to 6 are tested. Taking *Foxg1, Hes5*, and *Mab21l2* as examples, we tested the 4 nearest neighbors (NN) of these genes according to Euclidean distance based on N top spatial features. (Extended Data Fig. 6). From the result we observed that a similar spatial expression pattern is detected in each 4NN gene. The higher the N spatial feature is, the more similar spatial expression patterns can be observed (Extended Data Fig 6). Additionally, as the N spatial feature reaches 5, the 4NN genes tend to be constant. Since the N spatial feature can reflect the spatial expression feature, we tested its influence on TGPI (Supplementary Fig. 11a). The result indicated that the increment of N spatial feature resulted in higher consistency of genes in a temporal pattern to some extent (blue box). Moreover, with the help of spatial features, TGPI can distinguish genes with similar temporal patterns. For example, Cluster 2 and Cluster 8 of TGPI with N spatial feature equal to 3 are similar in temporal expression pattern and divergence in spatial expression pattern (Supplementary Fig. 11b-c).

To evaluate TGPI’s performance on real datasets, we compared the TGPI algorithm with another time series gene pattern method called Mfuzz [33]. The Stereo-seq mouse embryo brain data from E9.5 to E16.5, which is the subset of mouse embryo dataset with annotation as ‘Brain’, is used to evaluate the performance of TGPI [3]. Genes were clustered into 8 clusters for both Stereopy and Mfuzz. To evaluate the performance of the gene pattern results, we calculate the Pearson’s correlation of gene expression with the pseudotime and ANOVA test among time points. The F-score of ANOVA test is used to reflect the divergence between time points. If a certain gene is more related to time points, the F score will be higher. We calculated top 100 genes of each TGPI cluster ordered by weights of clustering results for both Stereopy and Mfuzz. The results show that most TGPI clusters exhibit not only a higher F score in the ANOVA test among time points but also higher Pearson’s correlation with pseudotime, which means genes within the gene pattern identified by TGPI are more related to both time point and pseudotime (Extended Data Fig. 7a). Moreover, from the GO enrichment results, we concluded that TGPI is more capable of grouping genes with the same expression pattern and functions (Extended Data Fig. 7b-c). We conducted GO enrichment analysis on top 20 genes of each cluster for both TGPI and Mfuzz. As a result, 7 gene pattern clusters of TGPI enriched GO terms while only 3 clusters enriched GO terms for Mfuzz with the same p value cut off (p=0.05). Meanwhile, TGPI’s genes are more related to neuron development. For example, Cluster 5 of TGPI’s result has a similar tendency to Cluster 8 of Mfuzz results, both of which enriched GO terms related to synapse organization and assemble. However, the gene count of GO in TGPI reached 5 while the gene counts of Mfuzz’s cluster 8 reached 3. Additionally, the TGPI enriched more GO terms related to neuron development (sensory perception in Cluster 1, learning or memory in cluster 8) and mitotic (mitotic nuclear division in cluster 7).

### 3D cell-niche regulatory network prediction algorithm

Stereopy-NicheReg3D starts with the cell-niche communication prediction. To ensure the accuracy and specificity of this juxtacrine signaling model, we extract cells bordering their niches and statistically calculate their CCC activity scores of L-R pairs under the assumption that intercellular L-R communications routinely exist among closely neighboring cells. The niche is defined as all the neighboring cells from other types whose Euclidean distance to any of the cells from the center type is less than a pre-defined radius *r*. Let *S*_*k*_ denote the set of all cells of type *k,c*_*i*_ represent cell *i*, d(·, ·)represent Euclidean distance between any two cells, then the niche can be formulated as:

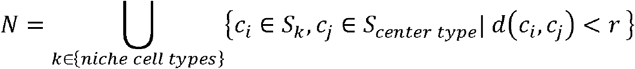

Next, we perform a label permutation-based statistical CCC analysis to generate significant cell-niche L-R pairs by incorporating both L-R gene co-expression and 3D location of the cells. In brief, we collect potential L-R pairs and construct a customized Liana consensus database [43] (https://github.com/saezlab/liana-py/tree/main/liana/resource/omni_resource.csv). We then follow a similar approach reported by CellPhoneDB [45] to compute the average expression level of the ligand in the sender cells and that of the receptor in the receiver cells in the cell-niche boundary. The communication score is defined as the mean value of the average L-R expression within a 3D niche:

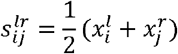

where 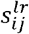 is the communication score for ligand *l* in cell type *i* and receptor *r* in cell type *j*,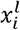 is the average expression level of ligand *l* in sender cell type *i*,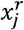 is the average expression level of receptor *r* in receiver cell type *j*.

The significance of the communication scores is evaluated through random shuffling of the cell type labels of cells in the niche multiple times,*m*. The p-value is defined as the number of random shuffles that reach a score higher than the true score:

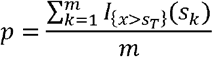

where *s*_*T*_ is the true communication score, *s*_*k*_ is the calculated communication score at the *k*th shuffle,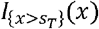 is the indicator function which equals to 1 if *x*>*s*_*T*_, and 0 otherwise. Typically, a p-value smaller than 0.05 suggests that the corresponding L-R interaction is statistically significant.

To comprehensively demonstrate the possible regulation mechanism, we eventually connect significant L-R interactions detected in the cell-niche communication analysis with the TF-centered regulons identified by the SpaGRN[67] analysis based on the integrated weighted ligand-signaling network from Nichenet-v2 [59] (https://zenodo.org/record/7074291/files/weighted_networks_nsga2r_final_mouse.rds). This database contains 3,865,137 rows, each of which represents a pair of directed signaling interactions with a specific weight prioritized using 57 data sources. We convert the whole network data into a weighted directed graph *G* = ⟨*V,E,W*⟩. For a given receptor and TF, we search for the shortest path between the two nodes and consider it as the potential signaling path between them. The distance of each graph edge is defined as the reciprocal of its weight:

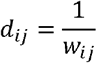

Where *w*_*ij*_ is the weight of edge *e*_*ij*_ connecting node *i* and *j*.

### Benchmark of cell-niche communication prediction algorithm

To demonstrate the algorithm efficiency, we systematically compared the general features of the Stereopy-NicheReg3D module with CellPhoneDB [45] and NICHES [50] to the same mouse heart dataset (Supplementary Table 6). We slightly modified two software tools to enable them to analyze the 3D SRT data. For the CellPhoneDB implementation, the spatial relationship of VCM and other cell clusters was initially provided and the default parameters were used to obtain VCM-significant L-R pairs. NICHES was adopted to obtain single-cell-resolution interaction results. We then integrated the expression of L-R pairs coming from each niche component and landing on the VCM cells by summing the L-R expression, and identified the cell type-specific L-R pairs using the Seurat FindAllMarkers function.

We benchmarked the performance of this module and the other two tools on the same Linux system with Intel Core Processor (Broadwell, IBRS) of 30 threads and 512 GB memory. Both Stereopy and NIHCES enable the investigation in sender–receiver single cells, which is usually computationally prohibitive for CCC analysis thanks to Stereopy’s niche extraction and NICHES’s subsampling strategies. These strategies also accelerate the computation compared to the whole cluster-based CellPhoneDB (Extended Data Fig. 9). However, subsampling might preclude a complete view of CCC structure and risk obscuring significant L-R pairs. As a result, in terms of the number of specific CCC interactions, Stereopy obtains the most specific L-R pairs in all VCM-niche cases except ACM-VCM, almost covering those derived by other tools (Extended Data Fig. 9).

## Data availability

The processed datasets have been deposited in published papers. **Slide-seq2 datasets:** mouse kidney datasets are downloaded from [28], of which “P uck 191204 22.h5ad” and “P uck 191204 15.h5ad” are used as BTBR WT and *ob/ob* sample respectively and “P uck 191223 19.h5ad” and “P uck 200104 07.h5ad” are used as WT and UMOD KI sample respectively. **Stereo-seq datasets:** a sample of 12 weeks adult mouse brain, mouse embryo SRT samples from E9.5 to E16.5, and entire 3D mouse embryonic heart datasets are downloaded from StomicsDB MOSTA [68]. Three adjacent samples of coronal mouse brain are downloaded from Spatial-ID [64].

## Code availability

Stereopy is a pip installable Python package and is available at the following GitHub repository: https://github.com/STOmics/Stereopy, with documentation at: https://stereopy.readthedocs.io/en/latest/. All the code to reproduce the result of the analysis can be found at the following GitHub repository: https://github.com/STOmics/Stereopy/tree/main/docs/source/Tutorials.

## Acknowledgments

This work is part of the “SpatioTemporal Omics Consortium” (STOC) paper package. A list of STOC members is available at: http://sto-consortium.org. We acknowledge the Stomics Cloud platform (https://cloud.stomics.tech/) to provide convenient ways for analyzing spatial omics datasets. We would like to thank Kai Han, Dantong Wang, Ping Qiu, Yunjia Zhang, Haohao Deng, Chang Shi, Junfu Guo for their help. We acknowledge the CNGB Nucleotide Sequence Archive (CNSA) [69] of China National GeneBank DataBase (CNGBdb) [70] for maintaining the MOSTA database.

## Author information

These authors contributed equally: Shuangsang Fang, Mengyang Xu, Lei Cao, Xiaobin Liu, Vladimir Kovacevic, Liwei Tan, Zhiyuan Yuan

These authors jointly supervised this work: Xun Xu, Yuxiang Li, Ao Chen, Yong Zhang, Guangyi Fan, Yi Zhao

## Authors and Affiliations

**BGI Research, Beijing, China**.

Shuangsang Fang, Lei Cao, Tianyi Xia, Luni Hu

**BGI Research, Shenzhen, China**.

Vladimir Kovacevic, Liwei Tan, Longyu Guo, Marija Bezulj, Junhou Hui, Chao Liu, Li’ang Lin, Zhenbin Wen, Bojana Josic, Nikola Milicevic, Qin Lu, Yumei Li, Leying Wang, Chao Zhang, Qiang Kang, Fengzhen Chen, Junhua Li, Mei Li, Shengkang Li, Yong Zhang, Ao Chen, Yuxiang Li

**BGI Research, Qingdao, China**.

Mengyang Xu, Xiaobin Liu, Yao Li, Lidong Guo, Guangyi Fan

**Institute of Science and Technology for Brain-Inspired Intelligence, MOE Key Laboratory of Computational Neuroscience and Brain-Inspired Intelligence, Fudan University, Shanghai, China**

Zhiyuan Yuan

**BGI Research, Hangzhou, China**.

Mengnan Cheng

**Beijing Key Laboratory of Mobile Computing and Pervasive Device, Institute of Computing Technology, Chinese Academy of Sciences, Beijing**, **China**

Yi Zhao

**BGI Research, Wuhan, China**.

Xun Xu

## Contributions

X.X., Y.X.L, A.C. and Y.Z. conceptualized the study. S.S.F, M.Y.X, L.C., X.B.L. V.K. L.W.T, M.B. and L.D. G were responsible for algorithm development and implementation. Y.L., T.Y.X., B.J. and N.M. analyzed the data. C.L., M.N, Q.L. and Y.M.L. contributed to the data preprocessing. L.Y.G., J.H.X., L.A.L, Z.B.W., L.Y.W. and F.Z.C. contributed to cleaning up the code and maintaining the website of Stereopy. Y.Z.Y. and L.N.H, J.H.L. M.L, C.Z., Q.K., S.K.L contributed key ideas and advice. X.X., Y.X.L, A.C. Y.Z., G.Y.F. and Y.Z. supervised the study.

## Extended Data Figures

**Extended Data Fig. 1.**
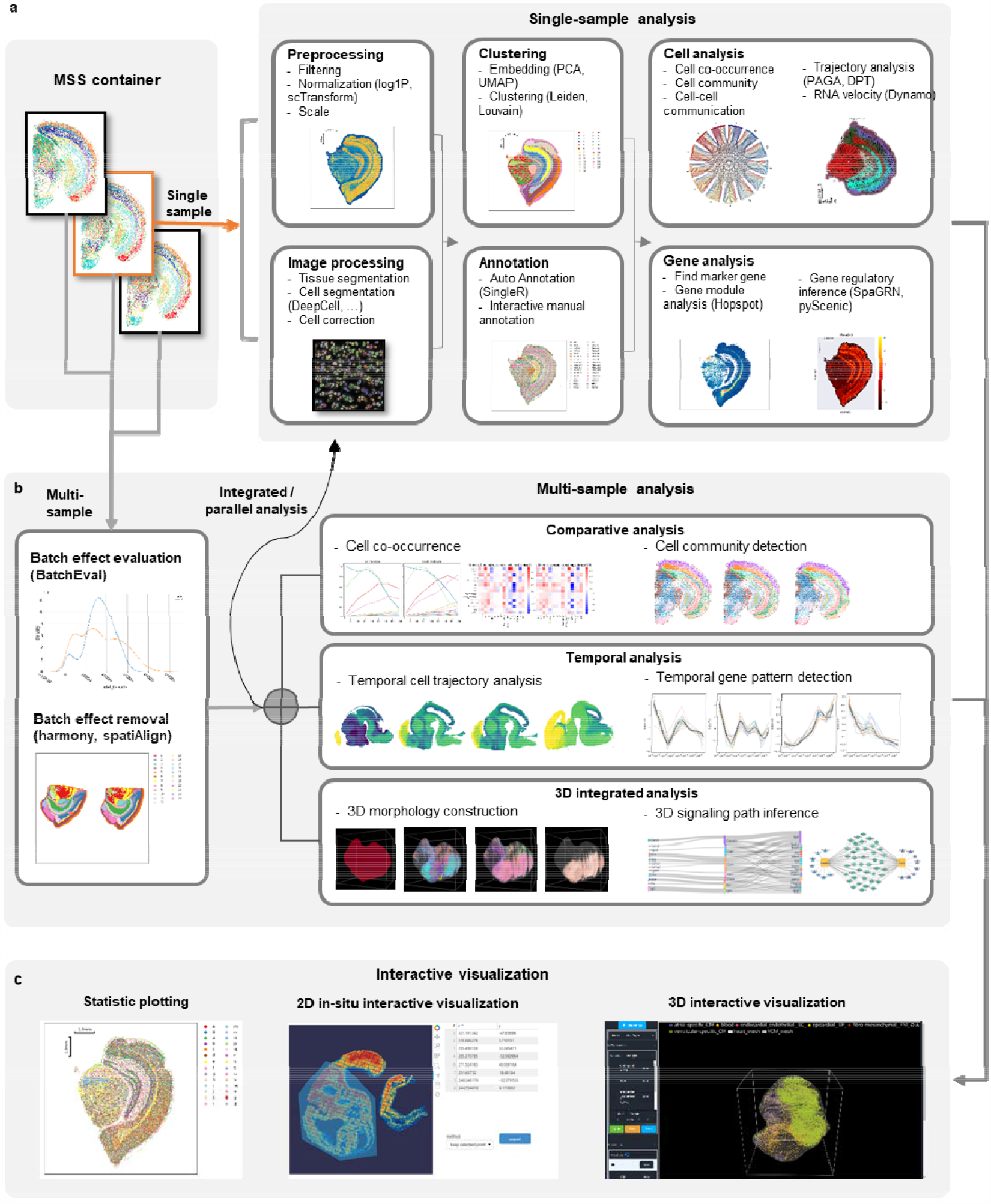
Overview of Stereopy functions. **a**, Analysis functions for single sample, including preprocessing for spatial gene expression and images, clustering and annotation, cell-level and gene-level analysis modules. **b**, Analysis functions for multi-sample joint analysis, including batch effect evaluation and removal processing, comparative analysis, temporal analysis and 3D integrated analysis. **c**, Interactive visualization, including statistic plotting, 2D in-situ interactive visualization and 3D interactive visualization.

**Extended Data Fig. 2.**
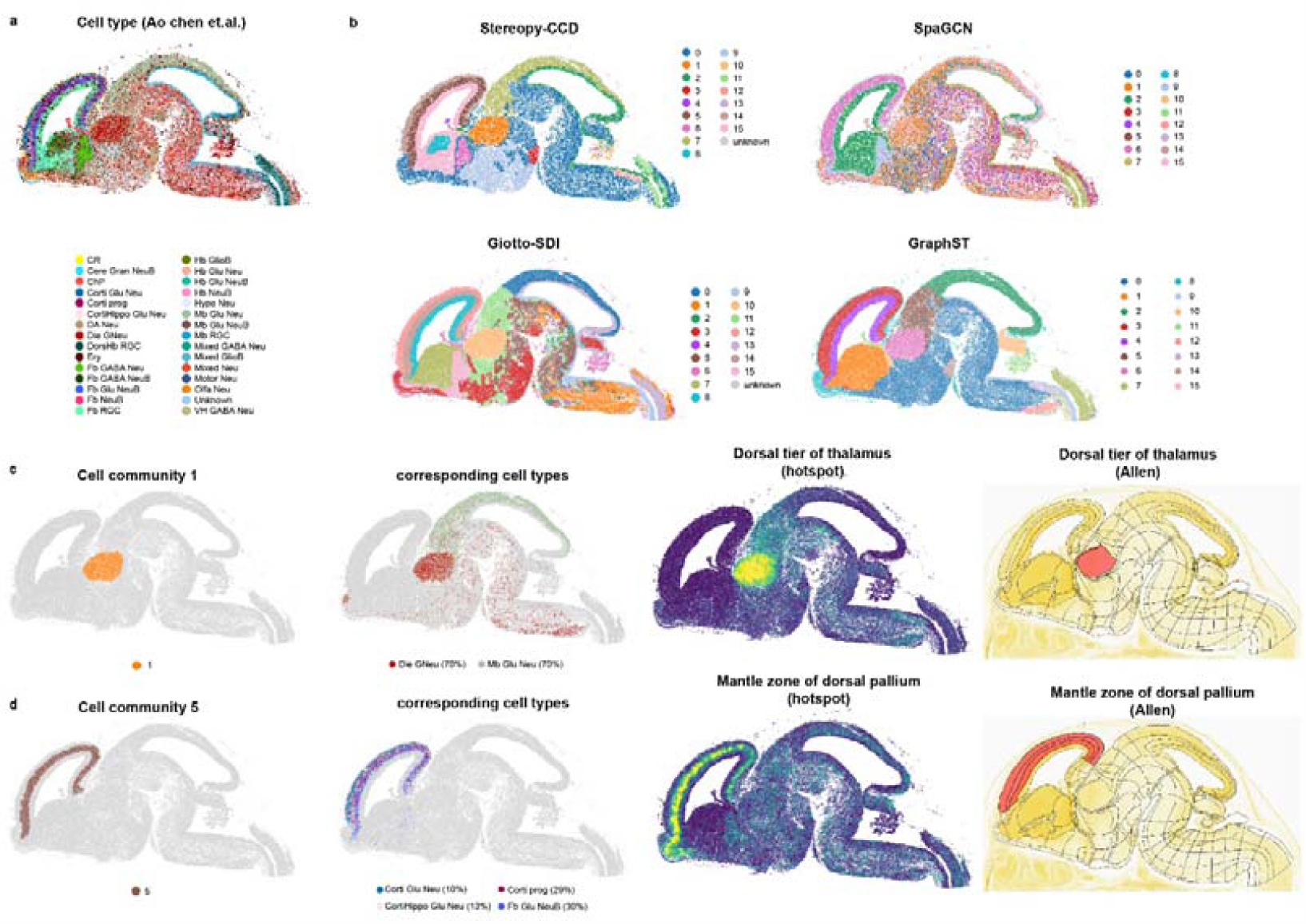
Cell community detection and comparisons on Stereo-seq mouse embryo whole-brain sample. **a**, Cell type annotation of the mouse embryonic brain obtained from MOSTA database. **b**, Stereopy-CCD, Giotto-SDI, SpaGCN, and GraphST results. **c-d**, Comparative display of detected cell communities and their corresponding functional and anatomical domains. Left to right: area of the community, tissue distribution of cell types comprising the community with community cell types and percentages shown in the legend, Hotspot domain, and anatomical region from the Allen mouse brain atlas corresponding to the community region. **c**, Community matching the dorsal tier of thalamus. **d**, Community matching the mantle zone or dorsal pallium.

**Extended Data Fig. 3.**
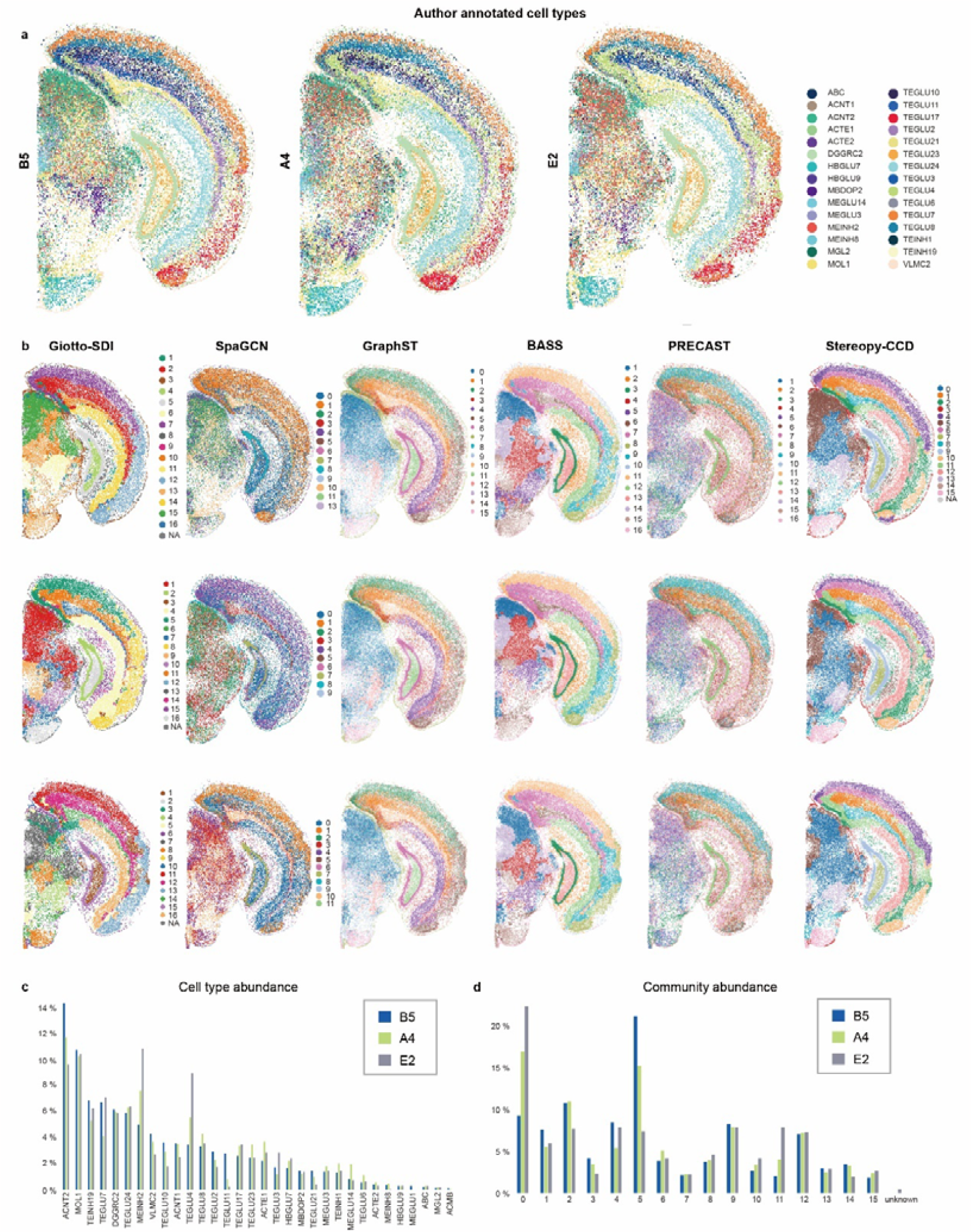
Cell community detection and comparisons on Stereo-seq multi-sample adult mouse brain sample. **a**, Cell type annotation of three adult mouse brains. **b**, Domain detected by Giotto-SDI and SpaGCN, for each sample as well as domain / cell community detected by GraphST, BASS, PRECST and Stereopy-CCD for multi-sample joint processing. **c**, Bar plot of per sample cell type abundance by Stereopy-CCD. **d**, Bar plot of per sample cell community abundance by Stereopy-CCD.

**Extended Data Fig. 4.**
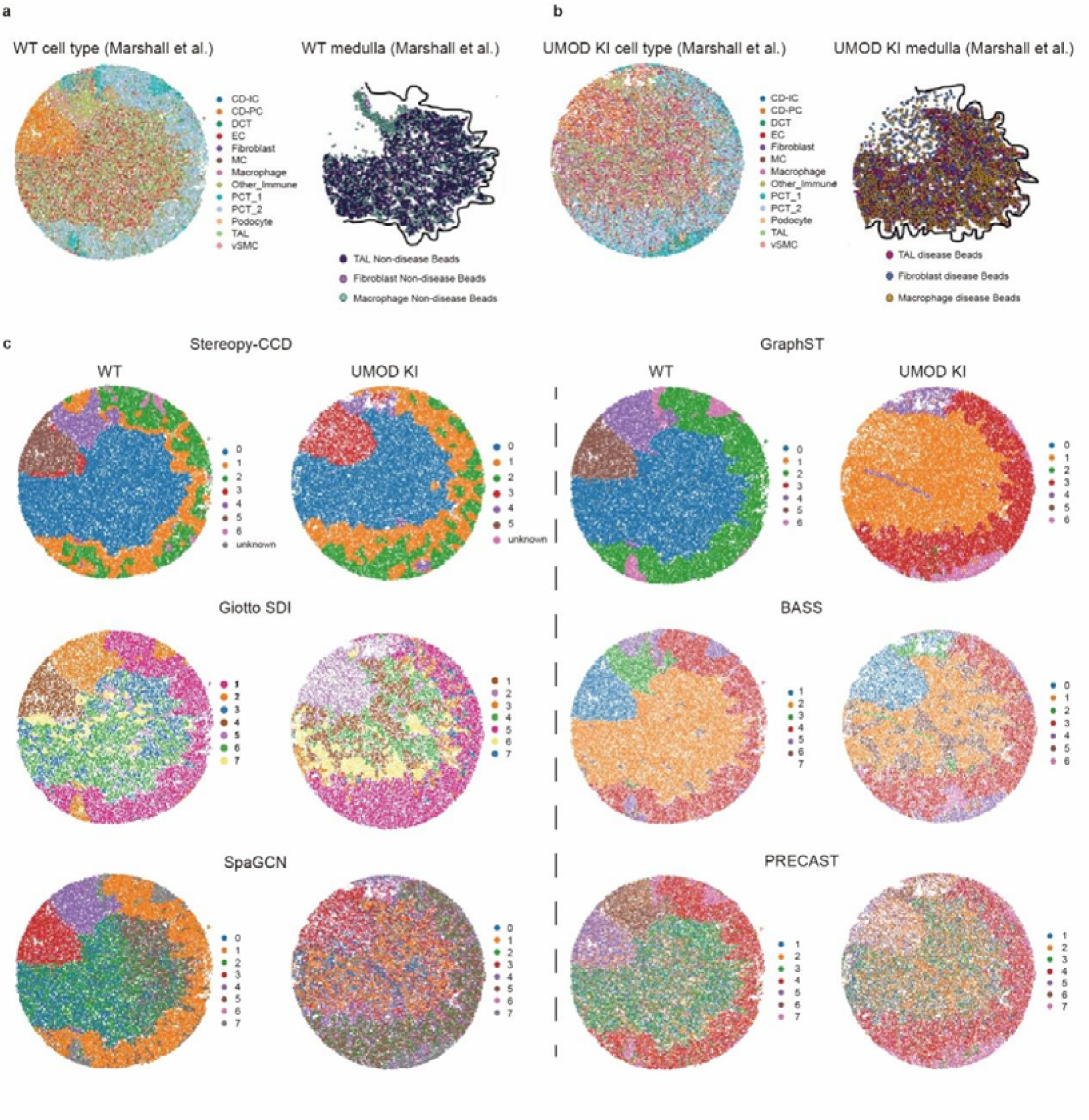
Comparison of Stereopy-CCD, Giotto-SDI, SpaGCN, GraphST and STAGATE on mouse kidney samples. **a**, Author annotated cell type and author annotated medulla region of WT kidney sample. **b**, Author annotated cell type and author annotated medulla region of UMOD KI kidney sample. **c**, Domain detected by Giotto-SDI and SpaGCN for each sample, as well as domain / cell community detected by GraphST, BASS and PRECAST and Stereopy-CCD for multi-sample joint processing.

**Extended Data Fig. 5.**
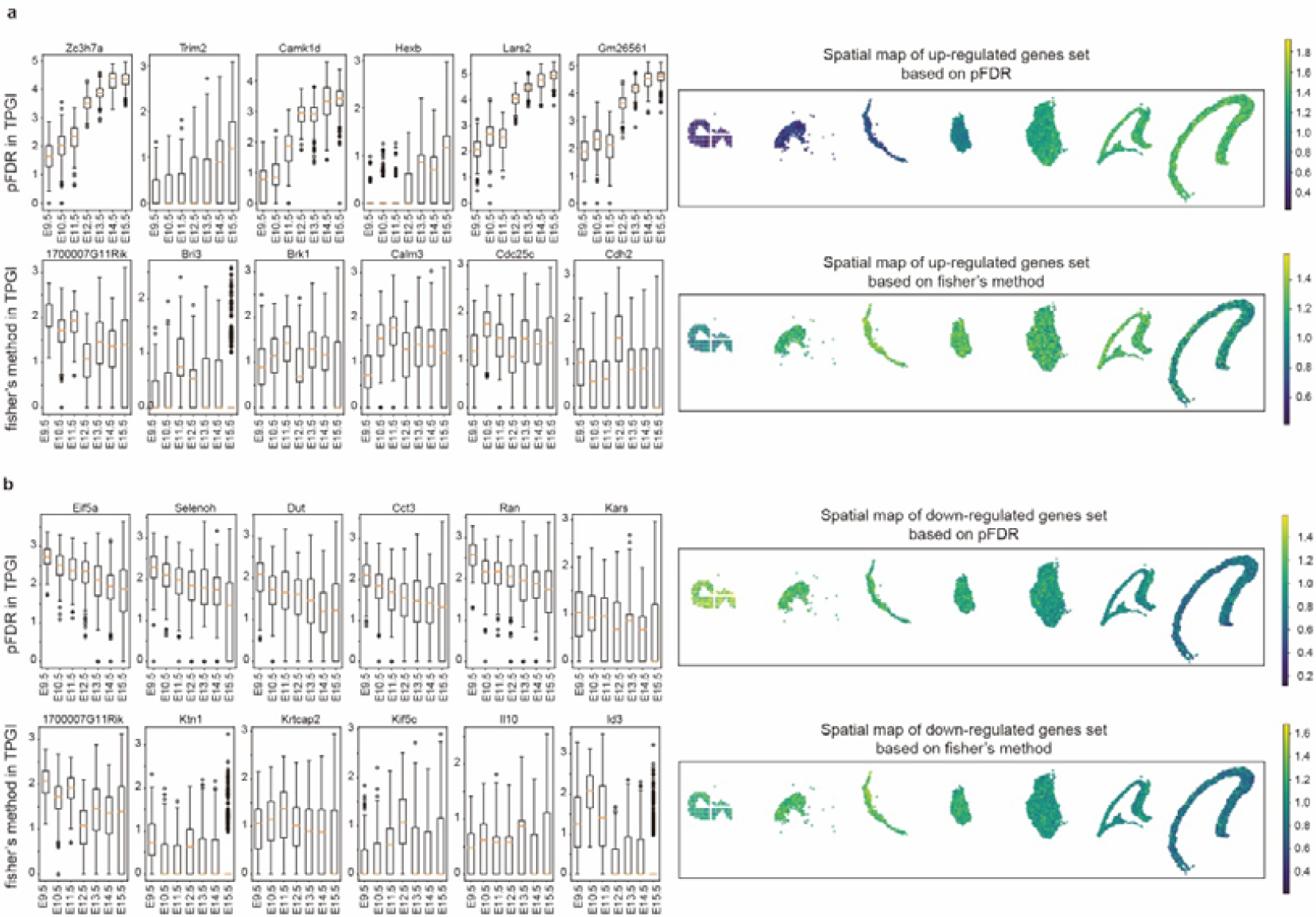
Up and down regulated genes identified by Stereopy-TGPI for temporal dorsal forebrain. **a**, Left: boxplot of up regulated genes based on different p value combination method; right: spatial map of gene set of mean expression of top 20 up regulated genes. The result is calculated based on pFDR in Stereopy-TGPI (Top), and fisher’s method (Bottom), respectively. **b**, Left: boxplot of down regulated genes based on different p value combination method; right: spatial map of gene set of mean expression of top 20 down regulated genes. The result is calculated based on pFDR in Stereopy-TGPI (Top), and fisher’s method (Bottom), respectively.

**Extended Data Fig. 6.**
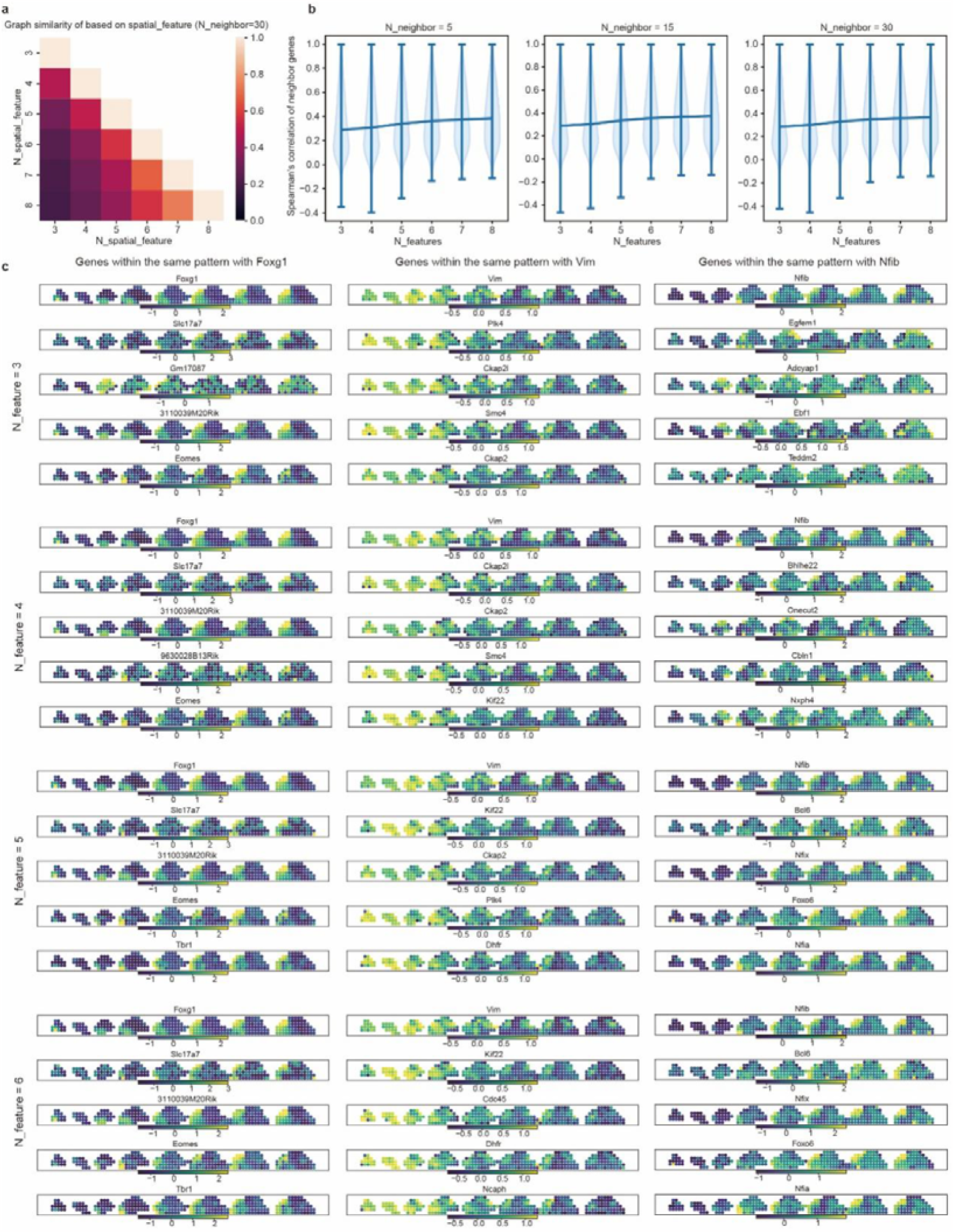
The influence of spatial features on the detection of temporal gene pattern. Each column shows the spatial maps of genes which have minimal distances with the certain gene calculated based on different number of spatial features range from 3 to 6. The spatial map is present on rasterized temporal mouse brain. The certain genes are *Foxg1, Vim*, and *Nfib* from left to right respectively. The distance is calculate based on 3,4,5,6 top spatial features from up to down, respectively. The red box indicates that when the top spatial features are increased to 5, the results become comparable to those obtained with a larger number of top spatial features.

**Extended Data Fig. 7.**
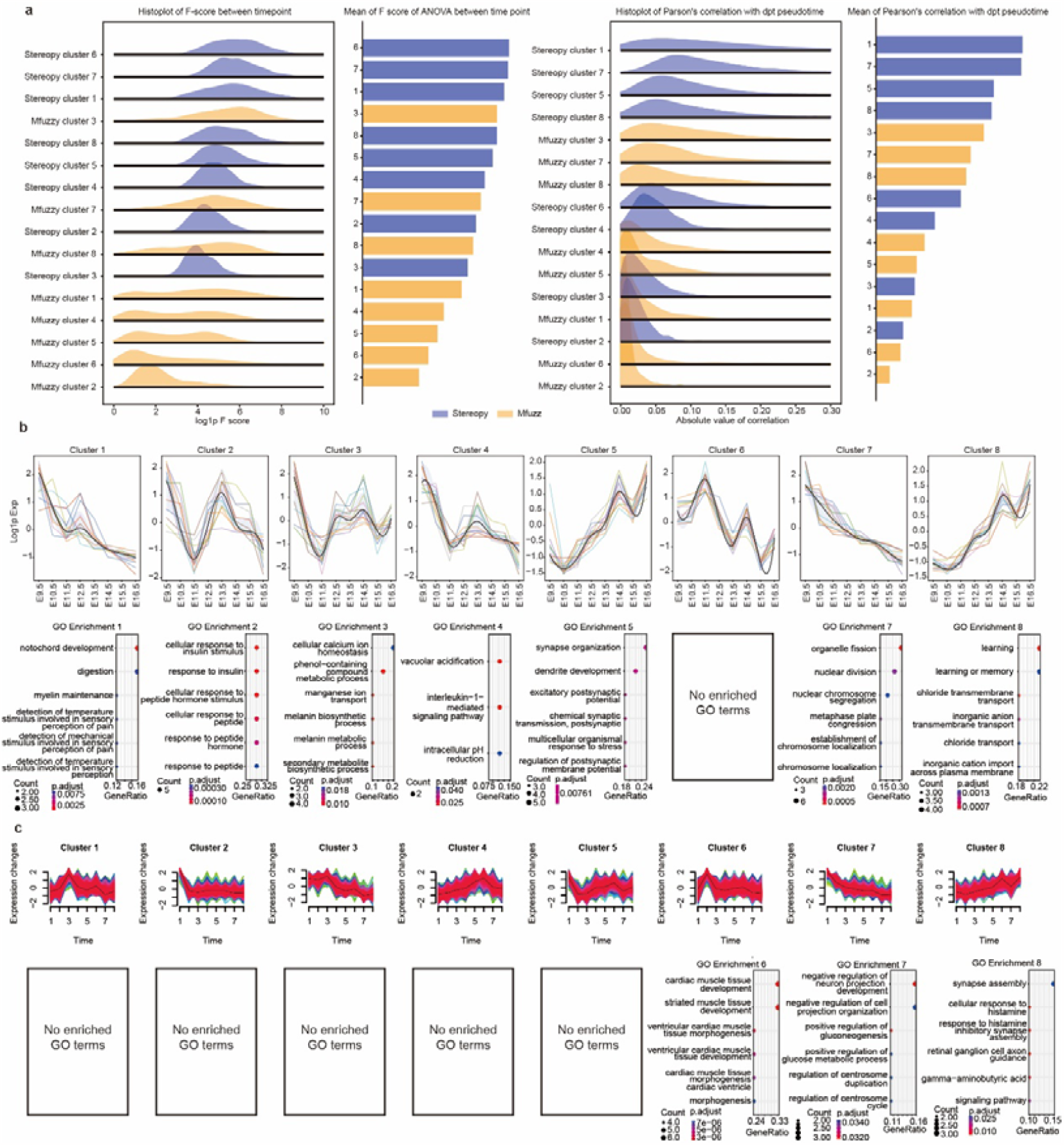
Comparison of TGPI in Stereopy and Mfuzz for the temporal gene pattern clusters result. **a**, The F-score among time point and correlation with pseudotime of top 1000 gene of each cluster of Stereopy-TGPI and Mfuzz. Blue and yellow represent Stereopy and Mfuzz, respectively. **b**, Stereopy temporal gene pattern for mouse brain along time series and corresponding GO enrichments for each temporal gene pattern cluster. **c**, Mfuzz temporal gene pattern for mouse brain along time series and corresponding GO enrichments for each temporal gene pattern cluster.

**Extended Data Fig. 8.**
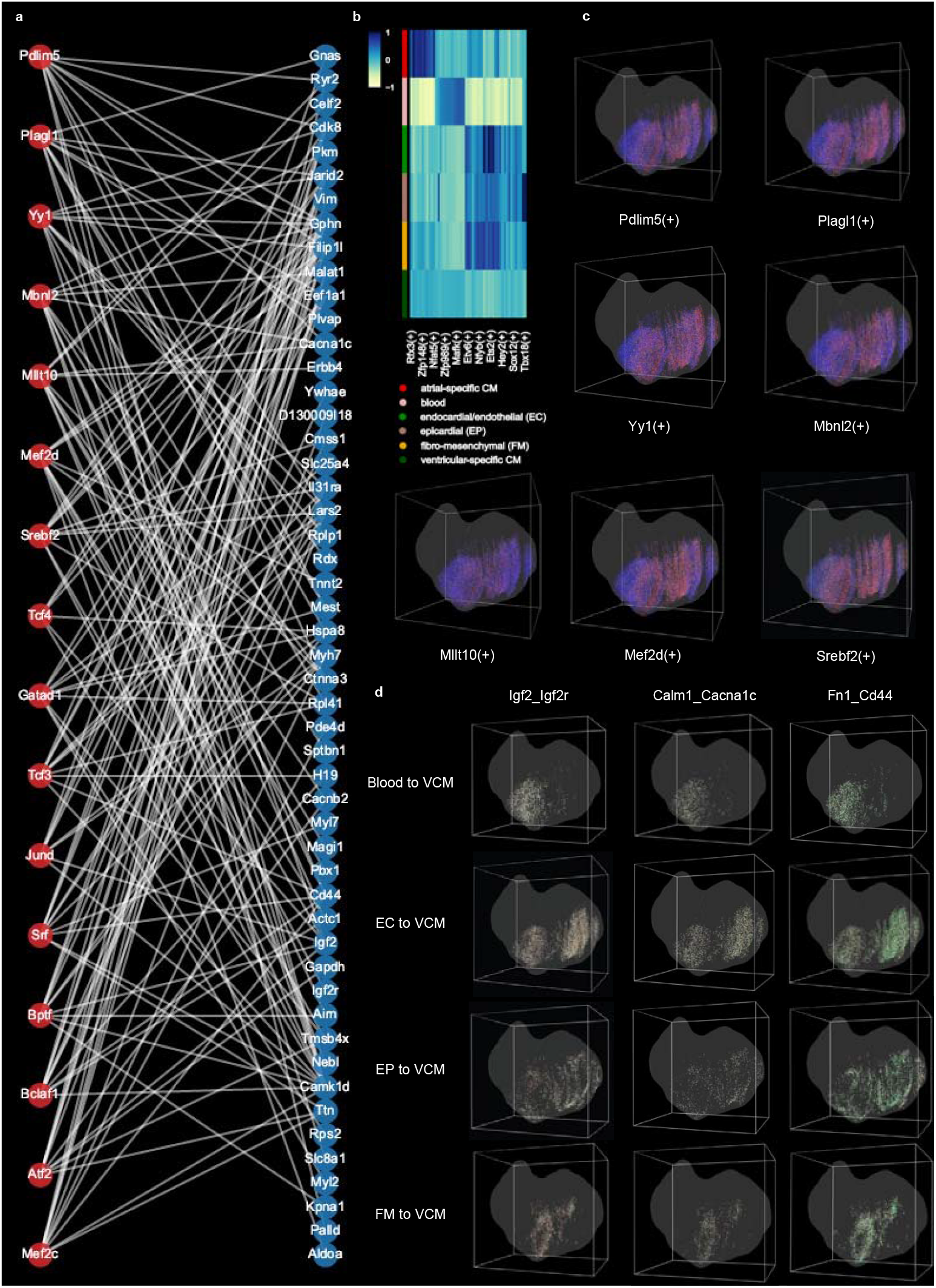
3D visualization of selected regulatory regulons and selected cell-cell communications in four VCM niches. **a**, TF and target genes in 16 representative VCM-specific regulons that are potentially regulated by VCM-niche L-R pairs. **b**, Heatmap disclosing top cell-type-specific regulons detected by NicheReg3D. **c**, 3D distribution of selected regulons in (a). **d**, 3D visualization of *Igf2_Igf2r, Calm1_Cacna1c* and *Fn1_Cd44* from blood, EC, EP, FM to VCM cells in each corresponding niche.

**Extended Data Fig. 9.**
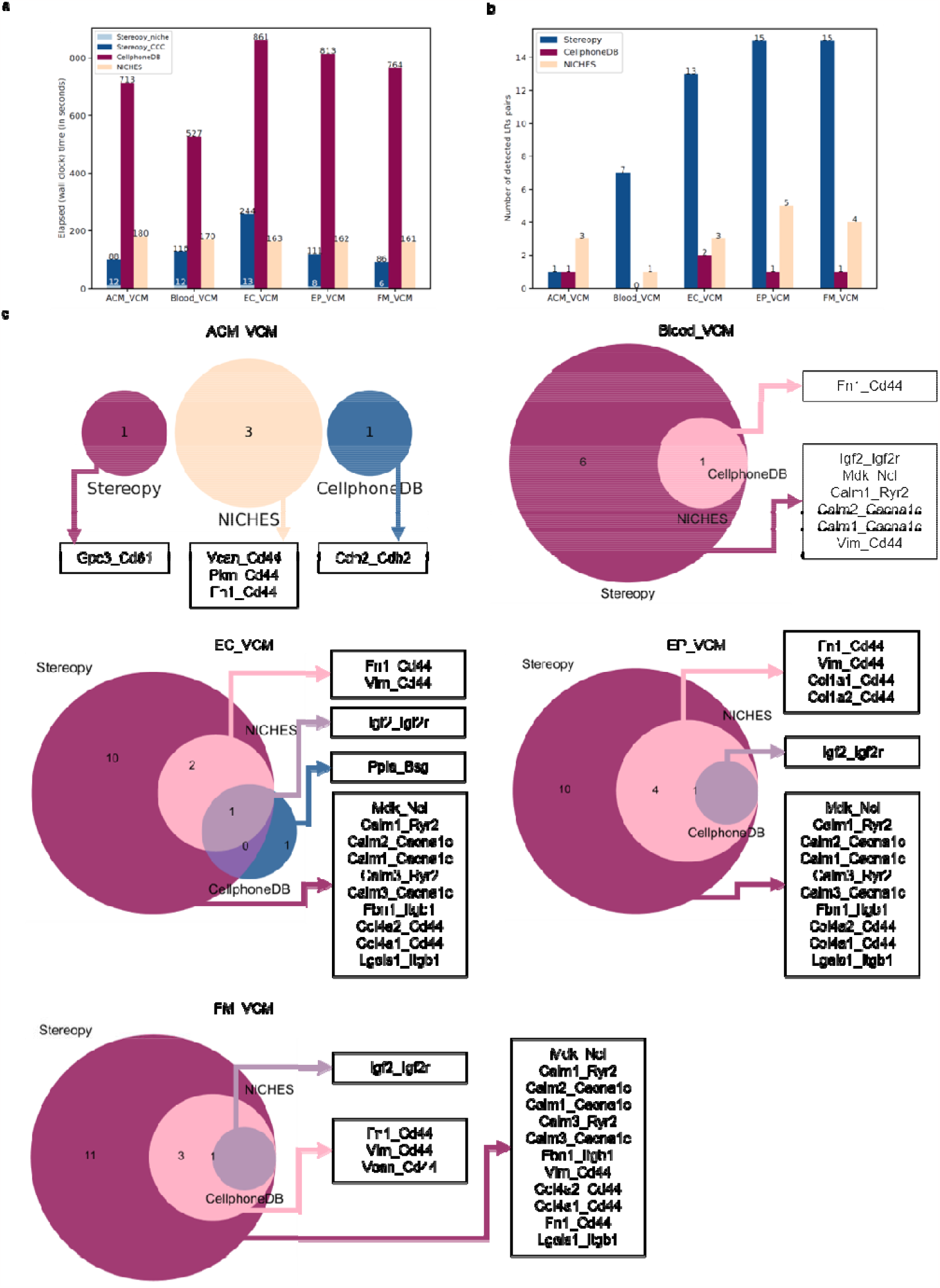
CCC comparison of Stereopy, CellphoneDB, and NICHES. **a**, Runtime of CCC analysis for five VCM-niche datasets using different CCC tools. **b**, Number of significant L-R pairs obtained by different CCC tools. **c**, Venn diagrams showing that the Stereopy CCC module detects more reliable L-R pairs, which almost cover those detected by the other two tools.

**Extended Data Fig. 10.**
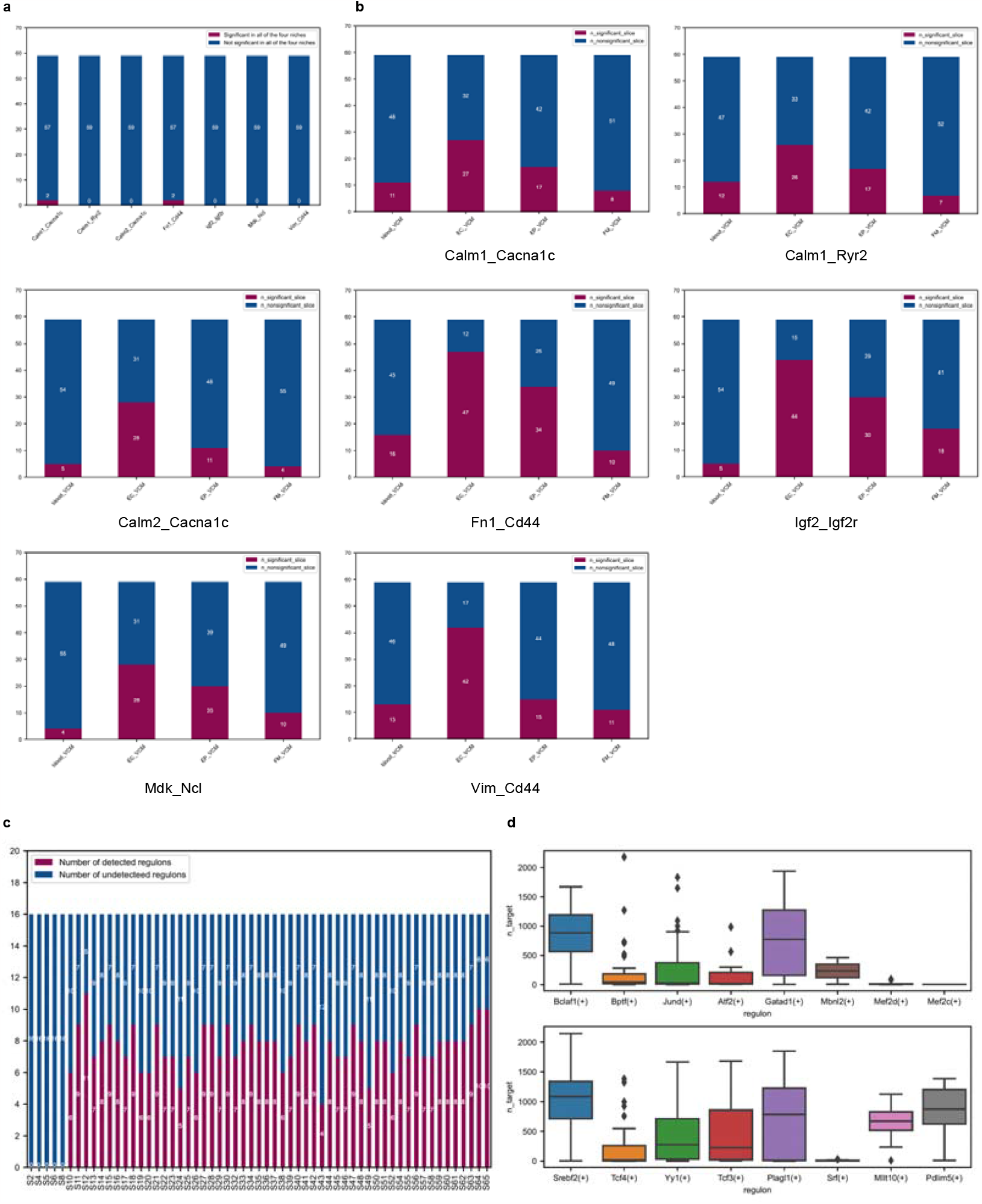
Comparison of 3D joint analysis and 2D analyses. **a**, Stacked chart showing the number of 2D samples that can detect significant L-R pairs demonstrating the completeness of CCC results from the 3D joint analysis. **b**, Stacked chart showing number of 2D samples that can detect each of the seven L-R pairs in (a). **c**, Stacked chart showing number of regulons among the 16 regulons potentially influenced by CCC in the VCM niche across the 2D slices. **d**, Boxplot illustrating the distribution of the number of targets in each regulon across the 2D slices.

## Supplementary Figures

See “Supplementary Figure”

## Supplementary Tables

See “Supplementary Tables”

## Supplementary Note

See “Supplementary Note”

