## Supplementary Figures for "Stereopy: modeling comparative and spatiotemporal cellular heterogeneity via multi-sample spatial transcriptomics"

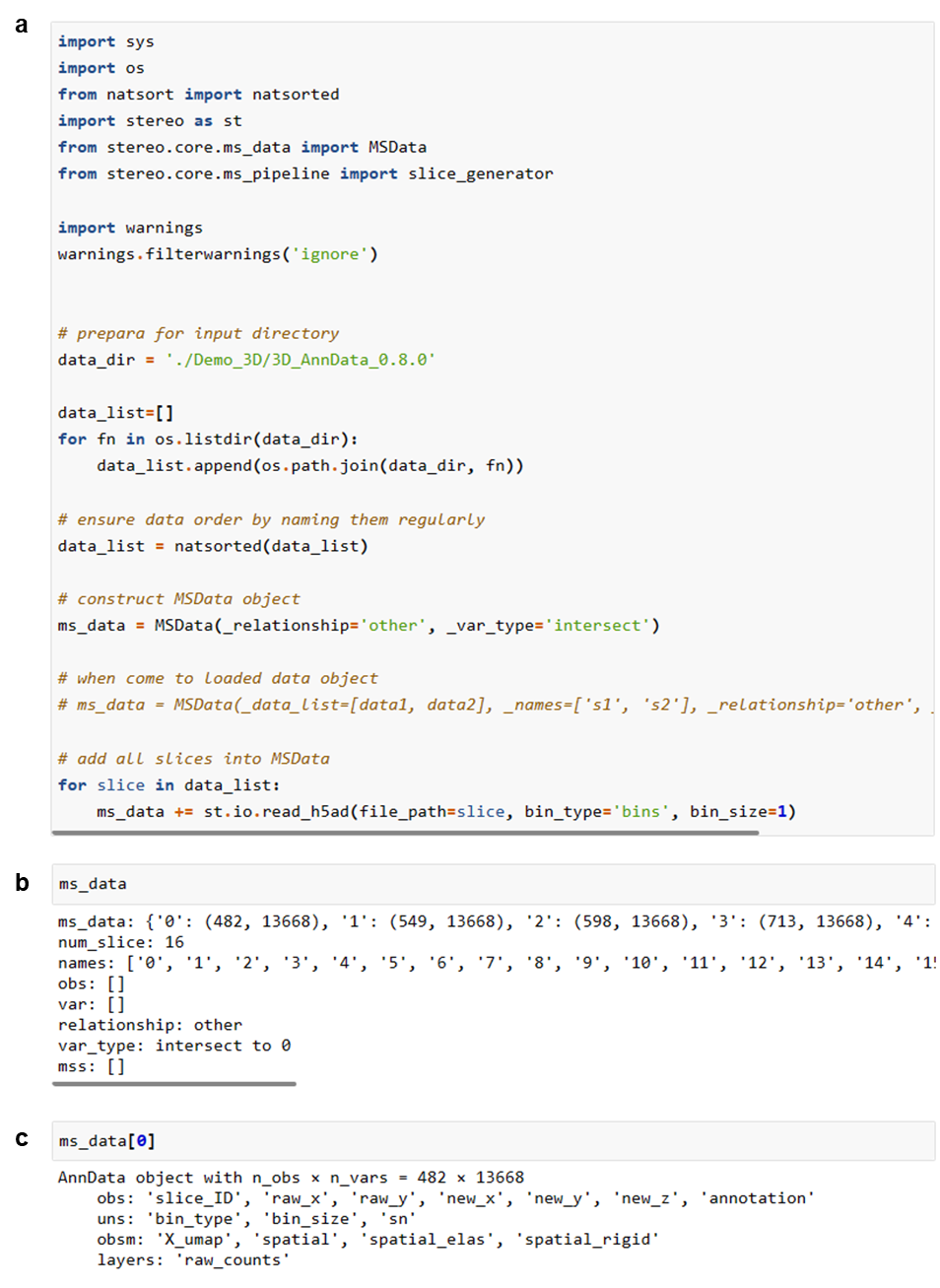

**Supplementary Fig. 1 An example of MsData in Stereopy.**

**a**, An example of loading multiple samples into MsData container in Stereopy. **b**, Display of MsData Data Content. **c,** Access one of the samples via the MsData handler.

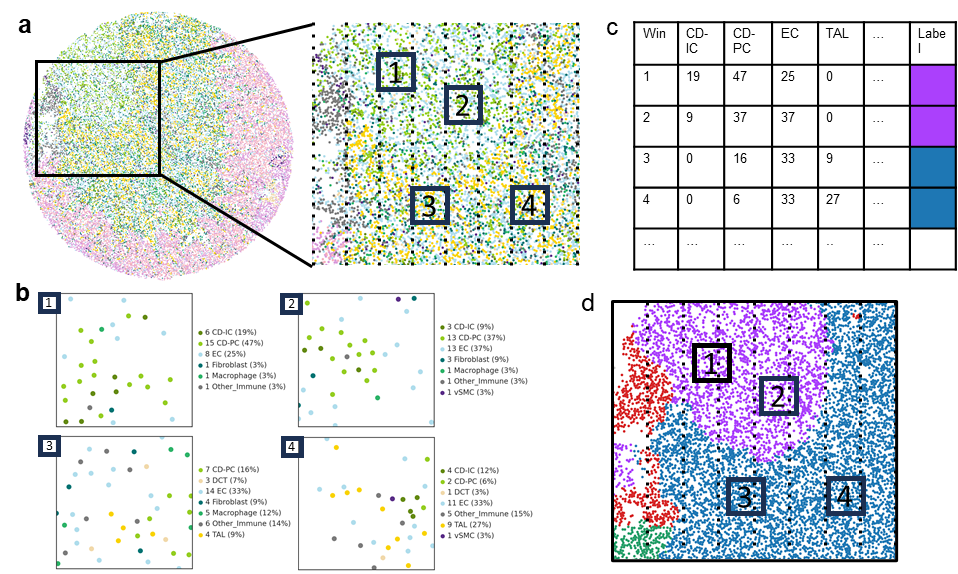

**Supplementary Fig. 2 High-level diagram of Cell community Detection algorithm.**

**a,** Sample of the tissue is divided into equally sized windows. **b,** Different cell types are present in each of the windows. **c,** The percentage of each cell type in the window is calculated and packed into the table containing windows as rows and cell types as columns suitable to be an input for the unsupervised clustering algorithm, such as Leiden or Spectral clustering. **d,** Each cell inside the window will obtain the label of the window(s) covering it.

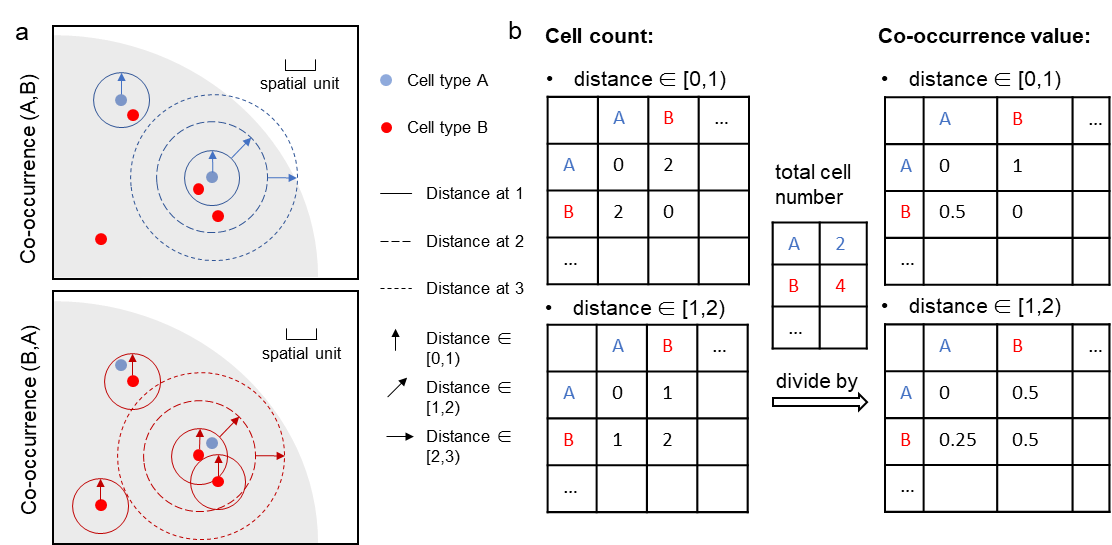

**Supplementary Fig. 3 Diagram of co-occurrence algorithm.**

**a,** A diagram of the co-occurrence of cell type A and B in SRT. The circle indicates a certain distance range to detect co-occurrence. **b,** The calculation for all cell types, left matrix represents the count of co-occurrence cells in a certain distance range. After being divided by the cell total count, it becomes the right matrix which represents co-occurrence result.

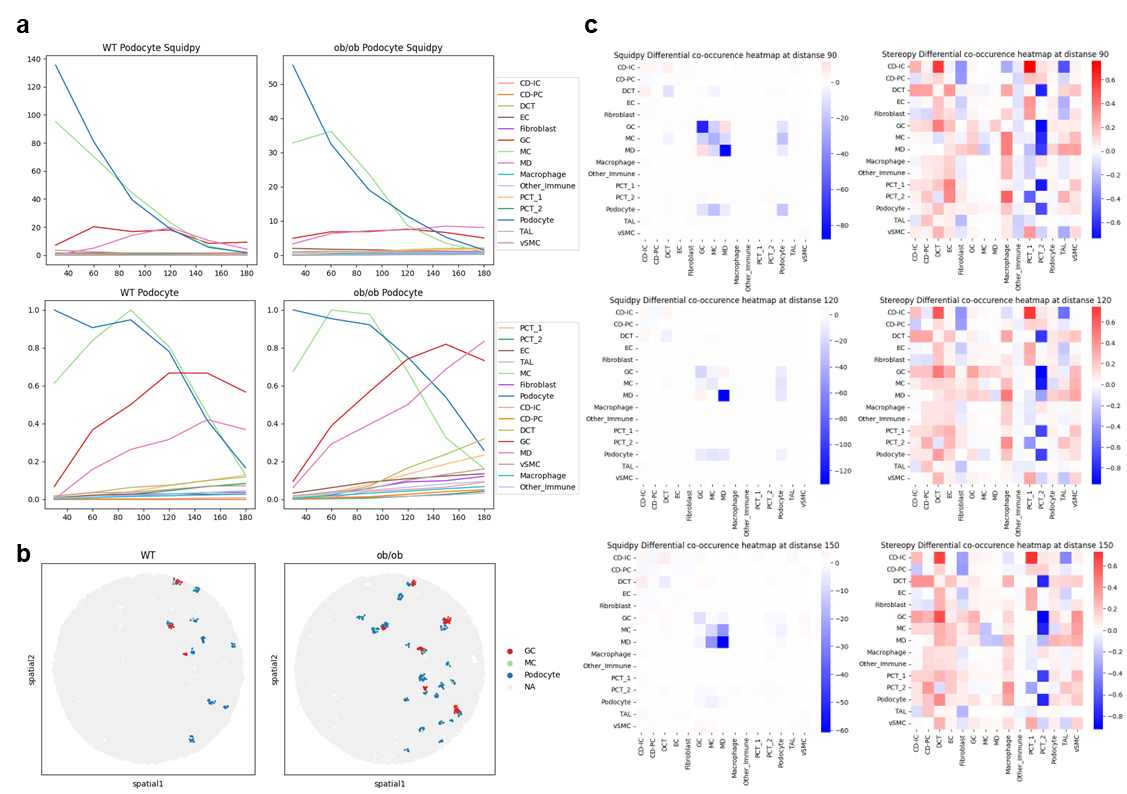

**Supplementary Fig. 4 Co-occurrence and DEG analysis on mouse kidney WT/BTBR samples of slide-seq V2.**

**a**, Line plot of podocytes’ co-occurrence with other cell types. Top: Squidpy’s method, bottom: Stereopy’s method; Left: WT sample, right: *ob/ob* samples. **b**, Spatial map of GC, podocyte and MC. **c**, Heatmap of cell co-occurrence with distance range from 90, 120 to 150. Left: Squidpy’s method, right: Stereopy’s method.

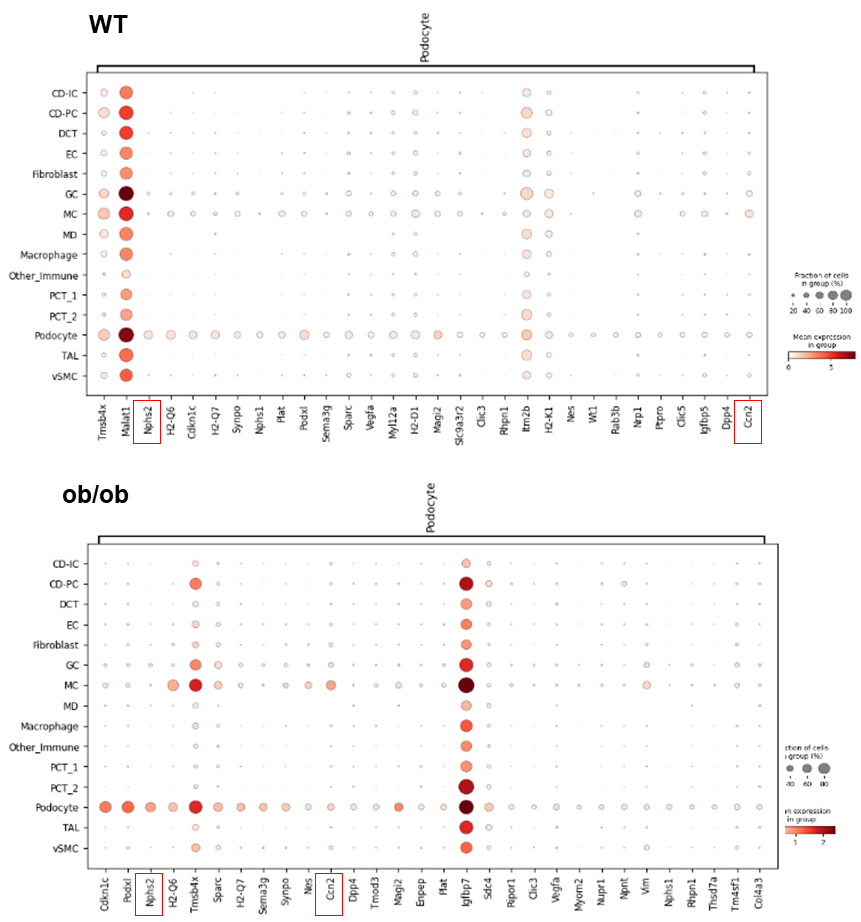

**Supplementary Fig. 5 Differential expression analysis for podocyte in WT and *ob/ob* samples respectively.**

The presence of two marker genes, *Nphs2* (a marker for podocyte) and *Ccn2* (a marker for injury podocyte), is visually indicated by two distinct red boxes in each plot. The rank of *Ccn2* has emerged within the top 10 in the injury sample.

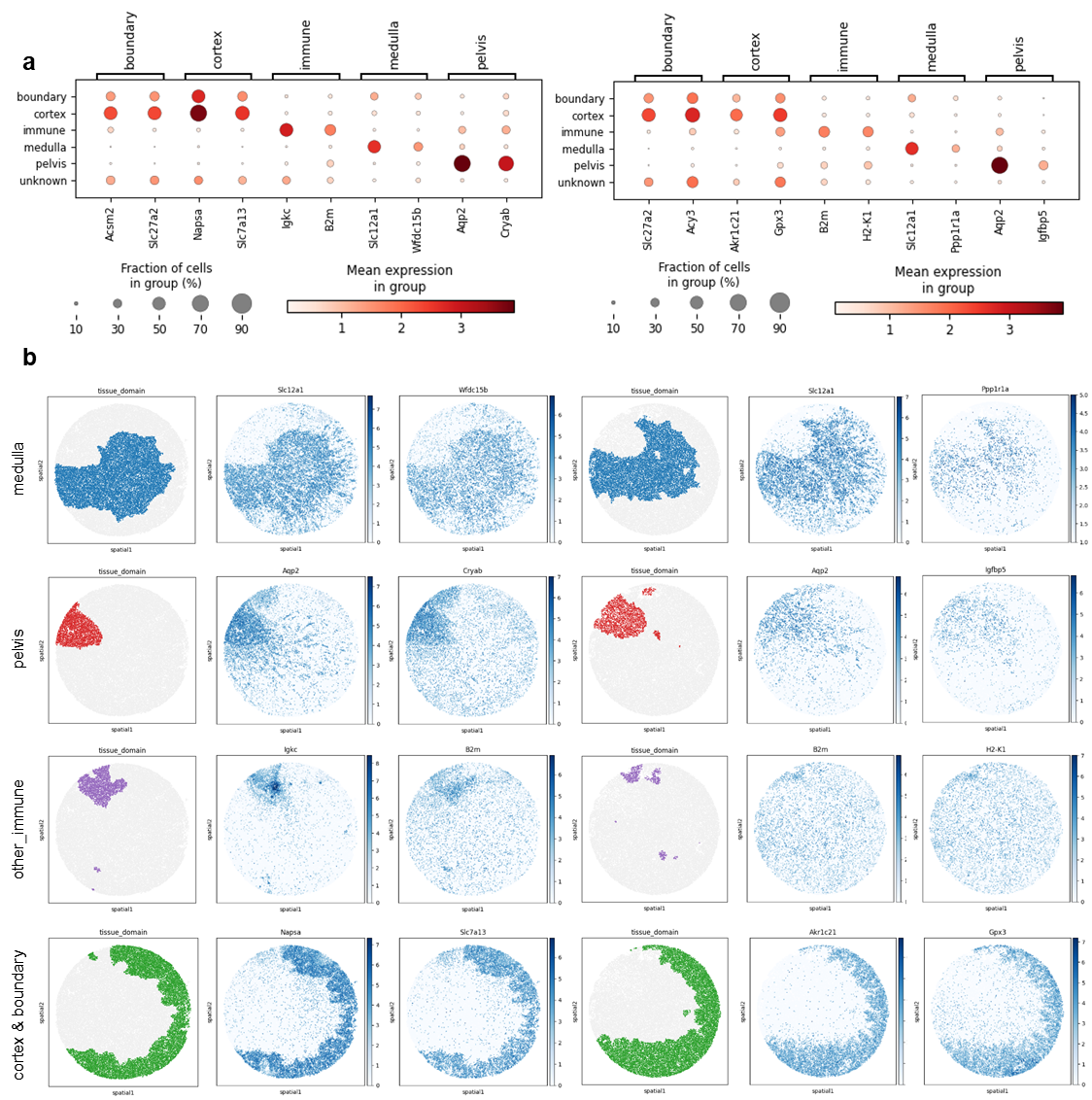

**Supplementary Fig. 6 Differential expressed genes of cell communities detected by Stereopy-CCD.**

**a,** Dotplot of differential expressed genes in cell communities detected by Stereopy-CCD. Left: WT mouse kidney sample, right: UMOD KI mouse kidney sample **b,** Spatial map of each cell community detected by Stereopy-CCD and corresponding differential expressed genes. Each row presents a cell community. Left: WT mouse kidney sample, right: UMOD KI mouse kidney sample.

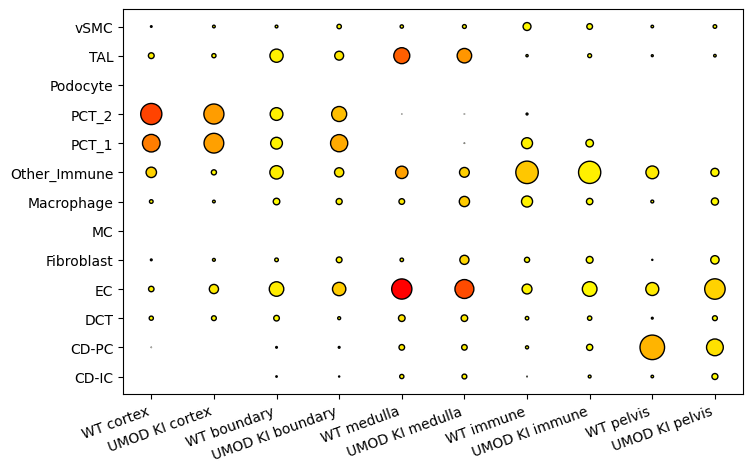

**Supplementary Fig. 7** **Cell type constitution in each cell community of both WT and UMOD KI mouse kidney sample.**

The dot size indicates ratio of cell type, color indicate cell amount of cell type.

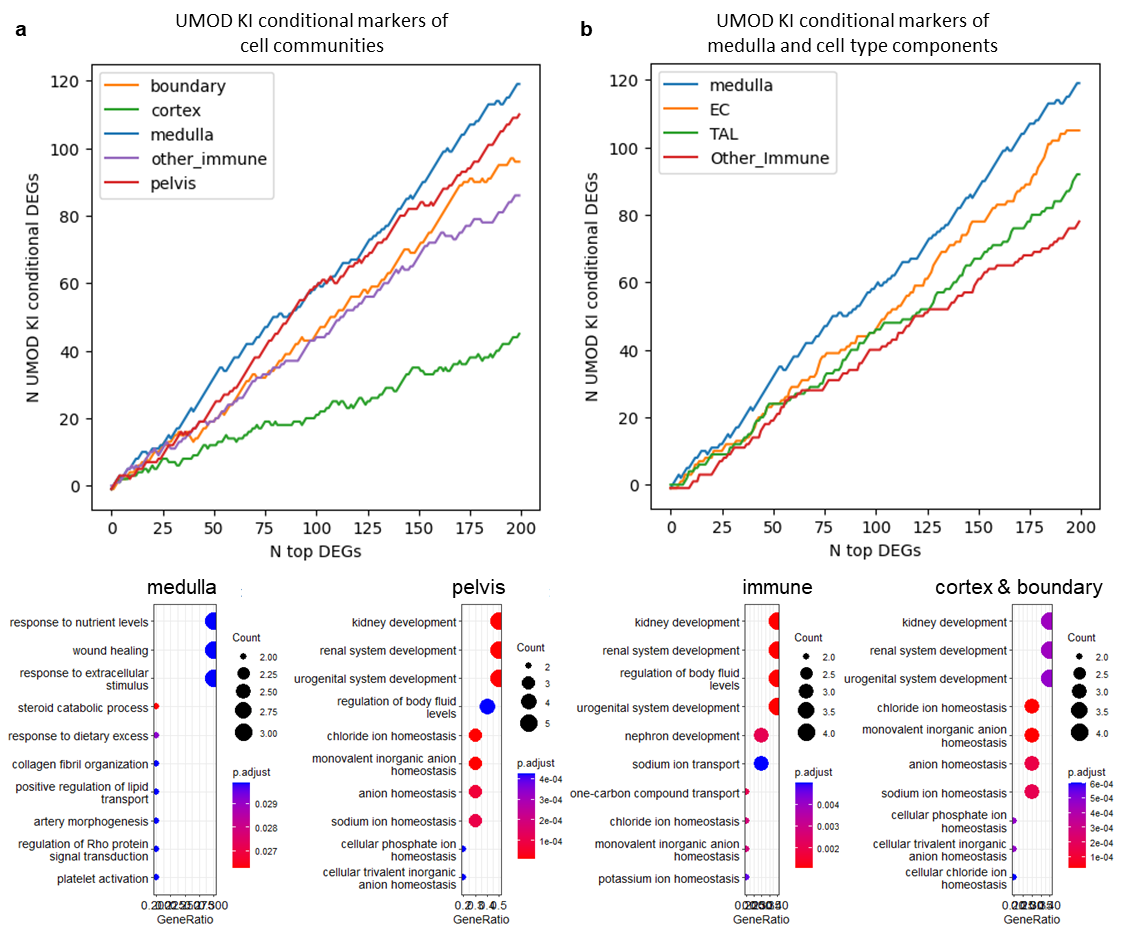

**Supplementary Fig. 8 UMOD KI conditional markers and enriched GO terms of medulla compared to other domains.**

**a,** Line plot of conditional marker amount for medulla and other tissue domains. X-axis represents *N* top DEG selected, Y-axis represent UMOD KI conditional markers amount. **b,** Line plot of constant marker amount for medulla and corresponding constitute cell types. X-axis represents N top DEG selected, Y-axis represent UMOD KI conditional markers amount. **c,** GO enrichment of UMOD KI conditional markers. From left to right present medulla, pelvis, immune, cortex & boundary, respectively.

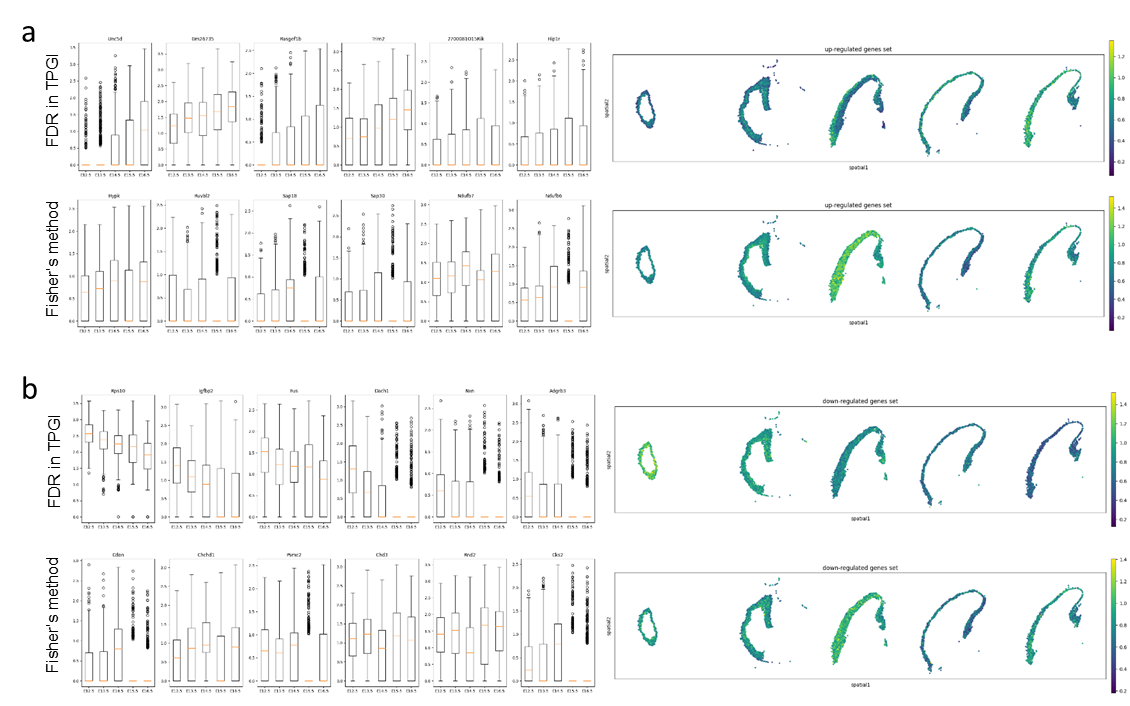

**Supplementary Fig. 9 Stereopy TPGI up and down regulated genes for temporal forebrain neuronal intermediate progenitor**.

**a,** Left: boxplot of up regulated genes based on different p value combination method; right: spatial map of gene set of mean expression of top 20 up regulated genes. The result is calculated based on pFDR in Stereopy-TPGI (Top), and fisher’s method (Bottom), respectively. **b,** Left: boxplot of down regulated genes based on different p value combination method; right: spatial map of gene set of mean expression of top 20 down regulated genes. The result is calculated based on pFDR in Stereopy-TPGI (Top), and fisher’s method (Bottom), respectively.

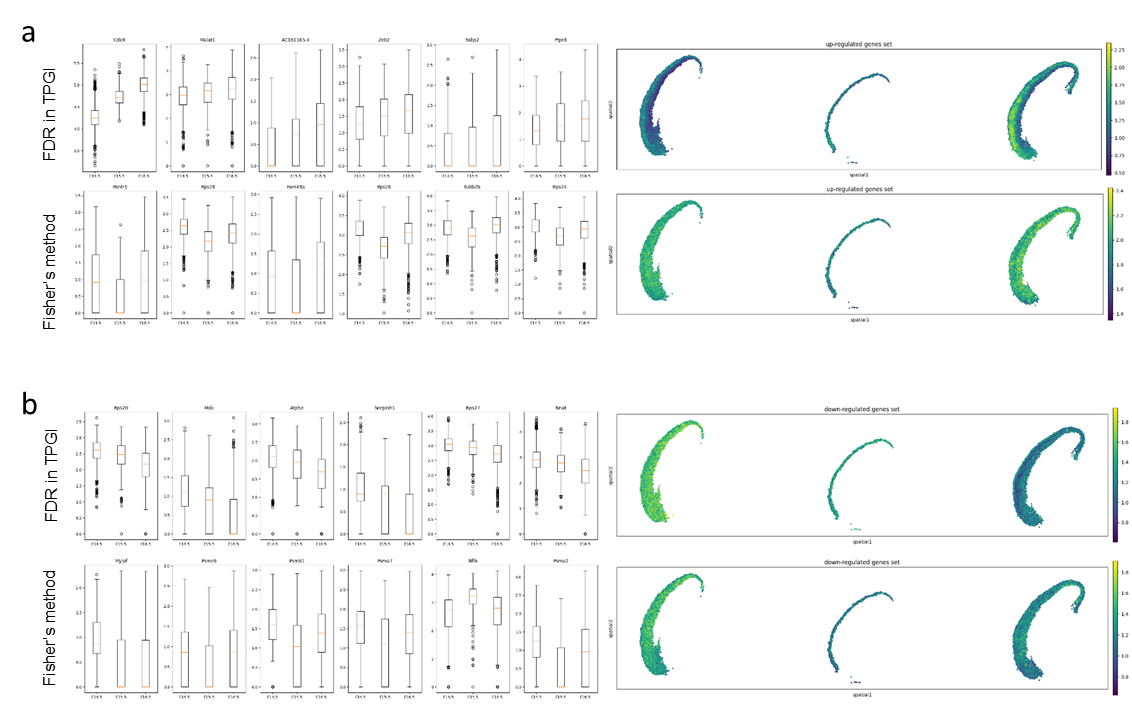

**Supplementary Fig. 10 Stereopy TPGI up and down regulated genes for temporal forebrain cortical glutamatergic**.

**a,** Left: boxplot of up regulated genes based on different p value combination method; right: spatial map of gene set of mean expression of top 20 up regulated genes. The result is calculated based on pFDR in Stereopy-TPGI (Top), and fisher’s method (Bottom), respectively. **b,** Left: boxplot of down regulated genes based on different p value combination method; right: spatial map of gene set of mean expression of top 20 down regulated genes. The result is calculated based on pFDR in Stereopy-TPGI (Top), and fisher’s method (Bottom), respectively.

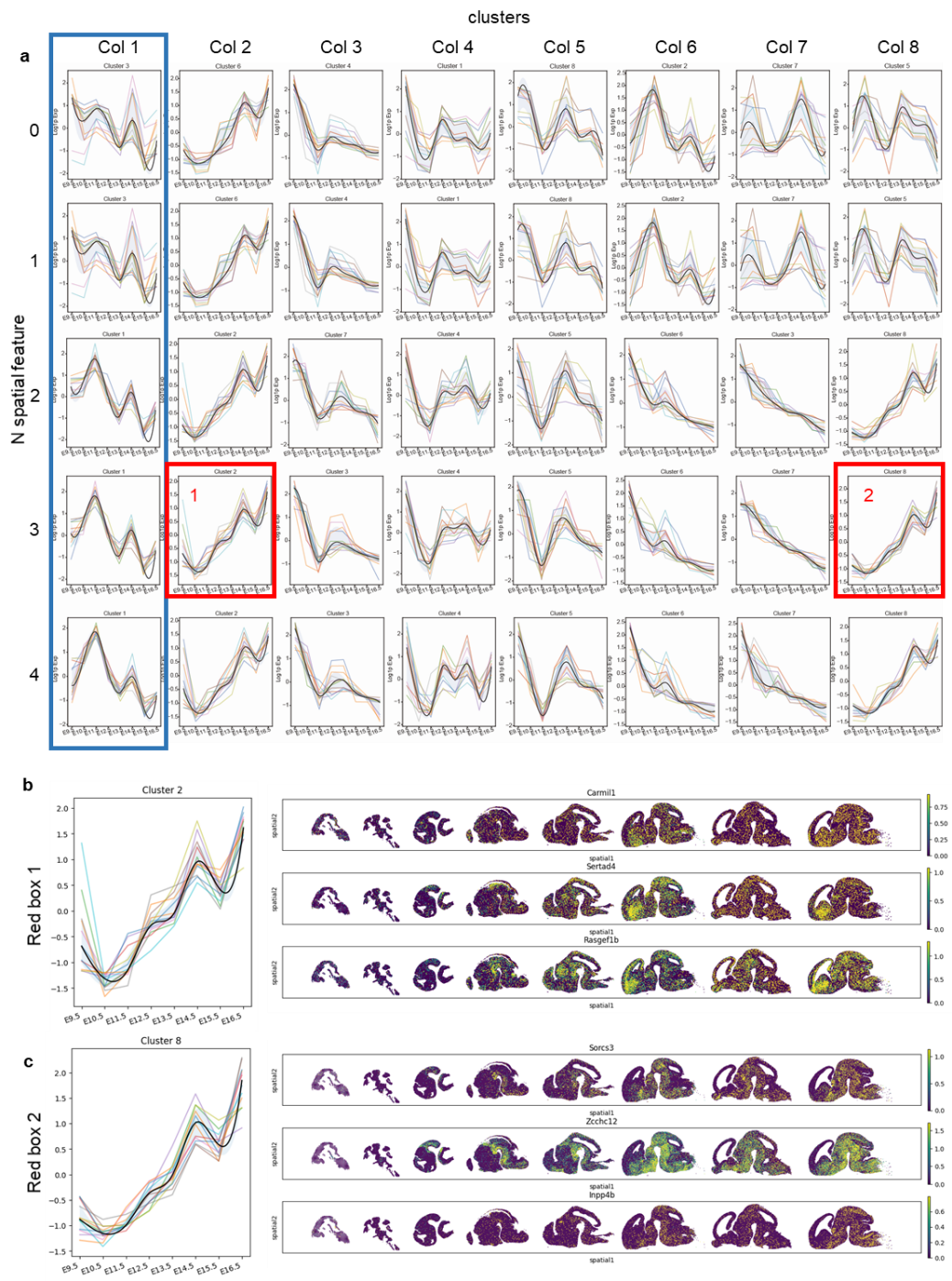

**Supplementary Fig. 11 The influence of spatial features on the detection of temporal gene patterns for temporal mouse brain.**

**a**, The temporal gene patterns calculated by Stereopy-TPGI with n spatial feature from 0 to 4. Take Col 1 (blue box) of gene patterns as example, the higher *n* spatial features result in higher consistent temporal expression of genes and temporal gene features in red boxes indicated spatial feature can distinguish genes with similar temporal pattern but existed in different regions. **b,** Left: Cluster 2 (red box 1) is amplified; Right: Spatial map of corresponding genes in this temporal gene cluster. **c,** Left: Cluster 8 (red box 2) is amplified; Right: Spatial map of corresponding genes in this temporal gene cluster. The clusters in two red boxes share the same temporal gene pattern but have different expression locations in temporal mouse brain datasets.

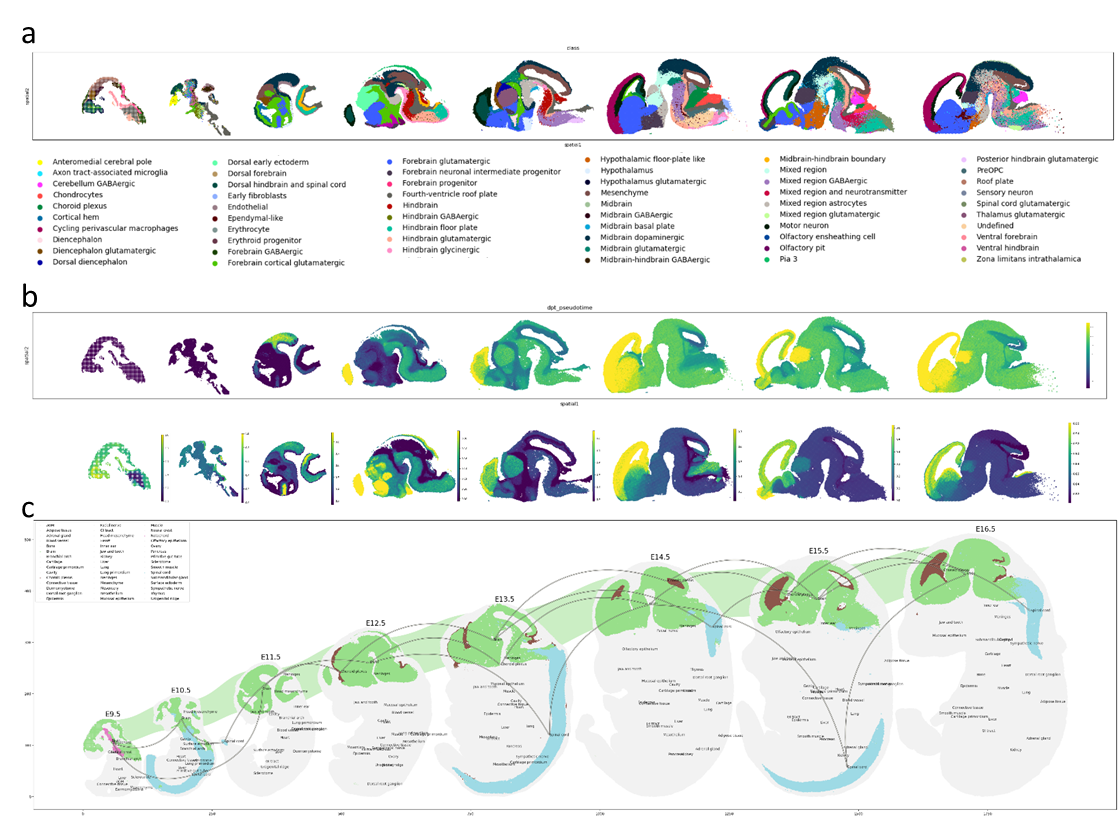

**Supplementary Fig. 12 Cell level diversity analysis among temporal mouse embryonic brain datasets.**

**a**, Manual annotation of cell types for mouse brain. **b,** Diffusion map of pseudotime for mouse brain calculated in integrated and parallel independent mode. **c,** Spatial trajectory plot with brain region emphasized.
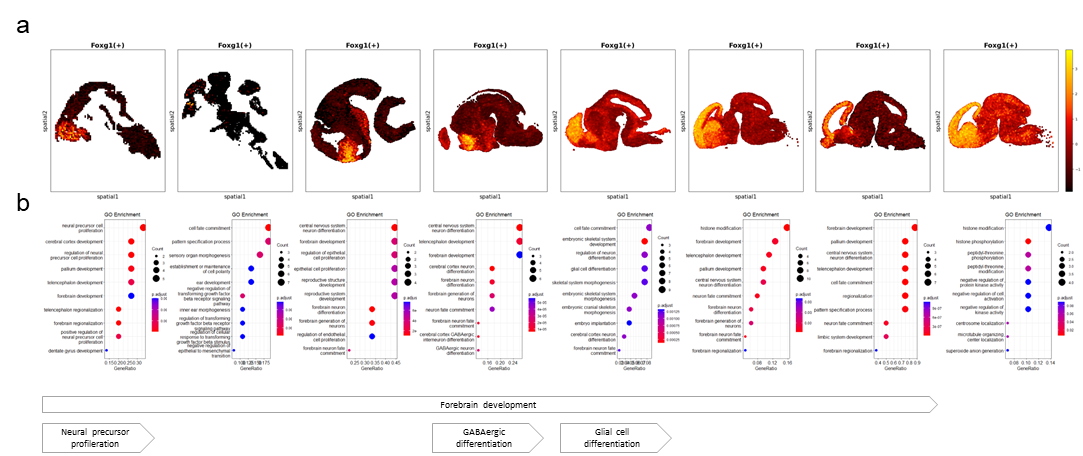

**Supplementary Fig. 13 The expression level of the regulon of *Foxg1* transcription factor and enriched GO terms calculated for the target genes regulated by *Fox1*.**

**a**, Spatial regulon plots for *Foxg1*(+) in eight time points of mouse brain. **b,** Corresponding GO enrichment for genes in regulons in eight time points of mouse brain.

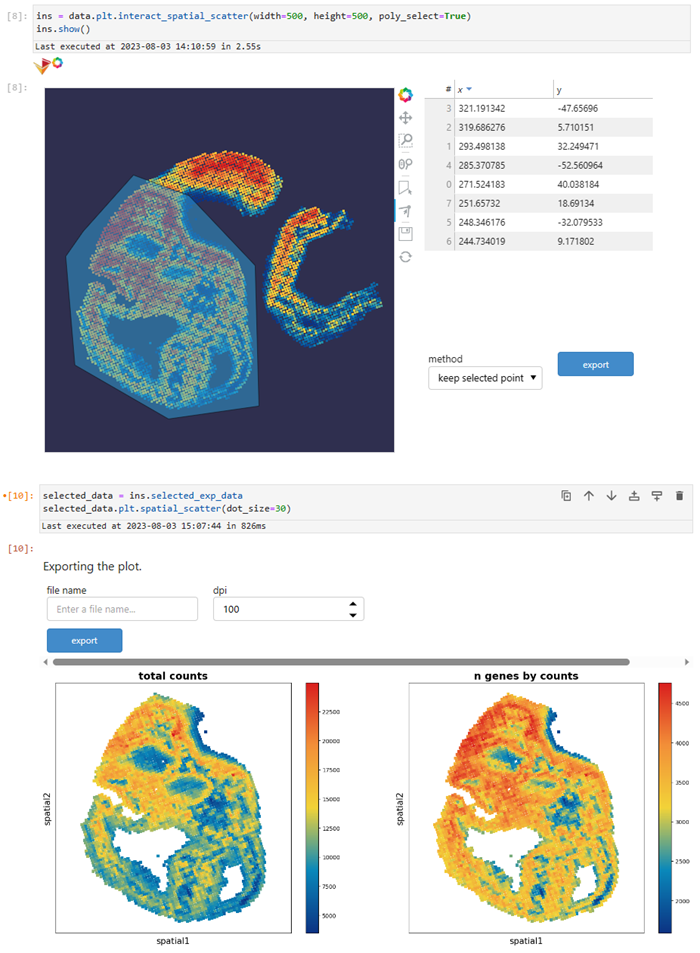

**Supplementary Fig. 14 Stereopy provides interactive analysis and visualization.**

Lasso function is applied on the mouse embryonic brain and the selected region is obtained.

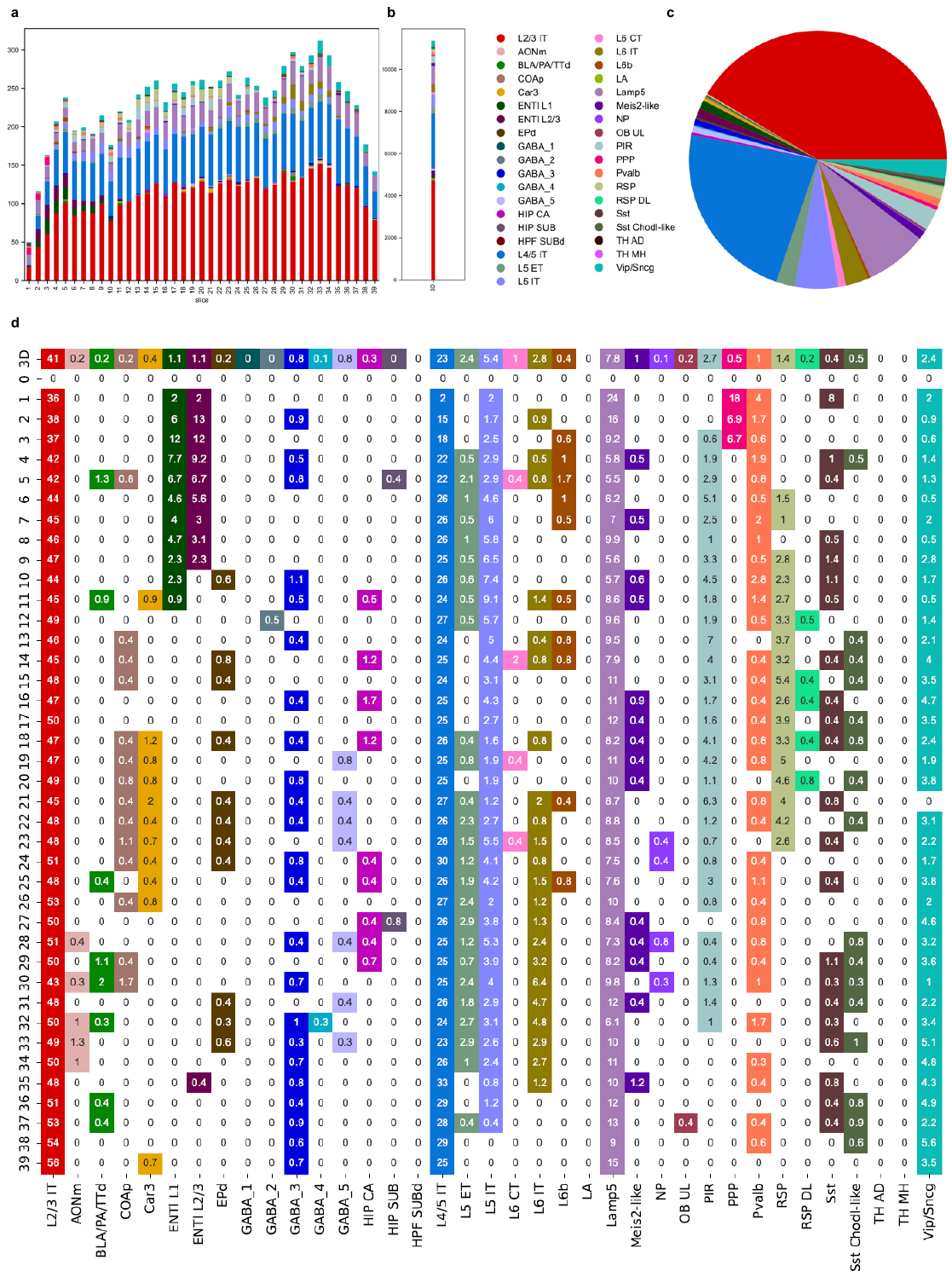

**Supplementary Fig. 15** **Comparison of 3D and 2D niche sizes and compositions on mouse brain BARseq data.**

**a**, Stacked chart showing cell type compositions of 2D niches constructed for each slice. **b, c,** Stacked chart and pie plot showing cell type compositions of the 3D niche. **d**, Cell type composition and percentages in 3D and individual 2D slices.

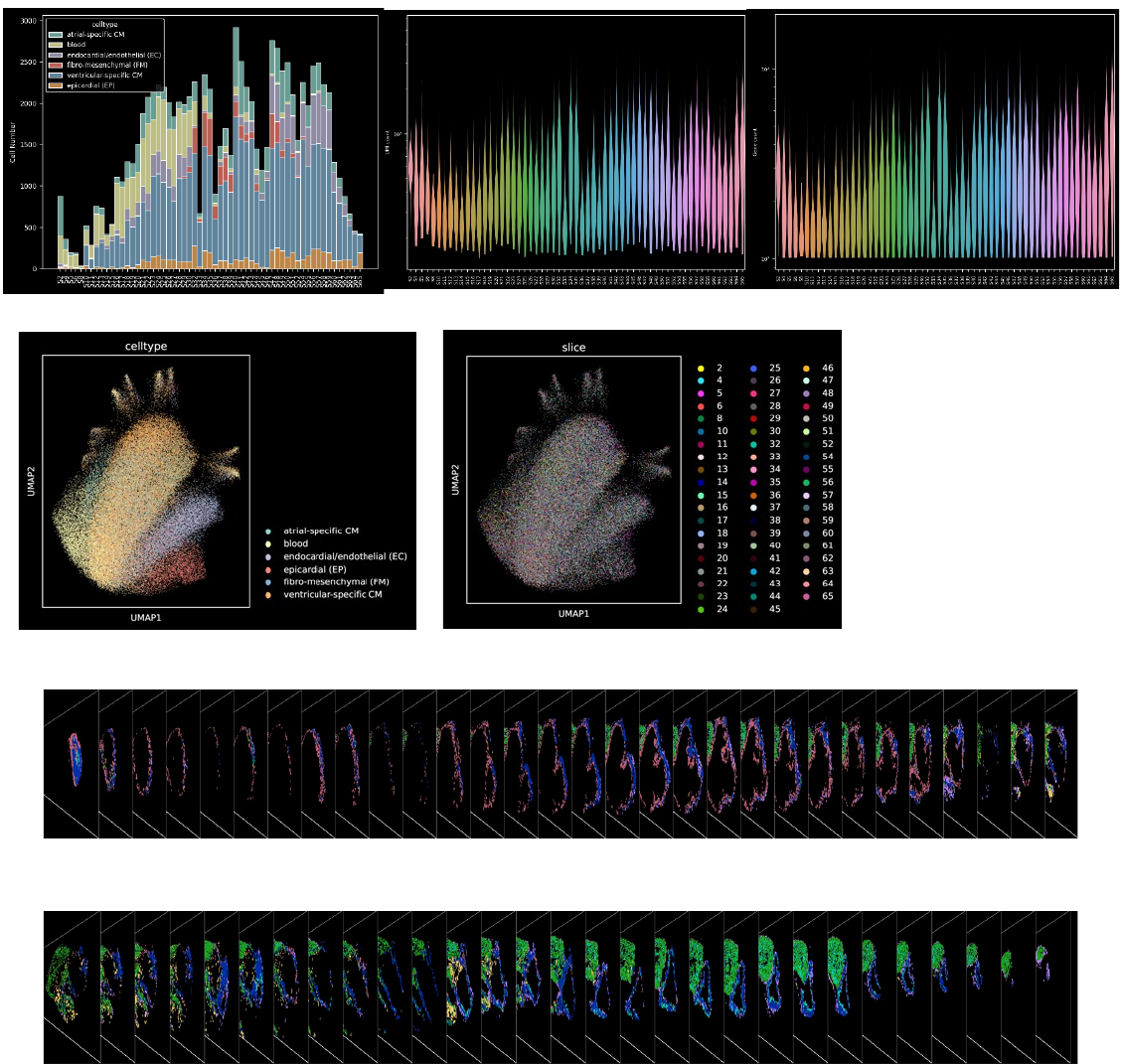

**Supplementary Fig. 16 Statistic and visualization results of 3D niche-based co-regulation.**

**a**, Stacked chart showing cell number distribution of six heart clusters in each of the 59 Stereo-seq sections. **b**, Violin plots of total UMI count and number of genes distribution in each of the 59 Stereo-seq sections. **c**, UMAP visualization of the six cardiac subclusters and 59 Stereo-seq sections. **d**, 2D visualization of the 59 Stereo-seq sections.

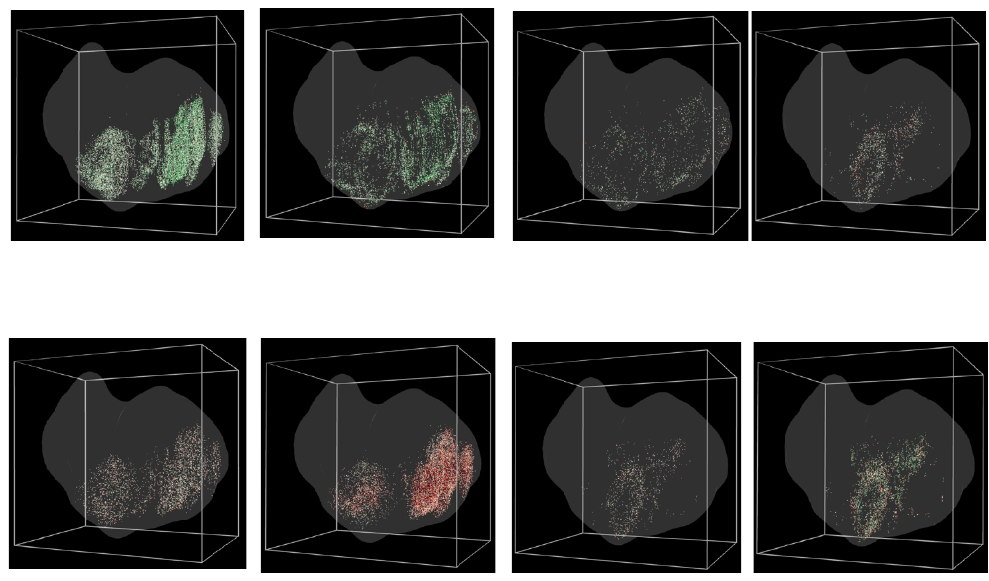

**Supplementary Fig. 17 3D visualization of selected cell-cell communications in four VCM niches.**

**a**, 3D visualization of four niche-specific L-R pairs from EC, EP, FM to VCM cells. **b**, 3D visualization of four niche-specific L-R pairs from VCM to EC and FM cells.

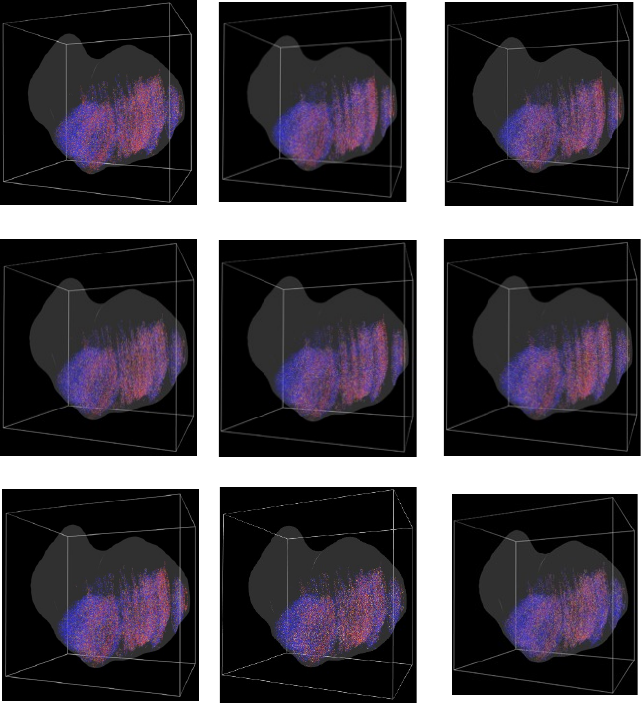

**Supplementary Fig. 18 3D visualization of selected regulons.**

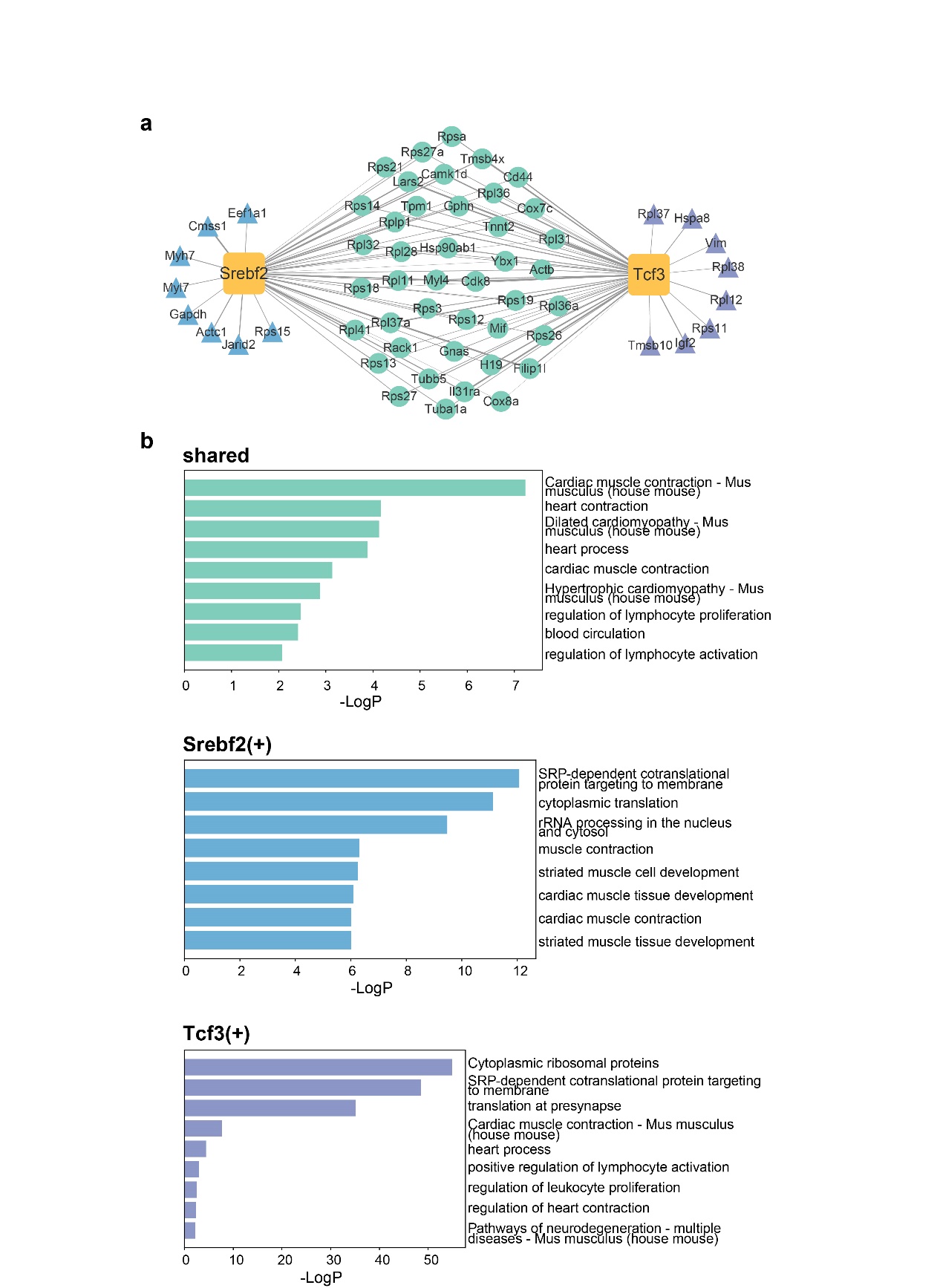

**Supplementary Fig. 19 Co-regulation function of *Srebf2*(+) and *Tcf*3(+).**

**a**, Shared and specific TGs in *Srebf2*(+) and *Tcf3*(+) regulons showing the 3D co-regulation function. **b**, GO enrichment analysis indicating the collective function of shared and specific TGs of regulons in (a) (shared, *Srebf2*(+)- and *Tcf3*(+)-specific from top to bottom).

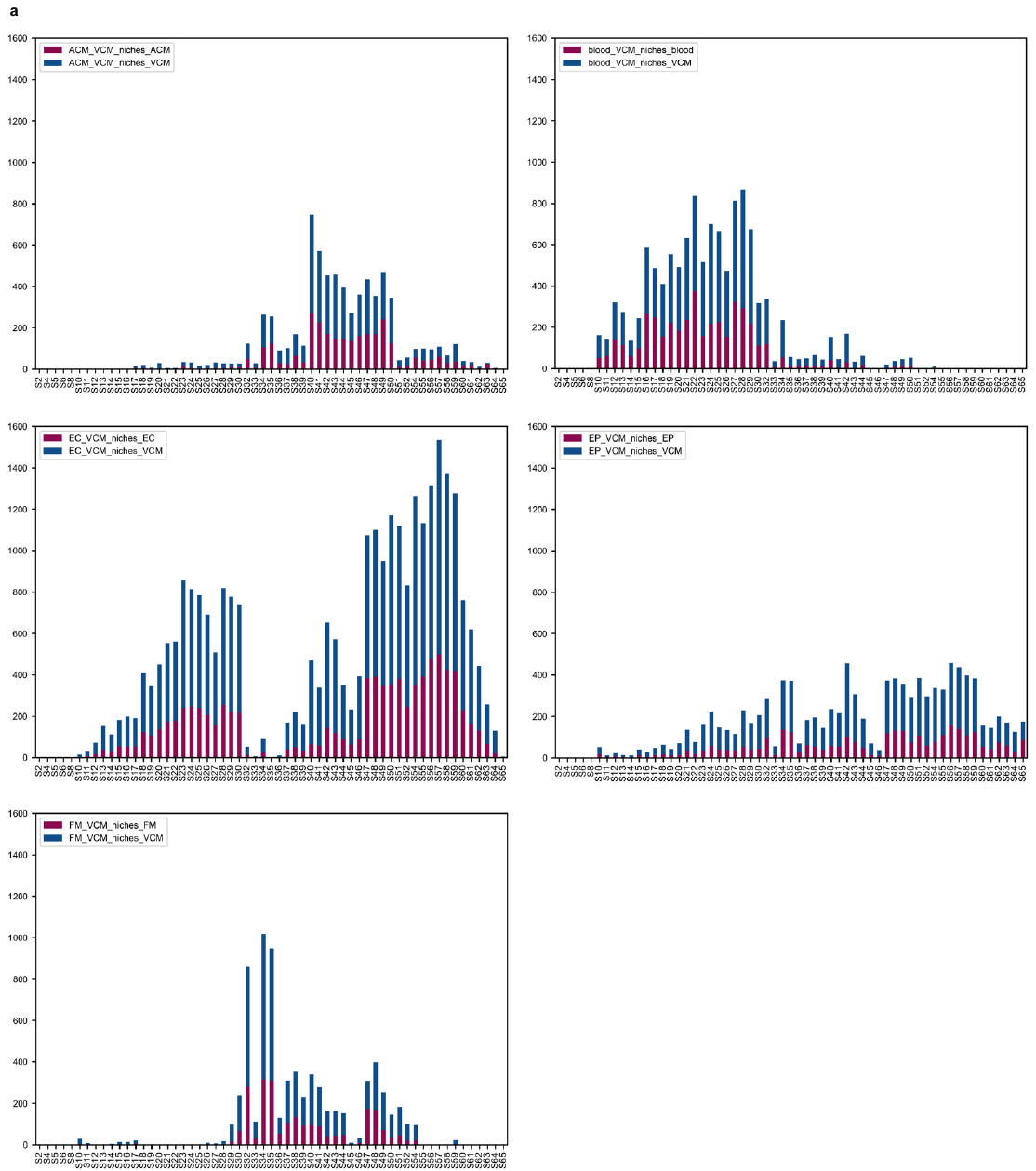

**Supplementary Fig. 20 Stacked charts of cell number distribution of five VCM niches among 59 Stereo-seq sections showing incomplete niche composition in each of the 2D samples.**

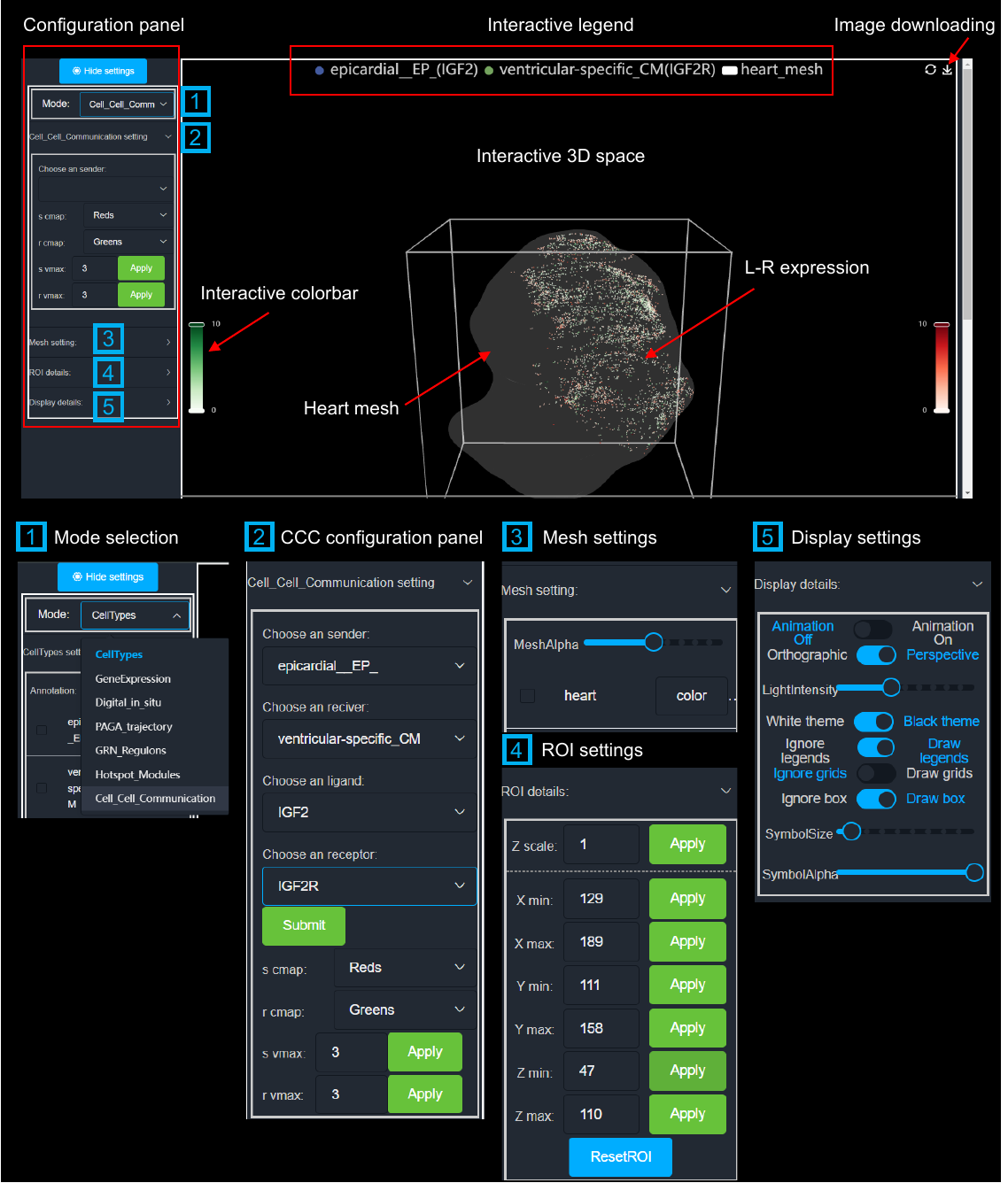

**Supplementary Fig. 21** 3D interactive visualization of L-R pairs rendered by VT3D.
