## Supplementary Note for "Stereopy: modeling comparative and spatiotemporal cellular heterogeneity via multi-sample spatial transcriptomics"

### 1. Overall analysis framework of Stereopy for multi-sample analysis

The analysis of multi-sample omics data encompasses a wide range of sample types, including replicated, differential, time-series, 3D, and various combinations thereof. The joint analysis of these multi-sample datasets plays a crucial role in uncovering key regulatory mechanisms in biology. However, most existing tools such as Squidpy [1], Giotto [2], Scanpy [3], and Seurat [4] are mainly designed for analyzing individual samples or integrating single samples. Thus, the functionality for multi-sample analysis is limited and lacks flexibility for comprehensive multi-sample analysis. Regrettably, the field of multi-sample analysis has not yet established a robust community due to the dearth of proper guidance and development. Consequently, progress in the advancement and application of multi-sample analysis methods has been sluggish, impeding the overall pace of scientific research. In stark contrast, the single-cell analysis community has made significant strides fueled by the continuous development of classical tools. Standard analysis modules, such as preprocessing, cell clustering, cell-cell interactions, gene regulation, and trajectory inference, have been readily adopted in scientific research, enabling the exploration of valuable information embedded within the data and accelerating scientific progress. To propel the development of multi-sample spatial data analysis, it is imperative to establish robust analysis tools, propose innovative analysis methods, and foster a thriving community of multi-sample analysis algorithms. These concerted efforts will not only bridge the existing gap but also facilitate comprehensive exploration of the complex interplay between spatial and multi-sample dimensions, ultimately driving scientific research forward.

Generally, the predominant approaches of multi-sample data analysis encompass comparative analysis, time-series analysis, and 3D integration analysis. To ensure the effectiveness of these analyses, it is imperative to establish appropriate data storage containers and analysis scheduling schemes that can cater to the personalized and diverse requirements of multi-sample datasets at their very core. These crucial aspects have been extensively expounded upon in a dedicated section of the main text. Furthermore, in our relentless pursuit to foster the growth of the multi-sample analysis community and genuinely facilitate the functionality of multi-sample analysis, we have meticulously planned and developed an array of comprehensive analysis content tailored specifically for these three essential analysis scenarios. To complement these advancements, we have also devised intuitive and user-friendly visualization tools, ensuring a seamless user experience.

The ultimate objective of these endeavors is twofold: firstly, to empower users to employ Stereopy for the fundamental analysis of multi-sample data, and secondly, to broaden the horizons of multi-sample data analysis through the introduction of novel and visionary features. By attaining these objectives, we aim to provide researchers with a powerful toolset that not only supports their immediate needs but also propels them towards groundbreaking discoveries in the realm of multi-sample data analysis.

#### 1.1 Comparative analysis

Comparative analysis is utilized to examine the diversities and similarities within data across various conditions, encompassing multiple hierarchical levels. Our objective is to provide researchers with comprehensive support in analyzing variations within the data, catering to their specific requirements for comparative sample analysis at both the cellular and genetic levels. To this end, Stereopy offers a range of analysis functions meticulously designed to explore cellular and genetic dynamics. Notably, we have developed an innovative co-occurrence detection algorithm and a robust cell community discovery method. These functionalities within Stereopy empower users to delve into important aspects such as cell co-occurrence, gene expression diversity, and specific gene expression patterns at the level of cell clusters and cell communities. By employing Stereopy, researchers can execute a myriad of diverse comparative analysis calculations, leveraging its comprehensive range of tools. The results of these analyses are diligently stored and thoughtfully visualized, facilitating thorough exploration and interpretation of the data.

#### 1.2 Temporal analysis

Temporal analysis typically focuses on studying the developmental trajectory and gene expression changes that unfold at different time points. Stereopy, our innovative platform, provides researchers with robust capabilities to ascertain cellular-level temporal trajectories and discern patterns in gene expression over time. To augment the detection of these gene expression patterns, we have devised novel algorithms that integrate both spatial and temporal information, thus enhancing the reliability of pattern detection. By harnessing the power of Stereopy, users gain the ability to infer cellular differentiation trajectories along the temporal axis and investigate corresponding alterations in gene expression patterns throughout the entire differentiation process. This comprehensive analytical framework facilitates the exploration of intricate regulatory mechanisms governing organisms as they undergo temporal changes, shedding light on the underlying dynamics of biological processes. By leveraging Stereopy's cutting-edge algorithms and integration of spatial and temporal data, researchers can uncover valuable insights into the intricate interplay between temporal dynamics and gene expression patterns. This empowers them to deepen their understanding of the regulatory processes driving cellular differentiation and ultimately advance our knowledge of organismal development.

#### 1.3 3D integrated analysis

The adoption of 3D joint analysis represents a significant advancement beyond the limitations of 2D data analysis, offering a transformative approach for holistic and comprehensive exploration of regulatory mechanisms in organisms. This analysis framework encompasses diverse regulatory processes, including cell interactions, gene regulations, and developmental trajectories. Stereopy, our comprehensive platform, presents a complete analysis workflow that seamlessly integrates data preprocessing, 3D alignment and reconstruction, cell-niche construction, inference of ligand (L)-receptor (R)-transcription factor (TF)-target (TG) pathways, and prediction of intracellular TF-centered regulatory networks. To facilitate accurate and convenient regulatory predictions within this 3D niche, Stereopy incorporates extensive and comprehensive databases. These databases serve as a valuable resource, enriching the analytical capabilities of Stereopy and enabling more precise regulatory predictions. Moreover, Stereopy offers additional functionalities, including 3D registration and 3D trajectory inference, further enhancing the analytical capabilities and predictive power of the platform. By harnessing the power of Stereopy, users can engage in predictive and analytical regulatory inferences within a highly accurate 3D niche, catering to diverse analytical needs. This transformative platform empowers researchers and scientists to unravel the intricacies of regulatory mechanisms with unprecedented depth and precision, thereby expanding our understanding of complex biological systems.

### 2. Algorithms improve analysis ability for multi-sample joint analysis scenarios

#### 2.1 Multi-sample cell community or domain detection (CCD) for Comparative analysis

**Question definition**

The comparative analysis between samples typically starts with cluster/domain annotation of cell types in the samples. This is followed by comparing and analyzing the differences in cells at the cluster/domain level. While there have been numerous published clustering annotation algorithms, researchers often experiment with various algorithms to achieve cell annotation during the research process. This helps to meet their research expectations more effectively. However, there is currently no suitable algorithm available to assist with the differential analysis between samples based on the cell annotation information that users have more confidence in.

In tissues, certain functional structures are often made up of collaborating cell types. These functional structures, composed of cells, play important biological roles. The cell communities that consist of these cells exhibit certain differences from the existing domain definitions. We define cell communities as areas that contain one or multiple cell types and have a relatively uniform distribution of these cell types. Due to the consistent distribution of cell types, cell communities also represent functionally consistent regions within localized tissues.

During the differential analysis between samples, analyzing the differences in cell communities or functional communities can provide support in exploring sample differences from a higher dimension. This, in turn, helps in discovering biological differences and related gene regulatory mechanisms. By searching for cell communities in multiple experimental and control samples, we can discover the following:

1) Cell communities that are unique to either the experimental or control samples. This enables in-depth analysis of the functional mechanisms of these cell communities.

2) Cell communities that are shared between the experimental and control samples. This reveals differences in gene expression and regulatory responses to different external perturbations.

By adopting this approach, we can broaden the scope of comparative data analysis at the functional region level.

**Existing solutions and their disadvantages**

1) Currently, the options for analyzing cell communities in multiple samples are quite limited.

2) There are only a few published algorithms available for annotating multiple samples together (eg. PRECAST [5] , BASS [6]), and some studies have conducted limited testing on some types of multi-sample data (eg. GraphST [7]). However, it is important to note that the performance of these algorithms greatly relies on their ability to effectively handle batch effects in multi-sample data. While these algorithms prioritize addressing batch effects, they tend to pay less attention to other important tasks, such as cell community or domain detection.

**PRECAST**: PRECAST is the unified and principled probabilistic model that can estimate low-dimensional embeddings and perform spatial clustering.

**BASS**: BASS employs Bayesian hierarchical modeling to perform cell type clustering at the single-cell scale and spatial domain detection at the tissue regional scale.

**GraphST:** GraphST is a graph self-supervised contrastive learning method that fully exploits spatial transcriptomics data for spatially informed clustering (spatial domain), batch integration, and cell-type deconvolution.

3) Single-sample clustering/domain detection algorithms tend to prioritize cell types rather than cell communities that demonstrate high similarity in gene expression, as these algorithms are developed based on this principle. For example, Giotto, SpaGCN [8], and GraphST [7] are some instances of such algorithms. However, these algorithms are unable to simultaneously identify cell communities across multiple samples. To achieve this, additional label matching is required.

**Giotto:** Giotto identifies spatial domains with coherent gene expression patterns by implementing their previously developed hidden Markov random field (HMRF) model. The authors argue that the detected spatial domains were consistent with anatomical layer structure on a SeqFISH+ dataset [2].

**SpaGCN:** SpaGCN performs spatial domain identification using aggregated gene expression data of neighboring spots obtained through its graph convolutional network approach. The authors claim higher coherency and integrity of domains and their biological interpretability compared to simple clustering methods [9].

**GraphST**: See previous section

**Our proposed cell community detection (SpaCCD) algorithm**

The Cell Community Detection (CDD) algorithm is designed to facilitate comparative analysis at the cell community level across different samples, utilizing cell type information (see Methods). It has been successfully applied to various spatial transcriptomics datasets. In order to assess its performance, we have compared CCD with other existing tools such as SpaGCN [8], GraphST [7] and Giotto [2], although there may be slight differences in target and input data as mentioned in the previous section. Stereopy-CCD outperforms Giotto, GraphST and SpaGCN in single-sample scenarios (e.g., whole mouse embryo brain) and outperforms SpaGCN and Giotto, GraphST, PRECAST and BASS in multi-sample scenarios (e.g., continuous adult mouse brain and mouse kidney) (Extended Data Fig. 2-4, Supplementary Table 1-2, and Methods). The results demonstrate that CCD is capable of effectively identifying cell communities or domains that align with existing knowledge. Moreover, CCD enables comparative analysis of multi-sample datasets at the cell community level and allows researchers to incorporate their own verified cell type information, providing flexibility. This approach expands the analytical framework for studying comparative spatial transcriptomic datasets.

#### 2.2 Spatial-resolved gene pattern identification (TGPI) for temporal analysis

**Question definition**

Temporal data analysis at the level of gene expression patterns can help uncover changes in important gene clusters during temporal development, thereby discovering gene clusters that are closely related at different temporal stages. Before the emergence of spatial transcriptomics techniques, the discovery of temporal variation patterns was done at the single-cell data level. The introduction of spatial information provides more features for exploring temporal variation patterns. Gene expression patterns that exist in both spatial and temporal dimensions are undoubtedly more reliable. Spatial features can promote the discovery of gene patterns that have both temporal expression consistency and spatial consistency, thus mining more reliable gene clusters that coexist in space and time. These genes are more likely to serve the same regulatory mechanism.

In addition to exploring temporal gene expression patterns in spatial dimensions, continuous changes in cell types can also uncover important gene clusters associated with cell changes. Here, we refer to gene patterns in the temporal dimension as “temporal gene patterns,” and gene patterns identified through cell trajectories or continuous changes in cell types as “trajectory gene patterns”. The most common gene pattern is the continuous up-regulation or down-regulation gene pattern, where these genes show a steady increasing or decreasing trend. Additionally, meaningful knowledge discovery can be obtained by combining other gene patterns with relevant prior knowledge.

**Existing solutions and their disadvantages**

1) Currently, there are limited algorithms available for detecting spatiotemporal gene patterns in multiple samples.

2) The existing algorithms for detecting temporal gene patterns include Mfuzz [10], which can be used for single-cell temporal gene pattern detection. Another algorithm called MEFISTO [11] can be applied to temporal data or spatial data to detect temporal and spatial factors instead of genes. Although the relationship between factors and genes can be analyzed, MEFISTO does not directly obtain the genes. Meanwhile, it is limited in its ability to work with spatiotemporal datasets and consider both spatial and temporal features simultaneously.

**Mfuzz:** Mfuzz is an R package that implements soft clustering tools. It is commonly used to identify gene patterns in various samples under different time points and conditions. [12-14].

**MEFISTO:** MEFISTO is a flexible and versatile toolbox, which incorporates the continuous covariate to identify temporal and spatial patterns of key factor rather than gene expression from multimodal data.

**Our proposed Spatial-resolved gene pattern identification algorithm**

Our proposed Stereopy-TGPI algorithm, which takes into account the consistency of gene expression in both temporal and spatial aspects (Extended Data Fig. 5), improves the accuracy of gene pattern identification (see Methods). We evaluated the significance of spatial features in Stereopy-TGPI and found that they play a crucial role in enhancing the consistency of temporal gene pattern detection (Extended Data Fig. 6). Compared with Mfuzz, Stereopy-TGPI’s identification was more correlated with real and pseudotime tendencies and capable of enriching significant GO terms relevant to neuron development in the time-series whole mouse brain (Extended Data Fig. 7 and Methods) [30]. The Stereopy-TGPI method provides an example of combining spatial and temporal features for subsequent exploration of multimodal characteristics.

#### 2.3 Niche-mediated regulation pathway prediction (NicheReg3D) for 3D integrated analysis

**Question definition**

The regulation of cellular behavior necessitates the intricate interplay of both intercellular and intracellular interactions, which, in turn, relies on the complete composition of the cellular niche. Accurate characterization of the niche is particularly crucial in a three-dimensional (3D) context, as it enables a comprehensive understanding of the regulatory mechanisms governing cellular behavior. In this study, our pipeline excels at capturing the precise composition of the 3D niche, allowing for the accurate assessment of both intercellular and intracellular regulations. By leveraging this capability, we can shed new light on the underlying regulatory mechanisms, providing valuable insights into cell fate determination, tissue development, homeostasis, and disease progression.

Understanding the importance of 3D regulation is of great interest to researchers and practitioners alike, especially concerning gene regulatory networks (GRNs) that can be governed by cell-cell communication (CCC). It has been observed that these GRNs often play pivotal roles in processes such as development or regeneration. Manipulating GRN expression through CCC pathways can exert significant effects on phenotype. However, conducting such analyses requires the extraction of accurate and precise 3D niche information, effectively masking irrelevant GRNs. This not only reduces the need for laborious and costly screening experiments but also enhances the efficiency and efficacy of studying these important regulatory processes.

Ultimately, the significance of studying regulation lies in its profound implications for determining cell fate and function within the spatial context of tissues. By comprehensively exploring the regulatory mechanisms at the single-cell level within the spatial context, we can elucidate the intricate processes underlying tissue development, homeostasis, and disease progression. This holistic understanding provides a solid foundation for advancing our knowledge in various fields of biology and medicine, paving the way for improved therapeutic strategies and diagnostic approaches.

**Existing solutions and their disadvantages**

At present, the availability of algorithms capable of comprehensively capturing the intricate spatial interactions within cellular niches in 3D multiple samples remains limited. While two-dimensional (2D) techniques offer valuable insights into cellular behavior, they fall short in capturing the complete nuances of cell-cell interactions along the z-axis.

The current landscape of GRN and CCC inference methods, including pySCENIC [15], iTALK[16], Nichenet [17], CellPhoneDB [18], NATMI [19], CellChat [20], CellCall [21], Tensor-cell2cell [22], and Connectome [23], primarily rely on single-cell transcriptomics data without considering cellular positions. These methods often predict cellular-specific GRN modules or ligand-receptor (L-R) pairs individually, thereby oversimplifying the intricate interplay of both intracellular and intercellular regulations within the spatial context.

Existing methods for deriving cell specific CCC from two-dimensional spatial transcriptomics (2D SRT) data, such as Giotto [2], CellPhoneDB 4.0 [18], and NICHES [24], tend to predefine possible pairs or cells involved in CCC inference. However, they overlook the critical importance of accurately extracting the 3D niche at the single-cell resolution, thus compromising the comprehensive understanding of cellular interactions within their spatial context.

**Our proposed niche-mediated regulation pathway prediction algorithm**

Our NicheReg3D algorithm has been meticulously designed based on the underlying hypothesis that the integration of CCC and GRN models holds the potential to enhance the accuracy of context-specific predictions concerning L-R-TF-TG interactions, particularly in relation to morphological phenotypical changes (see Methods). By employing our approach, we are able to achieve a comprehensive 3D reconstruction of 2D multiple samples, enabling the precise and accurate extraction of cellular niches at the single-cell resolution (Supplementary Fig.15). The analysis is then accomplished through the incorporation of both intercellular interactions and intracellular TF-based regulatory networks. Furthermore, to address the significant computational burden resulting from the large data size of SRT data, we have developed NicheReg3D based on a statistical-based algorithm for the prediction of cell cluster-specific CCC. In contrast to other tools connecting the outside and inside of the cells, such as NicheNet [17] and CellCall [21], Stereopy-NicheReg3D provided a more definitive and complete network for inferring how cell-niche-specific L-R pairs regulate intracellular regulon activities related to specific cellular functions. Compared to state-of-the-art CCC tools, including single-cell CellPhoneDB [18] and spatially resolved NICHES [24] (Supplementary Table 5), Stereopy achieved the most complete identification of specific L-R pairs that covered nearly all those derived by other tools, thanks to the precise niche extraction (Extended Data Fig. 9). This algorithm has been implemented using the Python programming language, leading to notable improvements in the speed and efficiency of CCC inference.

### 3. Stereopy analysis on multi-sample SRT datasets

#### 3.1 Analysis on Slide-seq mouse kidney *ob/ob* datasets

**Cell co-occurrence analysis.** We analyzed cell co-occurrence on mouse kidney BTBR datasets of slide-seq V2. Details of functions used and parameter settings can be found in section [Evaluation of the co-occurrence algorithm]. The differential co-occurrence is also implemented in Stereopy by function *ms_data.tl.integrate_cooccurrence(scope=’WT|BTBR’)*.

**Hotspot analysis.** The gene module within mouse kidney datasets was calculated by Hotspot by using *spatial_hotspot* function in Stereopy with parameters *use_highly_genes = False, use_raw=False and n_jobs=50*, WT and BTBR samples are calculated independently. Visualization of autocorrelation of certain parts of gene module is implemented by seaborn *clustermap.*

**Differential expressed gene analysis.** Differential expressed gene analysis is calculated by *find_marker_genes* function in Stereopy with default parameters based on the log-normalized expression matrix. WT and BTBR samples are calculated independently.

#### 3.2 Analysis on Slide-seq mouse kidney UMOD KI datasets

**Cell community detection.** Multi-sample multi-window processing was performed by Stereopy-CCD using *data.tl.ms_cellcommunity* with *window_sizes of 300 and 150*, and *sliding_steps of 150 and 50*, respectively. For community clustering the Leiden algorithm was selected with a *resolution = 0.2*.

**Differential expressed gene analysis.** DEGs are calculated on WT sample group by both tissue domain and user-defined cell type via *ms_data.tl.find_marker_genes* function in Stereopy based on log-normalized expression. Visualization of DEG result is implemented by *ms_data.plt.marker_genes_scatter* function with parameter *groups = ‘medulla’ and [‘EC’, ‘Other_Immune’, ‘TAL’]* for tissue domain and author defined cell type, respectively.

**Constant and conditional marker analysis.** To explore the difference expression considered both tissue domain and case or control condition, we firstly calculate the DEG of medulla in WT sample and UMOD KI sample independently. Then top 100 genes of t-test score are regarded as marker genes of medulla for WT and UMOD KI conditions. The intersection of genes under different conditions is assigned as a constant marker while the unique marker under certain conditional is assigned as the conditional marker. Constant marker is ordered by the average rank in DEG results; conditional marker is ordered by original rank in corresponding DEG result. Visualization is implemented by *ms_data.plt.marker_genes_heatmap* function in Stereopy.

**Gene ontology (GO) enrichment analysis.** We use clusterProfiler [25] to analyze GO enrichment for various types of genesets during analysis. Since all of our datasets are from mouse, biological process from the org.Mm.eg.db genome-wide annotation for the mouse was used as a GO database. Top 10 genes of each DEG or constant and conditional markers are used for GO enrichment.

#### 3.3 Analysis on Stereo-seq temporal mouse embryo samples

**Manually annotation on a subset of brain in mouse embryo datasets.** Thanks to the flexibility of Stereopy analysis on multiple samples, we can cluster and annotate each sample separately. First, we use Leiden to cluster each sample based on PCA embedding of each independent sample of the region with original annotation as the brain. The cluster number is a little higher than it should be to make data over-clustered. Then same cluster will be merged together during annotation. Then we compared our marker of each cluster with the markers according to developing mouse brain marker [26], and then chose the most similar group as the annotation of clusters. The data of mouse brain is from the brain region in Stereo-seq mouse embryo ST.

**PAGA and DPT pseudotime analysis.** Stereo-seq mouse embryo and brain We integrate PAGA and DPT pseudotime [27] into Stereopy to analyze trajectory inference and pseudotime estimation. We inferred trajectory of time series mouse embryo and mouse brain datasets by *data.tl.paga*. Both of the datasets are calculated based on PCA embedding (*data.tl.pca* with default parameters) cell neighborhood (*data.tl.neighbor* with default parameters) with highly variable genes (*data.tl.highly_variable_genes* with default parameters) after quality control (*data.tl.qc* with *n_genes_by_counts > 0.05*cell numbers*). With the help of time series, we can determine the direction of every edge, that the early emerged cell type should be the father nodes in a pair of cell types in an edge. Then, we mapped the result of PAGA tree in the spatial domain to visualize the trajectory for time series ST. The direct arrow is used to represent the edges in our direct PAGA tress. Every arrow from father-node cell type of preceding time point to child-node cell type of post time (Fig. 4b). The PAGA result of embryo is visualized by mapping the trajectory in spatial with samples position corresponding to time point by *ms_data.plt.paga_time_series_plot* while the PAGA graph is illustrated for mouse brain trajectory by *ms_data.plt.paga*. In addition, DPT pseudotime was calculated for mouse brain multi-sample data both integrated and separately to show the consistence of result by *ms_data.tl.dpt* with *mode = integrate* and *isolated*, respectively.

**Temporal gene pattern analysis.** In order to analyze the genes with similar gene patterns along time series or cell type trajectory, we analyzed temporal genes along time series and trajectory on mouse forebrain datasets. For temporal up or downregulated genes, we analyzed genes along cell type trajectory in forebrain from Forebrain progenitor, Cortical hem, Dorsal forebrain, Forebrain neuronal intermediate progenitor and Forebrain cortical glutamatergic using *ms_data.tl.time_series_analysis* with *run_method=’tvg_marker’* and *p_val_combination=’FDR’*. We further analyzed Stereopy-TGPI on cell type trajectory along forebrain cell type and along temporal forebrain after gene quality control utilizing *ms_data.tl.time_series_analysis* with *run_method=’other’*. The number of clusters *n_clusters* of Stereopy-TGPI on cell type trajectory and on temporal is set to 6 and 4 respectively.

**Temporal gene regulatory network (GRN) analysis.** Stereopy-NicheReg3D analyzed the temporal dynamic GRN. GRN is inferred based on the gene co-expression in spatial calculated by Hotspots. Each time point is calculated independently and all possible TFs are used for each sample. The visualization of AUCells score targets of a certain TF in spatial is also integrated in Stereopy.

**GO enrichment analysis.** GO enrichment analysis was performed on top 10 genes of each Stereopy-TGPI gene pattern which is ordered by weights of fuzzy C means cluster or combined p value in serial up and downregulated genes. All target genes are used as a gene set when analyzing the temporal GRN of TFs including Foxg1 and Tead1. Other parameters are the same as GO enrichment analysis on slide-seq V2 UMOD KI datasets of mouse kidney.

#### 3.4 Analysis on 3D E11.5 Stereo-seq mouse embryonic heart dataset

**3D alignment and reconstruction.** Stereopy-NicheReg3D starts with the 3D alignment and reconstruction. Each 2D mRNA heatmap was literately manually aligned based on the similarities of the morphology of its neighboring 2D samples in the same 3D x-y-z space. The space of each two neighboring samples along the z-axis was set to 20μm, the sampling separation of cryosections. Before 3D alignment, 2D mRNA heatmaps were masked by pairwise ssDNA-staining images if they were not located within the histological region. As a result, a new set of coordinates for each spot was calculated by the linear transformation matrices. The updated cell-gene expression matrices with aligned 3D x-y-z coordinates were converted into 3D triangular meshes by the Stereopy’s function elaunay_3d. Note that the batch effect among different 2D samples was neglectable, thus we did not use any removal algorithms (Supplementary Fig. 16c).

**Cell-niche communication prediction.** To investigate potential intercellular signals transduced between VCM and the other five types of cells, we constructed the niche for VCM by Stereopy-NicheReg3D. We first performed a DBSCAN clustering using the 3D coordinates of all the VCM cells in the mouse heart dataset to filter out rare cells distant from the ventricular area. Then we used the radius $r=0.025$ mm to determine the niche composition of VCM (*ms_data.tl.get_niche* with *niche_distance=0.025*). We performed a label permutation-based statistical CCC analysis based on the extracted VCM-niche dataset. We then evaluated the significance of the communication scores through random shuffling of the cell type labels of cells in the niche 1,000 times, and retained significant L-R pairs with a p-value smaller than 0.05 (*ms_data.tl.cell_cell_communication* with *analysis_type=’statistical’* and *database=’liana’*).

**L-R-TF-TG network construction.** We performed the analysis of spatially resolved GRN using the Stereopy-NicheReg3D’s multiple functions. In brief, the 3D spatially weighted single-cell expression data matrix is used to infer the gene-gene correlation network. The directional GRN modules involving the user-defined TFs are determined based on the pre-built database. The TF-TG networks are further purified using promoter and TF-binding information, which finally build regulons. The regulon activity is enriched using the AUCell algorithm to detect cell-identity-specific regulons (*ms_data.tl.regulatory_network_inference* with *auc_threshold=0.05* and *rank_threshold=9000*). We further connected these significant L-R interactions with these specific regulons, and thus chased the regulatory route from the niche signaling to TF-TG regulons inside VCM cells (*ms_data.tl.receptor_tf_path*).

**GO enrichment analysis.** GO pathway enrichment analyses of TFs and their TGs was performed using the Metascape online analysis platform [28], and the interaction network was rendered by Cytoscape [29].

**Advantages compared with 2D sample analysis.** Stereopy-NicheReg3D enables the 3D joint analysis that overcomes the data sparsity or bias induced by each single 2D sample [30] (Supplementary Fig. 20). As a result, Stereopy generates more comprehensive and accurate VCM-niche communications compared with any 2D sample (Extended Data Fig. 9).

**Flexible and interactive visualization.** Stereopy-NicheReg3D enables the visualization of possible signaling paths between L-R-TF-TG in VCM cells to analyze how they link together and influence the phenotypic changes, including L-R heatmap, L-R Circos plot, L-R bubble plot, L-R-TF Sankey plot, specific regulon enrichment heatmap, and influenced TF co-regulatory network. We also provided interactive browsing of VCM-niche L-R expression in the 3D space (Supplementary Fig. 21).
